# Transcriptional divergence of the zebrafish *sox17* lineage begins during gastrulation

**DOI:** 10.64898/2026.06.27.734843

**Authors:** Po-Shu Tu, John D. Thompson, Oscar A. Davalos, Gloria D. Ligunas, Nimra Khurram, Katrina K. Hoyer, C. Ben Lovely, Stephanie Woo, Stefan C. Materna

## Abstract

The endoderm is specified at the onset of gastrulation and subsequently undergoes extensive migration before forming an epithelial sheet that gives rise to multiple organs, including the gut and respiratory tracts. Although the gene regulatory network underlying endoderm specification and the later processes that regionalize the gut are increasingly well understood, comparatively little is known about the intervening developmental events. Using single cell transcriptomics, we profiled the zebrafish *sox17* lineage, comprising endoderm and dorsal forerunner cells, throughout and immediately after gastrulation. We found that dorsal forerunner cells remain transcriptionally homogeneous while undergoing coordinated temporal changes, associated with ciliogenesis and epithelial organization, during assembly of Kupffer’s vesicle. In contrast, endoderm cells transition from a migratory to an epithelial transcriptional state while progressively acquiring distinct regional identities. These findings indicate that endoderm regionalization emerges within the context of a broadly shared transcriptional program associated with migration and epithelialization.

## INTRODUCTION

The endoderm, one of the three germ layers, gives rise to multiple organs, including the gut and respiratory tract, urogenital system, thymus, and pharyngeal arches (Nowotschin et al., 2019; Weatherbee et al., 2026; Zorn and Wells, 2009). In vertebrates, endoderm and mesoderm are both induced by the TGF-β ligand Nodal, yet the two lineages rapidly diverge and adopt distinct migratory behaviors during gastrulation (Economou et al., 2022). In yolk-rich embryos such as teleosts and amniotes, endoderm cells ingress at the onset of gastrulation, undergo extensive dispersal, and subsequently converge toward the dorsal midline to form an epithelial sheet (Concha, 2025). In contrast, mesoderm cells migrate collectively, maintaining cell–cell contacts, while undergoing coordinated convergence and extension movements (Pinheiro and Heisenberg, 2020; Williams and Solnica-Krezel, 2020).

In zebrafish, endoderm morphogenesis is distinguished by particularly dynamic and prolonged cell dispersal during gastrulation. During early gastrulation, endoderm cells migrate in a random walk-like manner and actively avoid one another through contact inhibition of locomotion (CIL) (LaBelle et al., 2025; Pézeron et al., 2008; Woo et al., 2012). By mid-gastrulation (80% epiboly), endoderm cells are evenly distributed from the margin up to, but excluded from, the animal domain (LaBelle et al., 2025; Nair and Schilling, 2008; Tavano et al., 2025)(Fig. 1A). Around this time, cell movements become increasingly directional, with cells migrating toward the dorsal midline (Mizoguchi et al., 2008; Nair and Schilling, 2008; Tu et al., 2026); simultaneously, contact duration between cells increases (LaBelle et al., 2025), leading to the formation of a simple epithelial sheet shortly after gastrulation. Over the following two hours, remaining endoderm cells are integrated laterally and gaps within the sheet are closed. By early somite stages, this epithelium extends from the anterior region, where the mouth will form, to the tailbud, where it surrounds Kupffer’s vesicle (KV). By 24 hours post-fertilization (hpf), the endoderm becomes regionally organized into a broad anterior domain flanked by the pharyngeal pouches and a narrower trunk endoderm that extends posteriorly to the cloaca (Ng et al., 2005).

**Figure 1.**
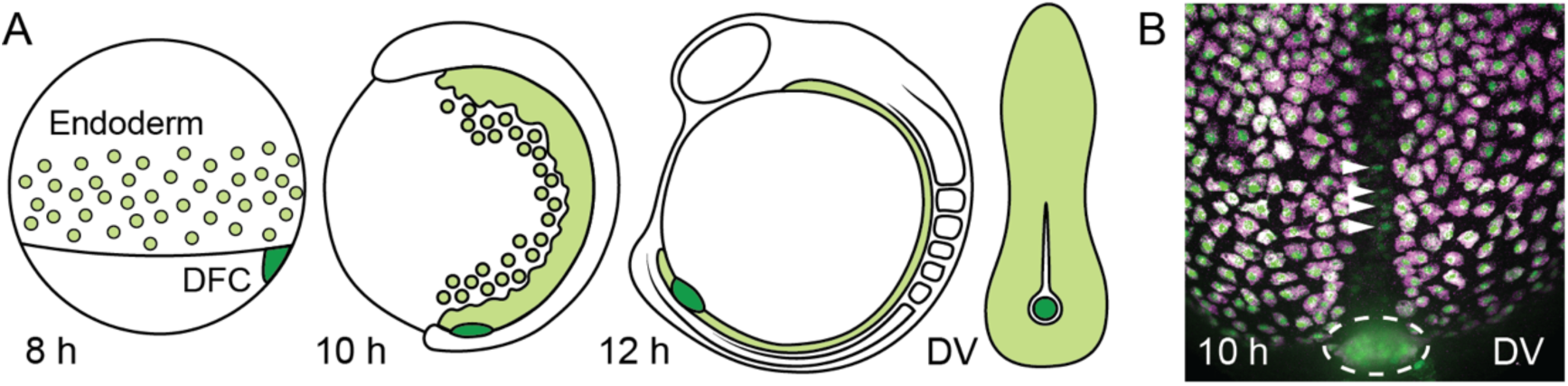
The *sox17* lineage from mid-gastrula to early somitogenesis. A) Schematic of endoderm and DFC morphogenesis. Following ingression, endoderm cells (light green) disperse during gastrulation, while DFCs (dark green) migrate as a group ahead of the margin (8 h). At bud stage (10 h) endoderm cells have started to converge on the midline where they form an epithelial sheet (12 h). Post-gastrulation, DFCs form Kupffer’s vesicle (KV), the organ of asymmetry, which is surrounded by endoderm. 8 h, 10 h, and 12 h (left) are lateral views, dorsal to the right. 12 h (right) is a dorsal view, anterior towards the top. B) In *Tg(sox17:GFP)^S870^* embryos, GFP expression (green) overlaps with endogenous *sox17* transcripts (magenta) in endoderm and DFCs (dashed circle). Notochord cells (arrowheads) express GFP but not endogenous *sox17*. DV–dorsal view, anterior towards the top.

Despite extensive characterization of endoderm morphogenesis, comparatively little is known about how transcriptional states change as endoderm cells transition from dispersed migratory cells to a polarized epithelium. More broadly, it remains unclear when transcriptionally distinct populations emerge within the endoderm and how regional patterning is coordinated with the global morphogenetic movements of this tissue. Recent advances in single-cell RNA sequencing have transformed our understanding of cell-state diversity in the gastrula-stage zebrafish embryo (Aponte-Santiago et al., 2026; Farrell et al., 2018; Lange et al., 2024; Wagner et al., 2018). However, endoderm comprises only approximately two percent of embryonic cells, limiting its representation and resolution in whole-embryo datasets (Economou et al., 2022).

Endoderm development is tightly linked to expression of *sox17* across vertebrates, including the specification of definitive endoderm in mammals (Johal et al., 2025; Viotti et al., 2014; Weatherbee et al., 2026). In zebrafish, a lineage-specific duplication gave rise to two genes, *sox32* and *sox17* (Kikuchi et al., 2001; Viotti et al., 2014). Although only *sox32* is required for endoderm specification, both genes exhibit nearly identical expression patterns during early development. Expression is first detected in the yolk syncytial layer (YSL) before being activated in endodermal cells and dorsal forerunner cells (DFCs), the precursors to the left-right organizer KV. Like endoderm, DFCs are specified downstream of Nodal signaling and undergo stereotypical morphogenetic movements during gastrulation (Feldman et al., 1998; Forrest et al., 2022; Kikuchi et al., 2001). However, unlike the highly dispersive migration of endodermal cells, DFCs migrate collectively as a cohesive cluster ahead of the blastoderm margin and subsequently form a simple epithelium during KV morphogenesis (Essner et al., 2005; Matsui et al., 2015a; Oteíza et al., 2008; Pulgar et al., 2021). Concurrent with lumen formation, KV cells assemble motile cilia that are required for left-right symmetry breaking (Aljiboury et al., 2022; Forrest et al., 2022). Thus, the YSL, endoderm, and DFCs together constitute the embryonic *sox17* lineage.

Here, we define transcriptional states of the *sox17* lineage from mid-gastrulation to early somitogenesis. Using single-nuclei RNA sequencing, we comprehensively sampled endoderm and the small population of DFCs. Consistent with their shared developmental origin, endoderm and DFCs exhibited substantial overlap in gene expression early in gastrulation. However, their transcriptional profiles became increasingly distinct as gastrulation progressed, paralleling their divergent cellular behaviors. DFCs remained comparatively homogeneous throughout gastrulation but underwent a pronounced transcriptional transition during KV formation.

In contrast, endodermal cells exhibited substantial transcriptional heterogeneity. Across the endoderm, gene expression progressively shifted toward programs associated with epithelialization as cells converge toward the dorsal midline; however, many endoderm-enriched genes were restricted to distinct cellular subsets, which became more pronounced over time, indicating substantial heterogeneity within this germ layer. Using hybridization chain reaction RNA-fluorescence in situ hybridization (HCR RNA-FISH), we mapped these transcriptional states to discrete and complementary spatial domains within the embryo. Notably, evidence of regional diversification was already apparent by mid-gastrulation, when cells near the margin activate genes associated with posterior endodermal identities. These findings indicate that endoderm patterning begins earlier than previously appreciated and proceeds concurrently with the extensive morphogenetic movements that shape the embryonic endoderm.

## RESULTS

### Comprehensive sampling of the *sox17* lineage across gastrulation

To enrich for endodermal cells and improve transcriptional resolution relative to existing whole-embryo datasets, we used the *Tg(sox17:GFP)^S870^* reporter line, which is widely used in studies of endoderm and DFCs during early zebrafish development (Chung and Stainier, 2008; Mizoguchi et al., 2008; Sakaguchi et al., 2006). To confirm that GFP reporter expression faithfully recapitulates endogenous *sox17* expression, we performed whole-mount HCR RNA-FISH for endogenous *sox17* transcripts in *Tg(sox17:GFP)^S870^* embryos (Fig. 1B). We observed complete concordance between *sox17* transcript and GFP expression in the endoderm, DFCs, and the YSL. However, the GFP reporter was also expressed in notochord cells, where endogenous *sox17* transcripts were not detected (Fig. 1B), and this ectopic expression became more pronounced after gastrulation. Despite this additional notochord expression, *Tg(sox17:GFP)^S870^* faithfully captures endogenous *sox17* expression during the developmental stages analyzed here and enables robust enrichment of endoderm and DFC populations.

To obtain *sox17:GFP*-positive cells, we dissociated *Tg(sox17:GFP)^S870^* embryos and enriched labeled cells by fluorescence-activated cell sorting (FACS). Cells in the *sox17* lineage comprise approximately 2% of all embryonic cells (Economou et al., 2022). To balance cell yield and viability with sorting duration, we targeted approximately 80% purity.

To characterize gene expression changes associated with endoderm and DFC morphogenesis during and immediately following gastrulation, we sampled cells at three developmental stages: 75–80% epiboly, bud stage, and 5-somite stage, corresponding to approximately 8, 10, and 12 hpf, respectively (Fig. 1A). These time points span multiple morphogenetic transitions — dispersal, directional migration, and epithelial differentiation for the endoderm and migration, KV assembly, and lumen formation for the DFCs.

### snRNAseq of the *sox17* lineage

To capture the current transcriptional state of cells, we isolated nuclei following FACS-sorting (Lake et al., 2016). We generated 3′-capture single nucleus RNA sequencing (snRNAseq) libraries using the 10X Genomics platform and obtained approximately 2×10^8^ sequencing reads per sample. Using the Cell Ranger software, we performed an initial analysis to filter out doublets and low-quality nuclei. For all three timepoints combined, we obtained a total of 11617 nuclei with an average of 1513 identified genes per cell. Because nuclear transcriptomes closely reflect cellular identity (Habib et al., 2017), we refer to recovered nuclei as cells throughout the remainder of the manuscript.

An important consideration when profiling early zebrafish embryos is the potential contribution of maternally deposited transcripts. Because nuclear isolation largely excludes mature cytoplasmic mRNAs, snRNA-seq preferentially captures nascent and nuclear-retained transcripts and therefore provides a more direct measure of ongoing transcriptional activity. To determine whether maternal RNAs might nonetheless influence our analysis, we quantified the abundance of the maternally expressed transcription factors *otx1* and *eomes* (Bjornson et al., 2005; Paraiso et al., 2019). Both transcripts declined to undetectable levels prior to our earliest sampling point at 8 hpf (Fig. S1). These findings indicate that maternally deposited transcripts are unlikely to persist at the developmental stages examined here and suggest that the recovered nuclear transcriptomes primarily reflect active zygotic gene expression programs.

We next performed clustering analysis of the integrated dataset containing all three samples and reduced dimensionality for visualization using UMAP (Becht et al., 2018) (Fig. 2A). We assigned cluster identities based on expression of genes with well-characterized spatial expression patterns. At mid-gastrulation, *met* is highly expressed in endoderm (Latimer and Jessen, 2008; Tu et al., 2026; Wan et al., 2026); *cftr* is restricted to DFCs (Navis et al., 2013); and *twist2* is a known marker of mesoderm/notochord (Germanguz et al., 2007). Each marker localized to a distinct major cluster, corresponding to endoderm, DFCs, and notochord (Fig. 2B).

**Figure 2.**
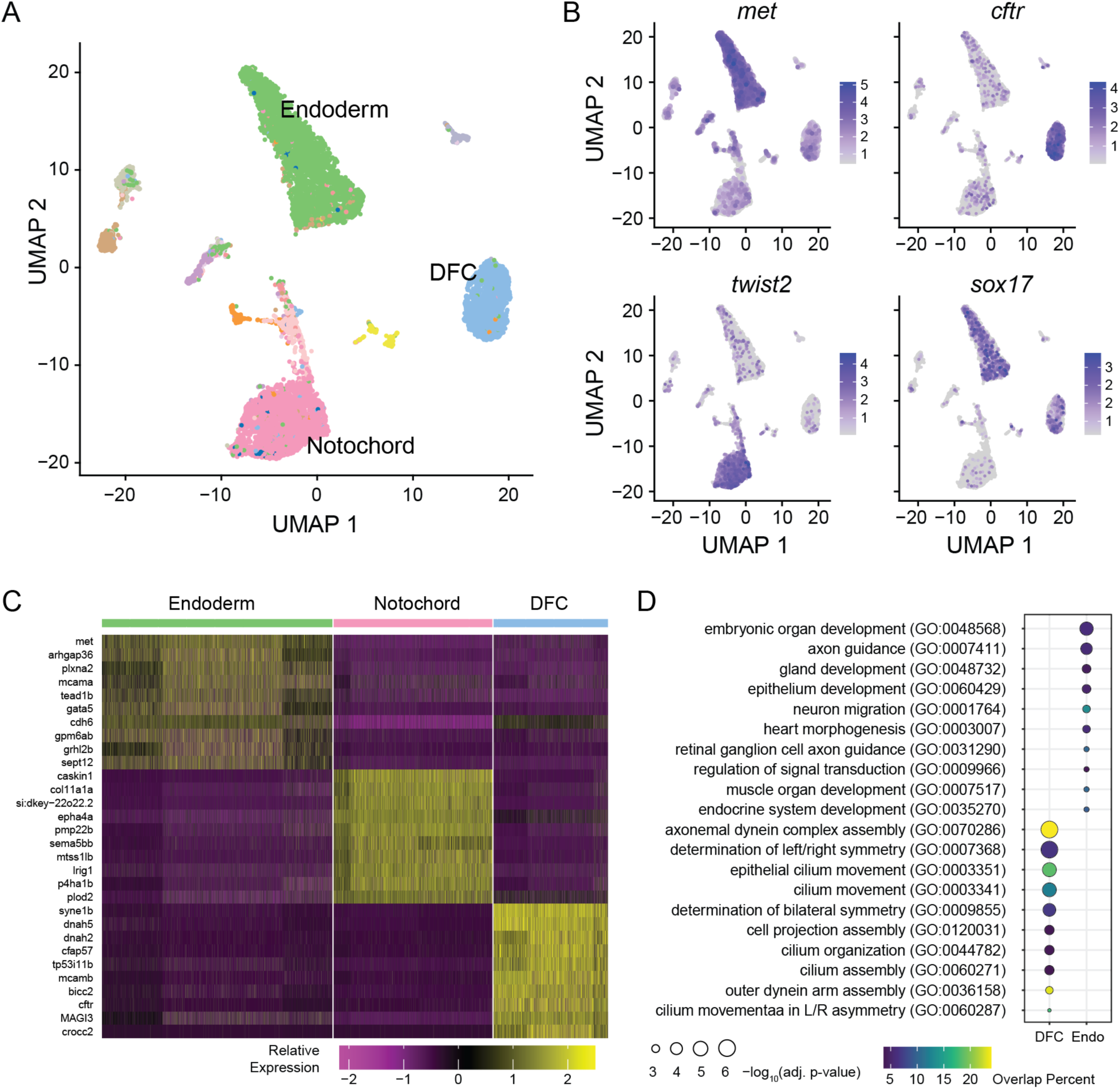
Integrated single-cell transcriptomic analysis of the zebrafish *sox17* lineage. A) Integrated UMAP projection of cells isolated from *Tg(sox17:GFP)*^S870^ embryos at 8, 10, and 12 hpf following unsupervised clustering. Clusters were annotated as endoderm, dorsal forerunner cells (DFCs), and notochord based on marker gene expression. B) Feature plots show expression of representative marker genes for endoderm (*met*), DFCs (*cftr*), and notochord (*twist2*). Feature plots are colored by normalized transcript abundance, with darker shades indicating higher expression. C) Heatmap showing the relative expression of the ten most discriminating genes for the endoderm, DFC, and notochord clusters. D) Gene Ontology (GO) term analysis of genes enriched in the endoderm and DFC clusters.

The minor clusters corresponded to additional embryonic cell types, including ectoderm (neural plate and non-neural ectoderm), mesoderm (prechordal plate and lateral plate mesoderm), the enveloping layer (EVL), and the YSL. Feature plots for all marker genes used in cluster annotation are available in Fig. S2. Except for the YSL, these cell types are not normally part of the *sox17* lineage, likely reflecting imperfect enrichment during FACS. Conversely, YSL cells were comparatively underrepresented, likely because the YSL is not fully cellularized during gastrulation and therefore is not efficiently recovered during FACS enrichment of intact GFP-positive cells. Overall, approximately 80% of recovered cells belonged to the endoderm, DFC, or notochord clusters, consistent with expectations based on our sorting strategy. As expected, we detected *sox17* transcript exclusively in the endoderm and DFC clusters; *sox17* transcript was largely absent from notochord and all minor clusters (Fig. 2B).

The top 10 genes that best discriminate between the major clusters, and thus drive cluster separation, are shown in the heat map in Fig. 2C. (Corresponding heat maps for the 8, 10, and 12 hpf datasets are provided in Fig. S3). Endoderm-enriched genes include *met*, the transcription factor *gata5*, an activator of *sox17* during gastrulation (Reiter et al., 1999; Reiter et al., 2001), and *cdh6*, a type II cadherin associated with migration (Clay and Halloran, 2014). Among the top 10 genes enriched in DFCs are several genes that function in ciliogenesis and left-right patterning, including genes encoding heavy chain dynein proteins (*dnah2*, *dnah5*) required for cilia beating (Yamaguchi et al., 2018), the RNA-binding protein *bicc2* (Bouvrette et al., 2010), and the cystic fibrosis transmembrane conductance regulator, *cftr (Navis et al., 2013)*. Finally, the notochord cluster was strongly enriched for the collagen gene *col11a1a* and *plod2*, which encodes a collagen-modifying enzyme, consistent with the well-established role of the extracellular matrix in the notochord (Parsons et al., 2002; Trapani et al., 2017).

Different functional terms were associated with the endoderm and DFC clusters (Sprague et al., 2003). The top 10 Gene Ontology (GO) terms associated with endoderm relate to cell migration, epithelium formation, and organ development (Fig. 2D). In contrast, GO terms associated with DFCs relate to ciliogenesis, cilia function, and left-right asymmetry, consistent with the known developmental roles of these cells. Together, these analyses indicate that our dataset faithfully captures the transcriptional signatures of endoderm and DFCs during and immediately following gastrulation.

At each stage, the most prominent clusters were endoderm, DFCs, and notochord (Fig. 3A); together these accounted for between 73% (8 hpf) and 88.7% (10 hpf) of all cells (Fig. 3B). Overall, the most common cell type was endoderm (36.3% of total), followed by notochord (24.9%). The proportion of notochord cells increased from 9.5% at 8 hpf to 42.6% at 12 hpf, reflecting progressive ectopic expression of the *Tg(sox17:GFP)* reporter in the notochord. As in the integrated dataset, the minor populations were diverse, and not all were detected at each timepoint; this suggests they are largely the result of permissive FACS settings. A fully annotated list of all clusters is available in Fig. S4.

**Figure 3.**
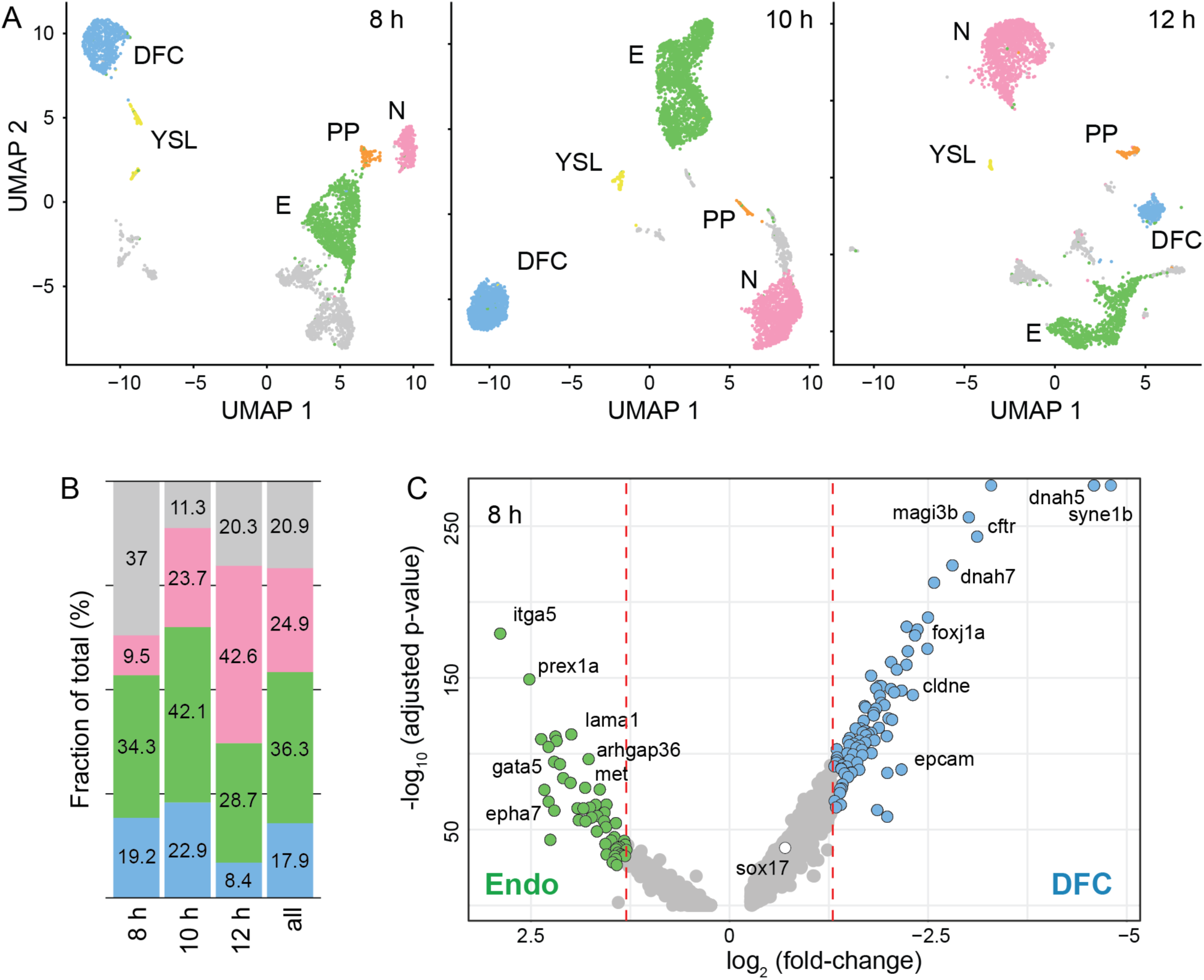
Transcriptional divergence within the sox17 lineage during gastrulation. A) UMAP projections of *sox17*:GFP-positive cells at mid-gastrulation (8 hpf), late gastrulation (10 hpf), and early somitogenesis (12 hpf). E, Endoderm. DFC, dorsal forerunner cells. N, notochord. YSL, yolk syncytial layer. PP, prechordal plate. B) Relative proportions of cells assigned to the endoderm (green), DFC (blue), and notochord (pink) clusters at each stage. C) Differential gene expression analysis comparing endoderm (Endo) and DFC clusters at mid-gastrulation (8 hpf). Selected genes are labeled to highlight endoderm enrichment for genes associated with cell migration and cell-matrix interactions and DFC enrichment for genes involved in epithelial organization and ciliogenesis.

At all time points, the major clusters were well separated in UMAP space and were not connected by obvious intermediate populations, indicating that their transcriptional profiles are distinct. Notably, we did not identify linkages between endoderm and DFCs even at our earliest time point (mid-gastrulation; Fig. 3A), suggesting that transitional transcriptional states between these populations were either absent or not captured in our dataset. Similarly, all three major clusters were largely isolated from the minor populations. Endoderm, DFC, and notochord clusters remained relatively compact throughout the sampling period, although the endoderm cluster at 12 hpf showed somewhat greater dispersion in transcriptional space (Fig. 3A). Thus, endoderm, DFC, and notochord maintained distinct global transcriptional identities throughout the sampling period. Because the notochord is not part of the *sox17* lineage we did not pursue further analysis of this cluster.

### Endoderm and DFC gene expression reflects their cell type-specific behavior

To compare gene expression in endoderm and DFCs, we performed differential gene expression analysis between these two clusters. Fig. 3C shows a representative volcano plot from the 8 hpf dataset, highlighting genes that distinguish endoderm from DFCs at the earliest stage analyzed. Fully annotated volcano plots for the 8, 10, and 12 hpf datasets are provided in Fig. S5. Across all time points, we found 86 genes to be enriched at least two-fold in endoderm over DFCs; conversely, 192 genes were enriched in DFCs over endoderm. An additional 40 genes were enriched in both endoderm *and* DFCs relative to the remaining dataset at the same threshold. Of these, only two genes (*cdh6*, *pltp*) were shared by endoderm and DFCs at all time points. The largest gene expression overlap was at bud stage (26/40), which is likely due to the somewhat higher number of cells captured at this time point. Regardless, the majority (87%; 86+192) of the 318 genes was preferentially expressed in either endoderm or DFCs.

Endoderm and DFCs differ sharply in the kinds of genes they activate. Endoderm was highly enriched in genes associated with signaling (22/86), extracellular matrix and adhesion (15/86), and transcriptional regulation (15/86). Among the transcription factors were *gata5*, a key regulator of endoderm specification (Reiter et al., 2001), and *tead1b*, suggesting a potential role for Hippo signaling during endoderm morphogenesis (Lin et al., 2017), as well as *tbx1* and *meis1b*, which are associated with anterior and pharyngeal patterning (Choe and Crump, 2014). Endoderm also expressed a number of signaling-related genes that directly regulate cell migration either as guidance cues or receptors (e.g., *met*, *nrp2a*, *robo3*) (Blockus and Chédotal, 2016; Scarpa and Mayor, 2016; Trusolino et al., 2010). Genes encoding extracellular matrix and adhesion proteins included the fibronectin *fn1a*, the fibrillin *fbn2*, three laminins (*lama1*, *lamb1a*, and *lamc1*), and the versican *vcana*. Notably, the integrin *itga5* is the most highly enriched gene at mid-gastrulation. Collectively, these gene expression patterns are consistent with the migratory behavior of endoderm cells during gastrulation and suggest that endoderm cells actively shape and interpret their extracellular environment.

In contrast, DFCs were highly enriched in genes associated with ciliogenesis and ciliary function (40/192 genes), consistent with their eventual differentiation into the ciliated epithelium of KV. DFC-enriched genes also included numerous regulators of cell-cell adhesion, epithelial polarity, and morphogenesis (approximately 25/192 genes), reflecting the collective migration and epithelial remodeling during KV formation (Forrest et al., 2022). In addition, DFCs preferentially expressed genes involved in protein synthesis and endoplasmic reticulum function, consistent with increased biosynthetic capacity associated with the assembly of a motile ciliated organ. Together, these data indicate that the DFC transcriptional program is specialized to support collective migration, epithelialization, and ciliogenesis well before KV forms.

In sum, the gene expression programs of endoderm and DFCs diverge early in gastrulation and reflect the distinct morphogenetic and developmental trajectories of these two populations.

### DFCs remain transcriptionally homogeneous during Kupffer’s vesicle morphogenesis

DFCs are a rare and transient cell population that is poorly represented in whole-embryo single-cell datasets. At the onset of gastrulation, embryos contain approximately 20–30 DFCs, and the mature KV comprises about 50 epithelial cells, although numbers vary with developmental stage and genotype (Matsui et al., 2015b; Moreno-Ayala et al., 2021; Oteiza et al., 2010). Across gastrulation and early somitogenesis, we recovered 2,081 DFCs, including 619 cells at 8 hpf, 1,195 cells at 10 hpf, and 267 cells at 12 hpf, providing sufficient resolution to characterize transcriptional changes associated with KV morphogenesis and function.

Temporal analysis of DFC gene expression revealed coordinated changes associated with the transition from collective migration to KV morphogenesis and function. These changes are summarized in a heatmap of the 100 most dynamic genes (Fig. 4A). At 8 hpf, DFC-enriched genes were associated with ciliogenesis, epithelialization, and polarity establishment. Notably, *foxj1a*, a well-established regulator of KV ciliogenesis (Caron et al., 2012; Tian et al., 2009; Yu et al., 2008), and *rfx2* (Neugebauer et al., 2013), a conserved activator of ciliary gene expression, were co-expressed with multiple cilia-associated genes (*dnah2*, *dnah5*, *hydin*, *crocc2*, *cfap57*), consistent with previous reports that ciliogenesis is initiated by mid-gastrulation (Aljiboury et al., 2022; Forrest et al., 2022). Genes involved in cytoskeletal organization and small GTPase signaling, including *dock11* and *rasgef1ba*, were also enriched, consistent with the extensive cell shape changes and tissue remodeling that accompany KV formation. In addition, the Eph receptor genes *epha2b* and *ephb4b* were enriched. Eph/ephrin signaling has been implicated in maintaining the boundary between DFCs and the neighboring endoderm and notochord (Zhang et al., 2016). DFCs further expressed junctional and polarity components, including the claudin *cldn7b* and *magi3a*, indicating that genes associated with KV epithelialization are expressed prior to KV morphogenesis.

**Figure 4.**
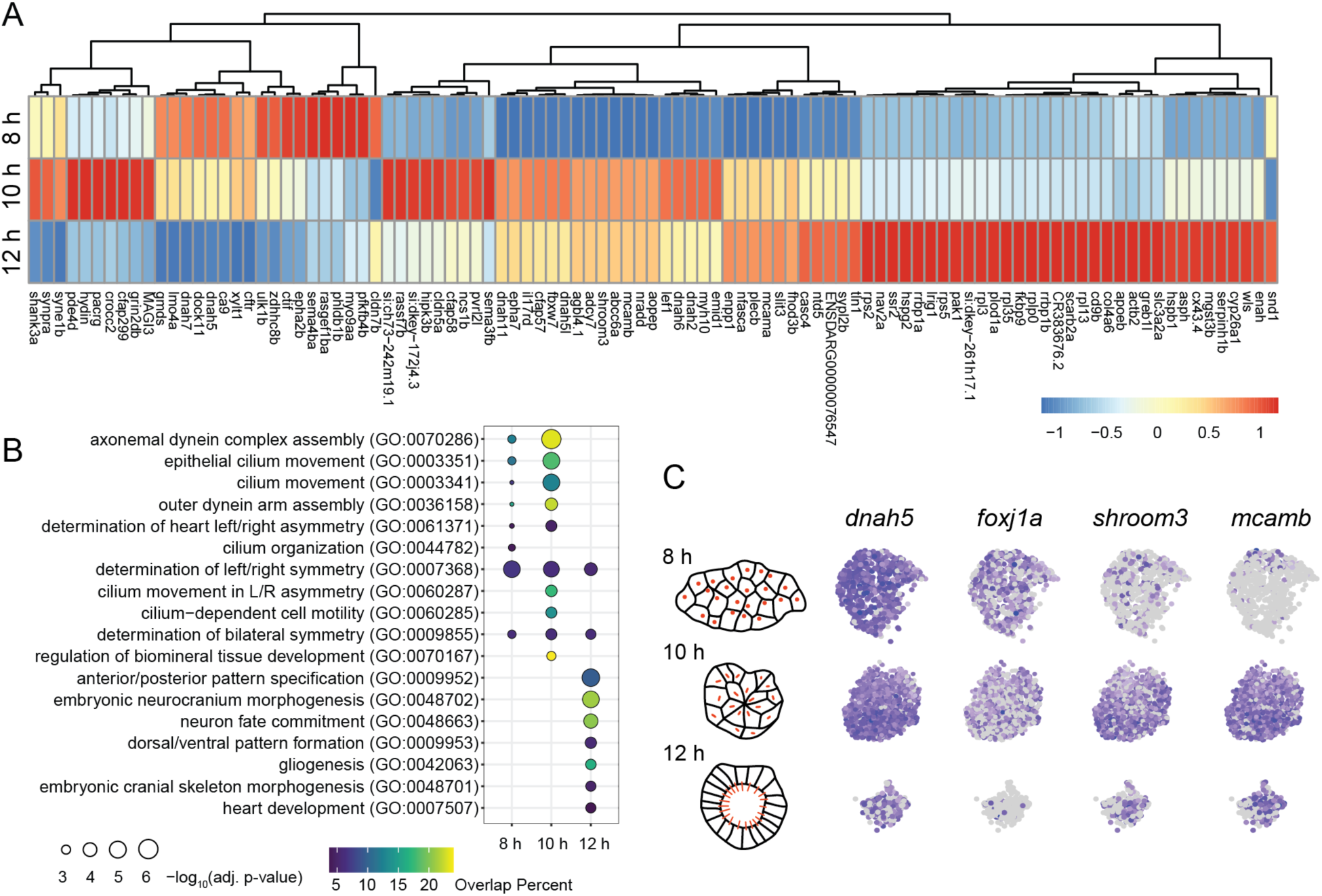
Transcriptional dynamics during Kupffer’s vesicle morphogenesis. A) Heatmap showing the relative expression of the 100 most dynamically expressed genes in dorsal forerunner cells (DFCs) across mid-gastrulation (8 hpf), late gastrulation (10 hpf), and early somitogenesis (12 hpf). B) GO term enrichment analysis of temporally regulated DFC genes reveals a shift in overrepresented biological processes coincident with Kupffer’s vesicle (KV) formation. C) Schematic illustrating the major morphogenetic transitions during KV formation, including collective migration of DFCs (8 hpf), rosette formation (10 hpf), and lumen inflation (12 hpf). Motile cilia (red) are assembled prior to lumen formation but are extruded into the lumen only after epithelialization. Representative feature plots show dynamic expression of DFC-enriched genes associated with migration, epithelial organization, ciliogenesis, and left-right patterning. Feature plots are colored by normalized transcript abundance, with darker shades indicating higher expression. Corresponding plots with quantitative color scales are provided in Fig. S6.

During rosette formation at 10 hpf, we observed an increase in expression of genes associated with apical constriction, epithelial organization, and cilium assembly. This included *shroom3*, a regulator of apical constriction (Haigo et al., 2003; Nishimura and Takeichi, 2008), together with *myh10, enah,* and *>od3b*, consistent with the coordinated cell shape changes required for rosette formation (Oteíza et al., 2008; Oteiza et al., 2010). DFCs also expressed adhesion and polarity genes (e.g., *mcamb, pvrl2l, cldn5a*) consistent with the establishment of apical-basal polarity. Genes associated with ion transport and KV lumen formation, such as *cftr* (Navis et al., 2013), were also expressed. Concurrently, expression of ciliary genes increased substantially, including additional axonemal dyneins (e.g., *dnah6, dnah11, dnah5l*) and cilia-associated proteins (*cfap57*, *cfap299*), consistent with active cilium assembly (Reiter and Leroux, 2017).

By 12 hpf, DFC gene expression shifted from genes associated with KV morphogenesis toward those associated with motile cilia function (e.g., *dnah5*, *dnah6*, *cfap57*) and left-right signaling. These included *dand5*, whose asymmetric degradation initiates left-right patterning, together with its post-transcriptional regulator *bicc2*. Additional signaling molecules expressed at this stage included *gdf11*, *fgf8a*, *notch1b*, and *shhb*.

Gene ontology analysis supported these temporal changes in gene expression. At 8 and 10 hpf, the most significantly enriched terms were associated with ciliogenesis and KV morphogenesis, including cilium organization, epithelial cilium movement, and determination of left-right asymmetry. By contrast, these categories were less strongly represented at 12 hpf, when terms associated with embryonic patterning and organogenesis, including anterior-posterior pattern specification, dorsal-ventral pattern formation, and heart development, became more prominent. Because fewer KV cells were recovered at 12 hpf, we cannot exclude the possibility that some ciliogenesis-associated transcripts remain expressed but were underrepresented in our dataset. Nevertheless, the observed shift in gene ontology terms is consistent with the transition from KV assembly to its role in left-right signaling.

Despite spanning the transition from migrating DFCs to a mature KV epithelium, we did not identify robust transcriptional subpopulations at any developmental stage. Instead, DFCs formed a single, compact cluster throughout the sampling period, suggesting that they collectively undergo coordinated changes in gene expression during KV morphogenesis. Feature plots of representative genes (Fig. 4C) associated with ciliogenesis (*foxj1a* and *dnah5*), epithelial morphogenesis (*shroom3*), and cell adhesion (*mcamb*) further supported temporally distinct expression dynamics while revealing no evidence of spatially restricted or transcriptionally distinct DFC subpopulations; a comprehensive set of feature plots is provided in Fig. S6.

### Endoderm transitions from a migratory to an epithelial transcriptional state

Temporal analysis of endoderm gene expression revealed a progressive transition from a migratory to an epithelial state. These changes are summarized in a heatmap of the 100 most dynamic genes in Fig. 5A. At 8 hpf, endoderm genes were enriched for factors associated with cell migration, guidance, and cytoskeletal regulation (e.g., *itga5*, *prex1a*, *robo3*, and *yap1*), consistent with the extensive dispersal and directional migration of endoderm cells during gastrulation. Endoderm cells also expressed *cdh2*, indicative of a mesenchymal-like state. A small set of genes peaked at 10 hpf, coinciding with the onset of epithelialization. These included *grhl2b*, a regulator of epithelial differentiation, *tead1b*, a transcriptional effector of Hippo signaling implicated in mechanosensing (Lin et al., 2017), and *met*, a receptor tyrosine kinase that promotes directional migration (Tu et al., 2026). qPCR analysis confirmed that *met* and *tead1b* transcript levels are near baseline at 6 hpf, when endoderm migration has already commenced, supporting the conclusion that these genes are transiently induced during convergence and epithelialization (Fig. S1). By 12 hpf, endoderm cells increasingly expressed epithelial and junctional components (e.g., *cdh1*, *krt8*, *f11r.1*, and *magi3a*), together with basement membrane and extracellular matrix-associated genes (e.g., *nid1b* and *col4a5*). The transition from *cdh2* to *cdh1* expression is consistent with progressive epithelialization of the endoderm (Nieto et al., 2016). Collectively, these temporal changes define a shared transcriptional program associated with the transition from migratory mesenchymal-like cells to a polarized epithelium.

**Figure 5.**
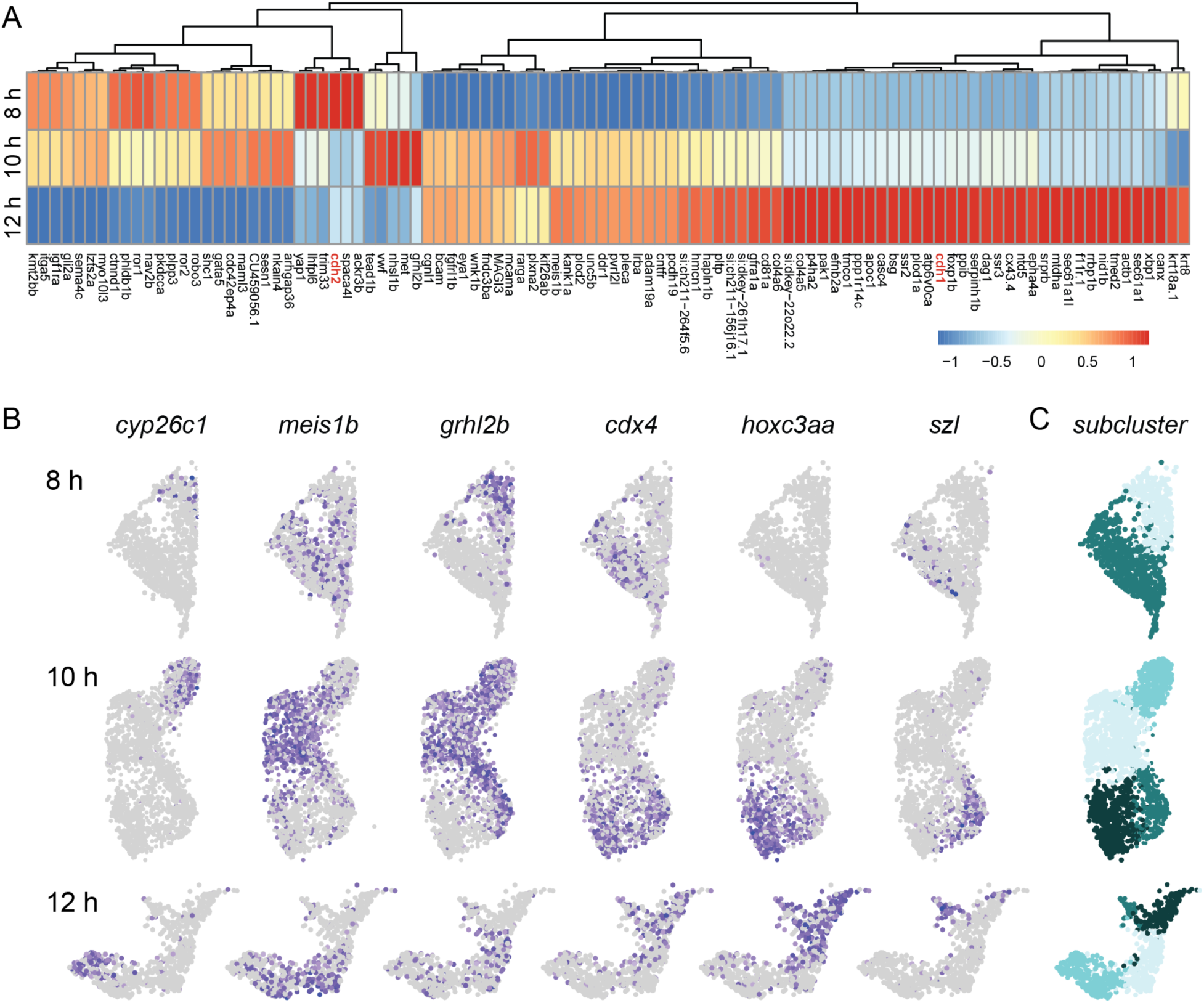
Endoderm transcriptional dynamics. A) Heatmap showing the relative expression of the 100 most dynamically expressed genes in endoderm cells across mid-gastrulation (8 hpf), late gastrulation (10 hpf), and early somitogenesis (12 hpf). *cdh2* and *cdh1* are highlighted in red, illustrating the transition from N-cadherin to E-cadherin expression during endoderm epithelialization. B) Feature plots showing representative endoderm-enriched genes exhibiting restricted expression patterns. Feature plots are colored by normalized transcript abundance, with darker shades indicating higher expression. Quantitative color scales are provided in Fig. S6. Unsupervised subclustering of endoderm cells identifies distinct transcriptional subclusters that emerge over the course of gastrulation and early somitogenesis.

### Transcriptional states of endoderm cells diverge during migration

Although endoderm cells collectively undergo a transition from a migratory mesenchymal-like state to a polarized epithelium, feature plots indicated that a significant number of endoderm-enriched genes are not broadly expressed but instead are restricted to distinct subsets of cells. Because our 10 hpf sample yielded the largest number of endoderm cells, we used this stage to illustrate representative expression patterns. At this stage, endoderm cells transition from directional migration to epithelial organization.

A small selection of genes with non-overlapping expression patterns is shown in Fig. 5B (a comprehensive set of feature plots is provided in Fig. S6). At bud stage (10 hpf), *cyp26c1* is enriched in a distinct subregion of the endoderm cluster that shows little or no *meis1b* expression. Conversely, *meis1b* is expressed throughout a neighboring subregion but is absent from a third *cdx4*-positive subregion. *hoxc3a* and *szl* exhibit expression patterns that partially overlap with *cdx4*. Some genes exhibit broader expression domains; for example, *grhl2b* is expressed throughout much of the endoderm but is excluded from the *hoxc3a*-positive region. These expression patterns persist into early somitogenesis (12 hpf), although boundaries between domains are less distinct, likely because fewer endoderm cells were captured at this stage. Notably, genes that exhibited restricted expression post-gastrulation were already restricted at mid-gastrulation. This suggests that transcriptional states diverge early rather than through progressive confinement of initially broad domains.

Although many endoderm-enriched genes exhibited restricted expression, genes associated with cell migration and morphogenesis were broadly expressed throughout the endoderm. These included *met*, the cytokine receptor *cxcr4a*, and the Rac-GEF *prex1a*, which promote endoderm migration (Mizoguchi et al., 2008; Nair and Schilling, 2008; Tu et al., 2026; Woo et al., 2012). These findings indicate that regional transcriptional diversification occurs alongside a shared morphogenetic program.

### Spatially distinct endoderm domains collectively span the post-gastrula embryo

We next subclustered endoderm cells to determine whether the transcriptional heterogeneity observed in feature plots reflected discrete cell states. At mid-gastrulation, endoderm resolved into two populations enriched for genes associated with posterior identity (including *cdx4* and *ved*) (Fig. 5C) or anterior identity (including *otx2a* and *tbx1*) (Fig. 5C; additional feature plots in S6). At post-gastrula stages, endoderm resolved into four clusters that corresponded to the expression domains observed in the feature plots. Similar clusters were identified at early somitogenesis, although the marker genes defining each cluster shifted over time (Fig. S7).

To determine whether these transcriptional states corresponded to spatially distinct embryonic domains, we performed HCR RNA-FISH for marker genes identified through feature plot and subclustering analyses (Fig. 6). We first examined *cdx4* because of its established role as a marker of posterior identity across multiple germ layers (Young and Deschamps, 2009). At mid-gastrulation, *cdx4* was expressed in a circumferential band around the margin (Fig. 6B,C). The *cdx4* expression domain extended farther toward the apical pole on the ventral than on the dorsal side. Consequently, dorsal endoderm cells, but not ventral endoderm cells, were located outside of the *cdx4*-positive domain, despite similar apical migration throughout the embryo (Fig. 6C). Post-gastrulation, *cdx4* became confined to the posterior half of the embryo, including the region surrounding KV (Fig. 6F).

**Figure 6.**
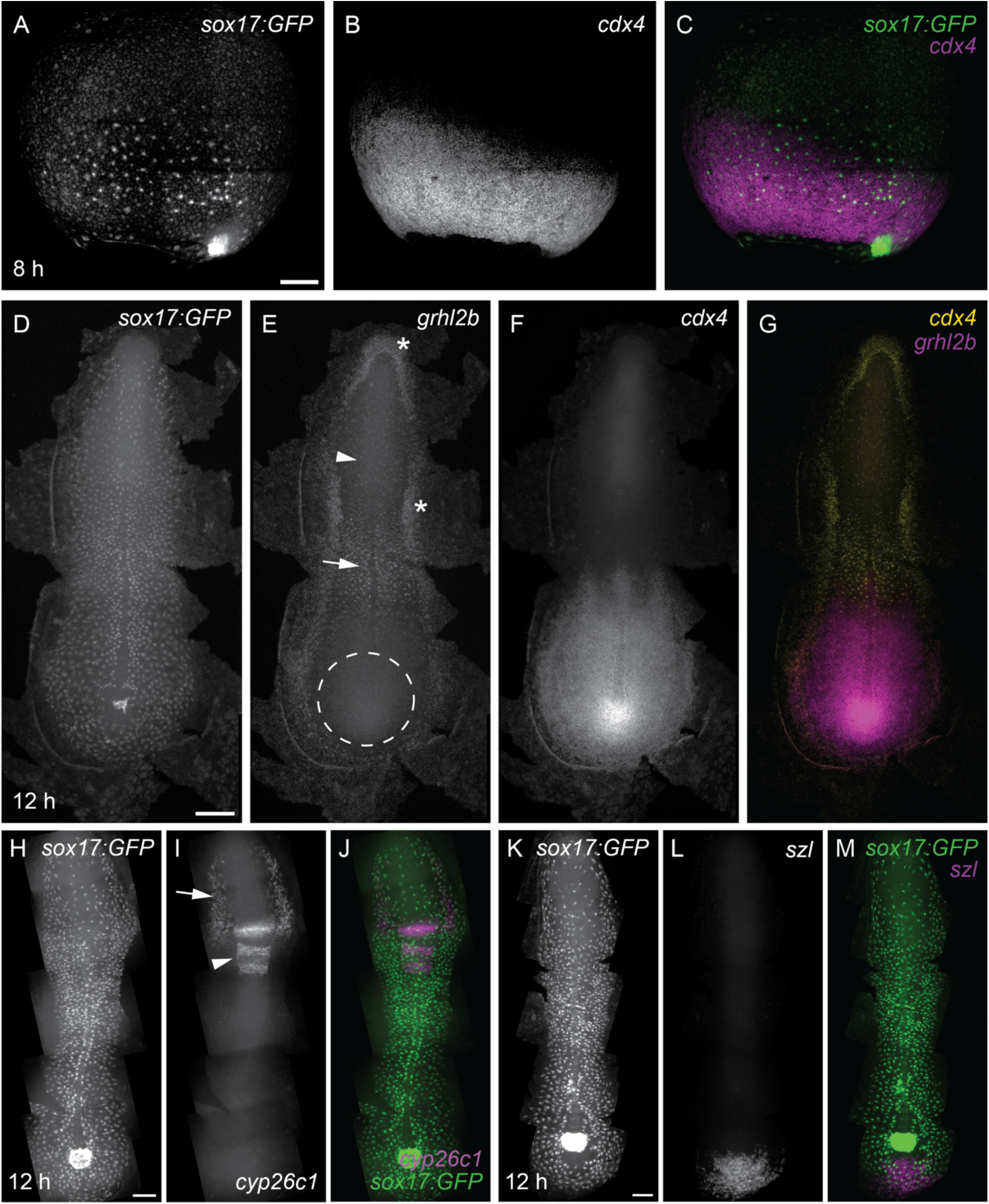
Spatial restriction of gene expression within the endoderm. A–C) Whole-mount HCR RNA-FISH at mid-gastrulation (8 hpf) reveals that endoderm cells, identified by *sox17:*GFP expression, adjacent to the blastoderm margin overlap the *cdx4* expression domain, whereas endoderm cells that have migrated away from the margin do not. Lateral views, dorsal to the right. D–M) Flat-mounted embryos at 12 hpf following HCR RNA-FISH; endoderm cells are identified by *sox17:GFP* expression. *grhl2b* is expressed throughout most of the endoderm (D–E, G), including anterior and central regions (arrows in E), but is excluded from a posterior domain surrounding Kupffer’s vesicle (KV) (dashed circle in E). *grhl2b* is also expressed in non-endodermal, cell-dense regions adjacent to the endoderm (asterisks in E). *cdx4* is expressed throughout the posterior endoderm and surrounding posterior tissues (F–G). *cyp26c1* expression is restricted to two anterior endoderm domains (H–J, arrows in I). *cyp26c1* is also detected in rhombomeres as previously reported (arrowheads in I). *szl* expression is confined to the caudal-most region of the embryo, including endoderm cells located posterior to KV (K–M). Scale bars A, H, K: 100 µm; D: 150 µm.

We next examined *grhl2b* and *szl*, whose expression patterns partially overlapped with *cdx4* in our sequencing data. Consistent with these findings, *grhl2b* was broadly but not uniformly expressed throughout the endoderm (Fig. 6E). In the posterior embryo, *grhl2b* expression overlapped with *cdx4* but was excluded from the domain immediately surrounding KV, consistent with distinct posterior-medial and posterior-lateral populations. Similarly, *szl*, a known marker of ventral and posterior embryonic territories, was restricted to the tailbud region posterior to KV, where it marked a subset of *cdx4*-positive endoderm cells (Fig. 6L).

We next examined *cyp26c1*, which feature plots identified as a marker of a distinct endoderm subpopulation. In addition to its known expression in rhombomeres and cardiac mesoderm (Rydeen and Waxman, 2014), we detected *cyp26c1* transcript in two bilateral domains within the anterior embryo. Our sequencing data indicated overlapping expression with *tbx1*, a transcription factor broadly associated with cardiopharyngeal development (Choe and Crump, 2014), although the *tbx1* expression domain was slightly larger (Fig. S6). Based on their location within the embryo and overlapping expression with *tbx1*, *cyp26c1*-positive endoderm cells likely correspond to pharyngeal endoderm (Wan et al., 2026).

Collectively, our clustering and HCR analyses identified four distinct endoderm populations that together span the post-gastrulation embryo: pharyngeal endoderm, marked by *cyp26c1*; anterior-medial endoderm, marked by *meis1b* and positioned between the *cyp26c1*-positive and *cdx4*-positive domains; posterior-lateral endoderm, comprising the anterior-most and lateral *cdx4*-positive endoderm; and posterior-medial endoderm, which surrounds KV and is encircled by the posterior-lateral domain (Fig. 7A,B). The transcriptional programs of the post-gastrulation domains are consistent with the broad anterior and posterior states identified at mid-gastrulation (8 hpf), suggesting that regional identities emerge during migration and are subsequently refined during epithelialization. In line with this interpretation, each domain is associated with a unique combination of enriched transcription factors (Fig. 7C), including *otx1*, *otx2a*, and *dmbx1a* in pharyngeal endoderm; *meis1b* and *meis2a* in anterior-medial endoderm; *hoxc3a* and *pax2a* in posterior-medial endoderm; and *ved* and *tbx3a* in posterior-lateral endoderm. These transcription factor profiles indicate that region-specific developmental programs are established early within the endoderm, with patterning beginning concurrently with migration.

**Figure 7.**
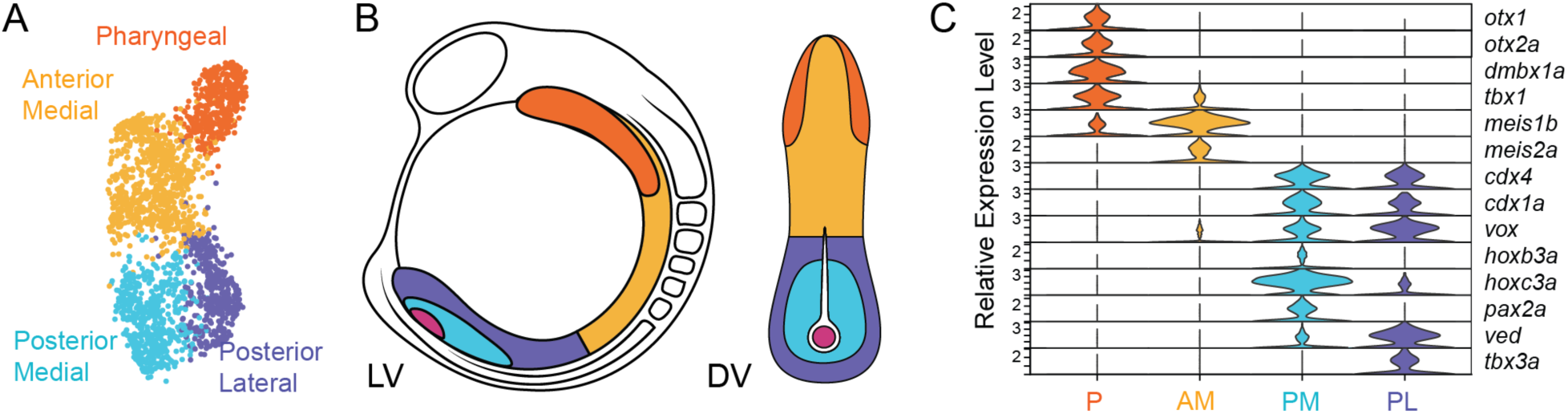
Endoderm regionalization at late gastrulation. A) UMAP projection of endoderm cells at late gastrulation (10 hpf) following unsupervised subclustering. Subclusters are annotated as pharyngeal, anterior-medial, posterior-lateral, and posterior-medial endoderm based on HCR RNA fluorescent in situ hybridization (HCR RNA-FISH) of marker genes. B) Schematic illustrating the inferred spatial organization of endoderm subregions in lateral (left) and dorsal (right) views during early somitogenesis. Colors correspond to the annotated subclusters in (A); the central fuchsia circle indicates Kupffer’s vesicle (KV). C) Violin plots showing the relative expression of representative transcription factors across endoderm subregions. Each subregion is characterized by a distinct combination of transcription factors.

## DISCUSSION

Here, we characterized the transcriptional dynamics of the zebrafish *sox17* lineage from mid-gastrulation through early somitogenesis, encompassing the period during which both endoderm and DFCs undergo extensive morphogenetic rearrangements. Our analysis reveals that, despite extensive overlap in their early transcriptional programs, DFCs and endoderm rapidly adopt distinct transcriptional states during gastrulation, consistent with their divergent behaviors and developmental fates. Whereas DFCs undergo coordinated temporal changes within a shared transcriptional program, endodermal cells progressively acquire distinct regional identities while maintaining a common set of migratory and epithelial characteristics. Our findings suggest that endoderm regionalization begins substantially earlier than previously appreciated and is tightly coupled to morphogenesis.

### Coordinated transcriptional changes drive Kupffer’s vesicle assembly

Endoderm and DFCs arise in response to similar inductive signals and initially express many of the same lineage-associated genes. Despite these similarities, the two populations rapidly diverge and establish distinct transcriptional identities during gastrulation. Previous studies have reported that a subset of cells initially associated with the DFC population can detach from the cluster and contribute to endodermal derivatives, suggesting that DFC and endoderm identities may remain flexible during gastrulation (Pulgar et al., 2021). However, we observed DFC and endoderm transcriptional states were clearly separated, with no intermediate populations detected, suggesting that the transcriptional programs associated with these identities are established early and maintained throughout gastrulation.

In contrast to the endoderm, we found little evidence for substantial transcriptional heterogeneity within the DFC population. Instead, DFCs remained largely transcriptionally uniform, while undergoing coordinated temporal changes in gene expression that paralleled the cellular events leading to KV formation. Genes associated with ciliogenesis were already strongly expressed by mid-gastrulation, despite the fact that cilia are not extruded until after KV inflation. DFC-enriched genes at this stage included numerous structural components of motile cilia, including dyneins and cilia-associated proteins, indicating that DFCs initiate the ciliary differentiation program well before KV assembly is complete. By late gastrulation, DFCs strongly express adhesion molecules, junctional proteins, and regulators of epithelial polarity that are required for lumen formation. These findings are consistent with recent studies suggesting that cilia are assembled within DFCs prior to lumen formation and only later extruded into the lumen (Aljiboury et al., 2022). More broadly, our data suggest that the extensive cellular rearrangements required for KV assembly are orchestrated through temporally regulated changes within a shared transcriptional program rather than through the emergence of distinct DFC states. In contrast to the progressive diversification observed within the endoderm, DFCs appear to rapidly activate functional effector programs downstream soon after their specification.

Despite the small size of the DFC population, our dataset captured more than 2,000 DFCs across gastrulation and early somitogenesis. Within this population, we found no evidence for robust transcriptional asymmetry prior to KV assembly. These findings suggest that transcriptional asymmetry within the left-right organizer arises only after KV assembly, rather than through pre-existing transcriptional differences within the DFC population (Blum et al., 2014; Djenoune et al., 2023; Forrest et al., 2022).

### Endoderm regionalizes during gastrulation

Considerable progress has been made in defining the gene regulatory networks that first establish endoderm identity (Economou et al., 2022; Figiel et al., 2021; Wilcockson et al., 2023; Zorn and Wells, 2009) and the later patterning processes that regionalize the gut tube and position endoderm-derived organs such as the liver and pancreas (Weatherbee et al., 2026). It is therefore surprising that comparatively little is known about the developmental interval separating these events. Our data suggest that the period during which endodermal cells disperse and converge on the midline represents a phase of active regionalization during which cells acquire a distinct molecular identity.

This regionalization is reflected in the emergence of spatially restricted transcriptional states that span the entire endoderm as it forms the initial epithelial sheet. Distinct transcription factor combinations within each domain suggest that discrete regulatory states drive regionalization. Their correspondence with known anterior-posterior patterning genes is consistent with the idea that these early states may establish a framework upon which later region-specific programs are built.

Classical fate-mapping experiments have shown that gastrula-stage endoderm along the dorsal-ventral axis has a distinct anterior-posterior bias (Kimmel et al., 1990; Melby et al., 1996; Warga and Nüsslein-Volhard, 1999). At first glance, this appears inconsistent with our finding that *cdx4* expression, a well-established marker associated with posterior fates, initially partitions the endoderm along the animal-vegetal axis. However, cdx4 expression is asymmetric along the dorsal-ventral axis; at mid-gastrulation, ventral endoderm remains largely within the *cdx4*-positive domain, whereas a substantial fraction of dorsal endoderm has migrated beyond the *cdx4* expression boundary. The cdx4 expression pattern is consistent with the positional biases identified by classical fate-mapping studies. Consistent with this interpretation, experimental manipulations that alter endoderm position relative to the blastoderm margin can shift the allocation of cells to anterior and posterior fates (Nair and Schilling, 2008; Norris et al., 2017; Schmid et al., 2013). Whether the early transcriptional states identified here represent stable fate decisions or more transient states remains unclear and will require approaches that integrate lineage tracing with transcriptional profiling across development.

### Regionalization emerges within a shared morphogenetic framework

Not all endoderm-enriched genes exhibited regionally restricted expression patterns. Many genes associated with endoderm migration remained broadly expressed across endodermal populations, including well-established regulators of migration such as *cxcr4a* (Mizoguchi et al., 2008; Nair and Schilling, 2008), *prex1a* (Woo et al., 2012), and *met* (Tu et al., 2026). Consistent with a central role for cell-ECM interactions during migration, genes encoding fibronectins and laminins were among the most highly enriched transcripts in the endoderm, suggesting that this tissue is itself a major source of extracellular matrix during gastrulation. Together, these findings suggest that endoderm cells may not simply respond to extracellular matrix cues through receptors such as Itga5 but also actively shape the extracellular environment through matrix deposition. Thus, while regional identities diverge during gastrulation, the molecular machinery that drives migration remains broadly shared across endodermal populations.

The broad expression of migration-associated genes raises questions about the regulatory mechanisms that establish this shared morphogenetic program. Many key genes, including *met* (Tu et al., 2026; this study) and *cxcr4a* (Mizoguchi et al., 2008), are not expressed at the onset of gastrulation and are activated only after endoderm has been specified. Moreover, these genes are among the first to be restricted to the endoderm rather than being concurrently expressed in neighboring tissues. Their delayed and tissue-specific expression suggests that activation of the endoderm migration program depends on additional transcriptional regulators acting downstream of endoderm specification. Identifying these inputs will be important for understanding how endodermal cells acquire the capacity to execute coordinated morphogenetic behaviors with precise temporal control during gastrulation.

### Morphogenesis and regionalization reinforce one another

In contrast to the broadly shared expression of migration-associated genes, regionally restricted transcripts suggest that neighboring endodermal cells acquire distinct capacities to interpret and respond to their local environments. Importantly, these transcriptional differences are not limited to factors associated with regional identity but also include genes implicated in cell adhesion, cytoskeletal organization, and signaling. Thus, regionalization may not simply arise as a consequence of morphogenesis; rather, emerging transcriptional differences could themselves reinforce or modify cell behaviors, creating a reciprocal relationship between cell movements and patterning.

In sum, our findings suggest that regional specification within the endoderm begins earlier than previously appreciated and is tightly coupled to morphogenesis. Rather than occurring sequentially, endoderm migration and regional transcriptional diversification may reinforce one another during gastrulation, establishing regional identities that serve as a foundation for patterning post-gastrulation.

## MATERIALS & METHODS

### Zebrafish

Adult zebrafish were maintained under standard laboratory conditions. All embryos used in this study were of an early stage prior to sex identification. Embryos were raised in egg water at 28.5°C. Zebrafish in an outbred background (*AB*/*TL*/*EKW*) were used as wild type. *Tg(sox17:GFP)^s870^* has been previously described (Mizoguchi et al., 2008). This study was performed with the approval of the Institutional Animal Care and Use Committee (IACUC) of the University of California, Merced (Protocol #2023–1144).

### Embryo Dissociation and Nuclei Isolation

To generate samples for single-nucleus library preparation, *Tg(sox17:GFP)^s870^* embryos were grown to the indicated stage (8, 10, or 12 hpf), and mechanically dechorionated by agitation in 0.5 mg/mL Pronase in 0.3× Ringer’s solution. Embryos were deyolked and dissociated as previously described (Manoli and Driever, 2012). GFP-positive cells were isolated by fluorescence-activated cell sorting using a FACSAria III cell sorter. Nuclei were isolated according to the 10x Genomics protocol CG000365, with lysis in 0.1× Lysis Buffer for 30 seconds.

### Library Preparation and Sequencing

snRNA-Seq libraries were prepared using a 10x Genomics Chromium Next GEM Single Cell Multiome ATAC + Gene Expression Kit. The manufacturer’s instructions were followed for cell capture, barcoding, template switch reverse transcription, cDNA amplification, and library construction. The targeted nuclei recovery of 8,000 cells per sample was selected for a projected sequencing depth of 25,000 paired reads per cell. Libraries were sequenced by SeqMatic (Fremont, CA) on an Illumina NovaSeq 6000 platform.

### Bioinformatic Alignment

All fastq files were processed using 10x Genomics Cell Ranger ARC pipelines (cellranger-arc v.2.0.0). Raw fastq files for gene expression (GEX) were processed and aligned using the `cellranger-arc count` command. We created our own reference package for zebrafish (GRCz11), modified to incorporate EGFP, using the `cellranger-arc mkref` command. We used Ensembl reference Danio_rerio.GRCz11.dna.primary_assembly.fa and Danio_rerio.GRCz11.104.chr.gtf for GEX. For each sample, counts were generated using `cellranger-arc count` with default parameters. Relevant scripts are available on GitHub (https://github.com/MaternaLab).

### Single-nuclei RNA Sequencing Data Analysis

Analyses were conducted with R (v4.2.2). Data preprocessing and analysis were performed using Seurat (v4.2.0) 100. Default parameters (`min.cells = 3`; `min.features = 200`) were used to convert the raw count matrices into Seurat objects. Cell quality control thresholds were determined empirically for each sample based on distributions of UMI counts, detected genes, and mitochondrial transcript content. At a minimum, we excluded cells with fewer than 500 unique molecular identifiers (UMIs), fewer than 300 genes, or greater than 20% mitochondrial transcripts. Genes expressed in fewer than ten cells across each sample were removed prior to downstream analysis. Doublets were identified and removed using DoubletFinder (v2.0.3) 101. Following filtering, we identified the 2000 highly variable genes (HVGs) for each sample using the variance stabilizing transformation (`vst`) method with the `FindVariableFeatures` function. Cell cycle scores were calculated using the `CellCycleScoring` function. Data scaling was performed using all genes, and the following variables were regressed out: number of UMIs, percent mitochondrial genes, and cell cycle. Because the dataset spans multiple developmental stages and transcriptionally heterogeneous populations, 50 principal components (PCs) were retained for dimensional reduction. Clusters were identified using the default Louvain algorithm with a community resolution of 2 and subsequently evaluated based on marker gene expression. Uniform Manifold Approximation and Projection (UMAP) dimensional reduction was performed after clustering to visualize cell clusters (Becht et al., 2018).

We used the `FindAllMarkers` function with the default Wilcoxon rank sum test to identify marker genes, making specific parameter adjustments (`log2FC.threshold = 0.1`; `min.diff.pct = 0.1`; `only.pos = TRUE`). Gene ontology analysis was performed using enrichR (v3.1) 102–104. All data wrangling and visualizations not generated with Seurat were generated using the tidyverse (v2.0.0) 105. Differential expression tables were generated using the `FindMarkers` function. Zebrafish basic gene information, expression data for wild-type fish, mouse and zebrafish orthology, and human and zebrafish orthology were obtained from the ZFIN Data Reports (Sprague et al., 2003).

### Hybridization chain reaction RNA-fluorescence in situ hybridization

Whole mount RNA in situ hybridization analysis was performed by HCR RNA-FISH as previously described (Ibarra-García-Padilla et al., 2021) on wild-type or *Tg(sox17:GFP)^s870^* embryos; GFP fluorescence survived fixation and processing, allowing for identification of endodermal cells without the need for additional labeling. Briefly, embryos at the indicated stages were fixed in 4% paraformaldehyde overnight at 4°C, rinsed with phosphate-buffered saline (PBS), and dehydrated in 100% methanol overnight at -20°C. Prior to hybridization, embryos were sequentially rehydrated with methanol in PBST. HCR RNA-FISH amplifiers, buffers and probes against zebrafish were obtained from Molecular Instruments. Specifically, HCR RNA probes against *sox17* (NM_131287.2), *cdx4* (NM_131109.2), *grhl2b* (NM_001083072.1), *cyp2Cc1* (NM_001029951.2), and *szl* (NM_181663.1) were designed based on the zebrafish genome assembly (GRCZ11/danRer11).

### Microscopy, image analysis, and image processing

For HCR RNA-FISH imaging, embryos were incubated in 5X saline-sodium citrate buffer plus 0.1% Tween-20 (SSCT) supplemented with 30% glycerol and flat mounted on glass slides as previously described (Cheng et al., 2014).

Images were acquired with a 10x/0.4 NA or 30x/1.05 NA objective lens on a microscope (IX83; Olympus) equipped with a spinning disk confocal unit (CSU-W1; Andor), a scientific complementary metal–oxide–semiconductor (sCMOS) camera (Prime 95b; Teledyne Photometrics), and MicroManager software. Z-stacks were acquired at 2 µm intervals. For HCR RNA-FISH imaging, multiple tiled Z-stacks were acquired to capture the entire sphere of embryo, or entire length of flat-mounted embryo.

Image analysis and processing was performed using Fiji software (Schindelin et al., 2012). All subsequent processing was performed from maximum projections of confocal Z-stacks. For HCR RNA-FISH imaging, at least 3 embryos were analyzed per gene. For tiled images, maximum Z-projections were stitched together using the pairwise stitching function in Fiji.

### qPCR Analysis

Absolute transcript abundance was determined by qPCR as previously described (Ligunas and Materna, 2026). Briefly, known quantities of in vitro-transcribed GFP and mCherry RNA were added to each sample immediately prior to RNA extraction and used as external spike-in standards. Total RNA was extracted using the Qiagen RNeasy Micro Kit with on-column DNase digestion, and 1 µg of RNA was reverse transcribed using the qScript cDNA Synthesis Kit (Quantabio).

Quantitative PCR was performed using PerfeCTa SYBR Green FastMix (Quantabio) on a Jena Analytik qTOWER instrument. Reactions were performed in triplicate using 1 µl of cDNA per reaction. Transcript abundance was calculated from ΔC_t_ values relative to the combined mean C_t_ of the GFP and mCherry and converted to absolute transcript numbers based on the known quantity of spike-in RNA added to each sample. Primer specificity was verified by melt-curve analysis and primer efficiencies were between 95% and 100%. Sequences of all gene-specific primers are listed in Table S1.

## ACKNOWLEDGEMENTS

We thank the UC Merced Vivarium staff for animal care and Dr. Gravano for assistance with cell sorting. We thank Dr. Christian Mosimann for many helpful discussions and suggestions. We are grateful to the members of the Materna and Woo labs for their insights and support.

S.W., S.C.M., and J.D.T. were supported by NIH grant R15HD102829. O.A.D. and K.K.H. were supported by NIH grant R15HL146779-01S1. C.B.L. was supported by NIH grant R01AA031043.

## SUPPLEMENTAL FIGURE LEGENDS

**Figure S1. Absolute transcript quantification during early zebrafish development.** Absolute transcript abundance of *cdx4, eomes, met, otx1, sox17, sox32,* and *tead1b* was measured by quantitative PCR in whole embryos lysates from 0 to 12 hpf relative to an external standard (Ligunas and Materna, 2026). Maternal transcripts (*eomes* and *otx1*) declined rapidly and were largely undetectable before the onset of gastrulation, whereas transcripts associated with endoderm specification, migration, and regionalization (*sox32, sox17, met, tead1b,* and *cdx4*) were activated prior or during gastrulation. Transcript abundance is reported as transcripts per embryo. Circles and triangles represent independent biological replicates; lines indicate the mean.

**Figure S2. Feature plots of marker gene expression used for cluster annotation.** UMAP feature plots showing expression of representative marker genes used to assign identities to the major cell populations captured in the *Tg(sox17:GFP)* dataset. Expression levels are overlaid on the integrated UMAP shown in Fig. 2. Color intensity indicates normalized transcript abundance.

**Figure S3. Cluster-defining gene expression.** Heatmaps showing the top 10 differentially expressed genes for each cluster identified at 8, 10, and 12 hpf, as well as the integrated dataset. Genes were ranked by differential expression relative to all other clusters. Color intensity indicates scaled expression levels.

**Figure S4. Fully annotated UMAPs for individual developmental stages.** UMAPs showing all annotated cell populations identified at 8, 10, and 12 hpf. In contrast to the simplified annotations shown in Fig. 3, all cluster identities are displayed for each developmental stage. Colors correspond to cluster identities shown in Fig. 2.

**Figure S5. Differential gene expression between endoderm and dorsal forerunner cells.** Volcano plots showing differential gene expression between endoderm and dorsal forerunner cells at 8, 10, and 12 hpf. The x-axis indicates log2 fold change, and the y-axis shows the adjusted *P* value (−log10-transformed). Genes exhibiting a ≥2-fold difference in expression (|log2 fold change| ≥ 1.3) are highlighted and annotated.

**Figure S6. Feature plots of genes enriched in the sox17 lineage.** UMAP feature plots showing the expression patterns of genes enriched in endoderm and dorsal forerunner cells relative to the remaining dataset across all developmental stages. Expression levels are overlaid on the UMAPs corresponding to the 8, 10, and 12 hpf datasets, with color intensity indicating normalized transcript abundance.

**Figure S7. Differentially expressed genes defining endoderm subclusters.** Heatmaps showing genes differentially expressed among endoderm subclusters identified at 8 hpf, 10 hpf, and 12 hpf. These heatmaps correspond to the subclustering analyses shown in Figs. 5 and 7 and highlight the transcriptional programs underlying regional diversification of the endoderm. Genes were ranked by differential expression relative to the remaining endoderm subclusters at each developmental stage. Color intensity indicates scaled expression levels.

**Table S1:**
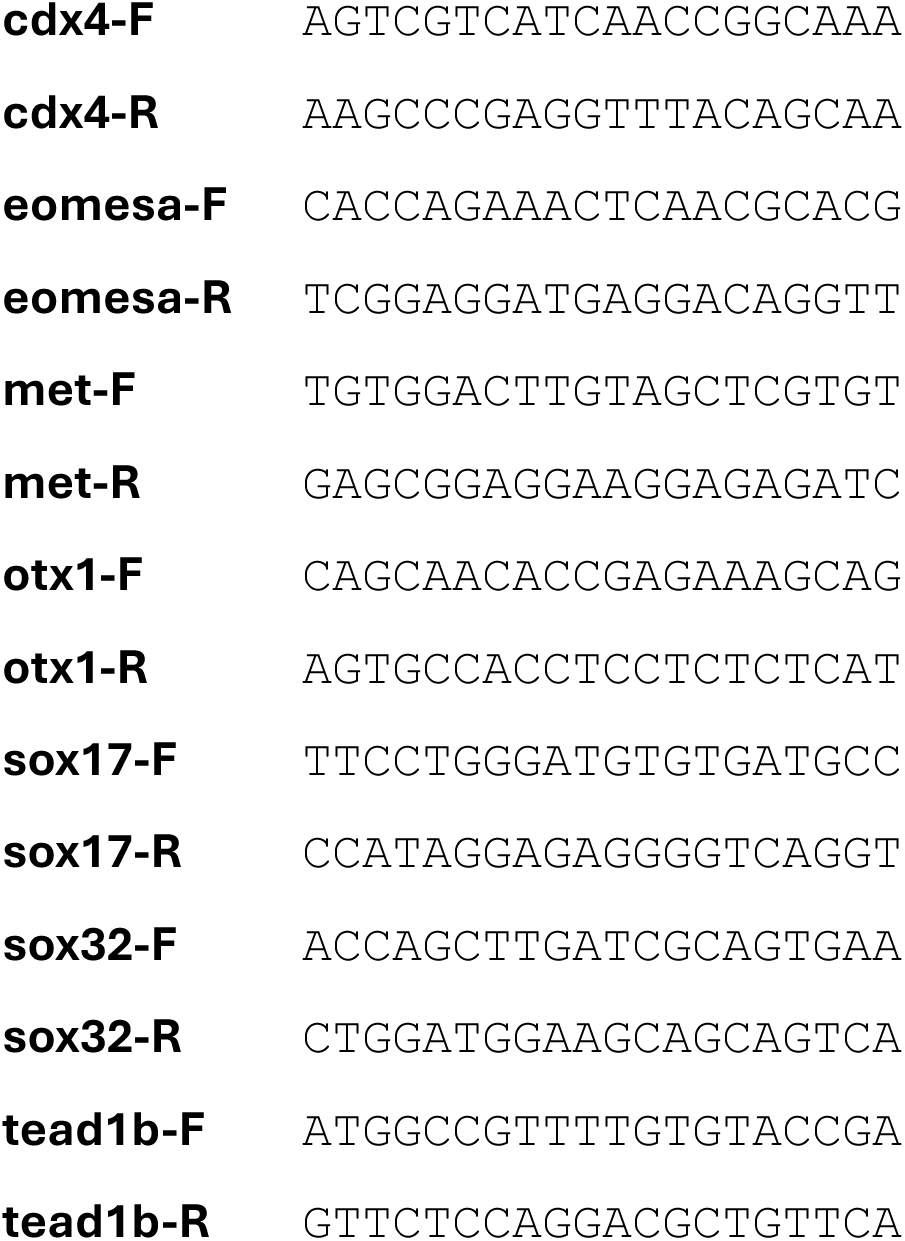
Sequence of qPCR primers used for time course measurements in Fig. S1. All sequences shown in 5′→3′ orientation.

