## Supplemental Figures for "Transcriptional divergence of the zebrafish *sox17* lineage begins during gastrulation"

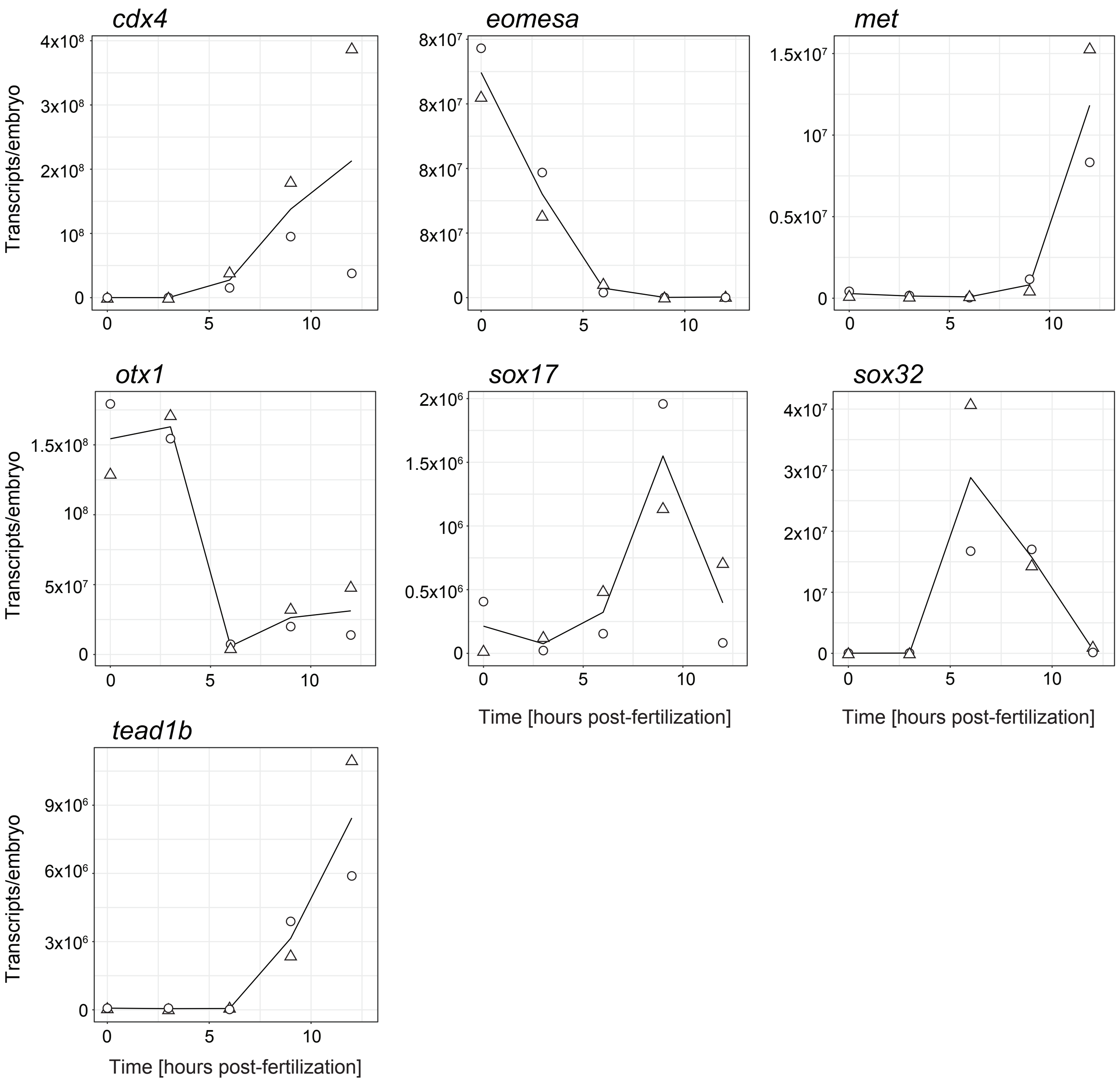

cdx4
eomesa
met
otx1
sox17
sox32
tead1b
Transcripts/embryo
Time [hours post-fertilization]
4x10^8
3x10^8
2x10^8
10^8
0
8x10^7
1.5x10^7
10^7
0.5x10^7
1.5x10^8
5x10^7
2x10^6
1.5x10^6
10^6
0.5x10^6
4x10^7
3x10^7
2x10^7
9x10^6
6x10^6
3x10^6
5
10

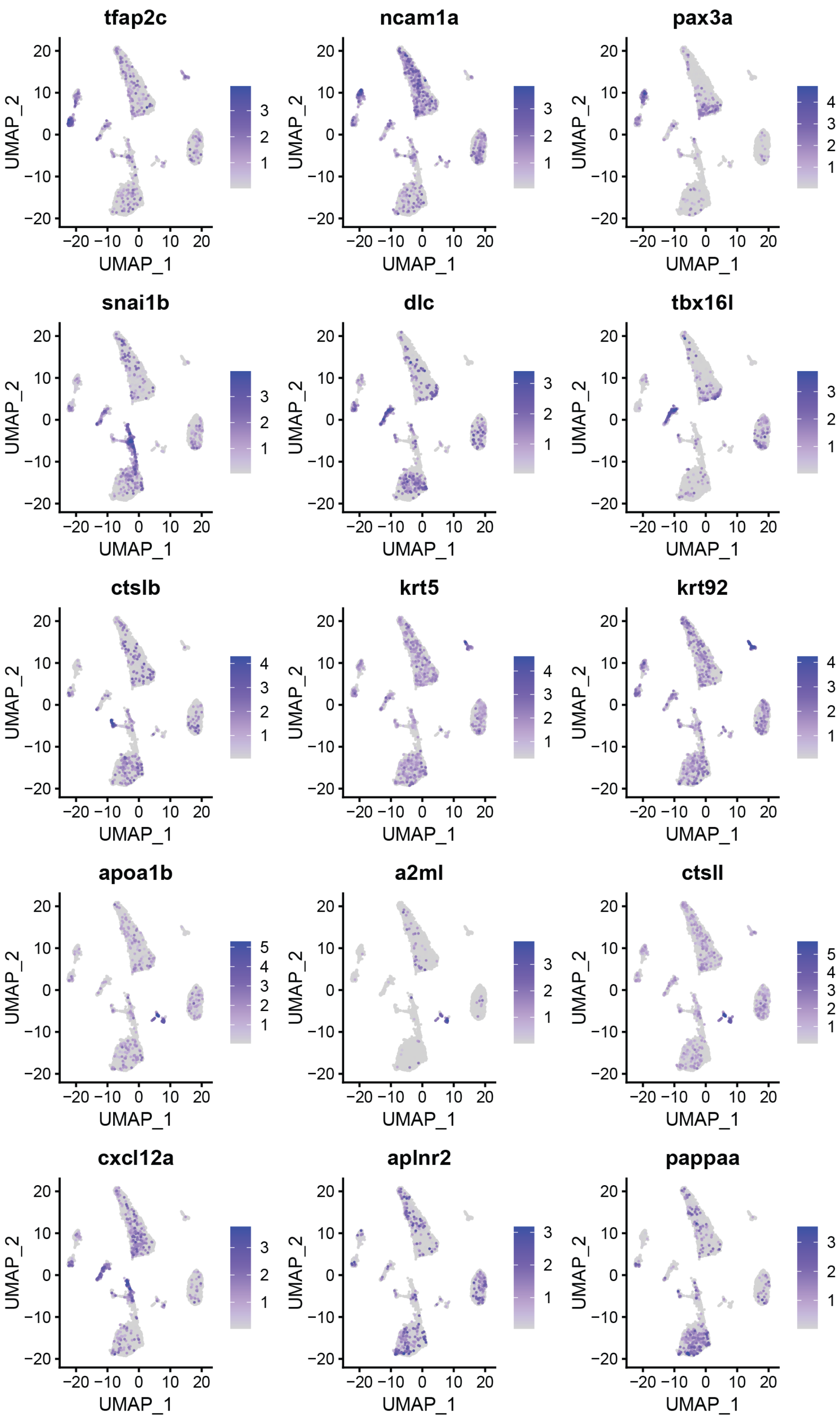

tfap2c
ncam1a
pax3a
snai1b
dlc
tbx16l
ctslb
krt5
krt92
apoa1b
a2ml
ctsll
cxcl12a
aplnr2
pappaa
UMAP_1
UMAP_2

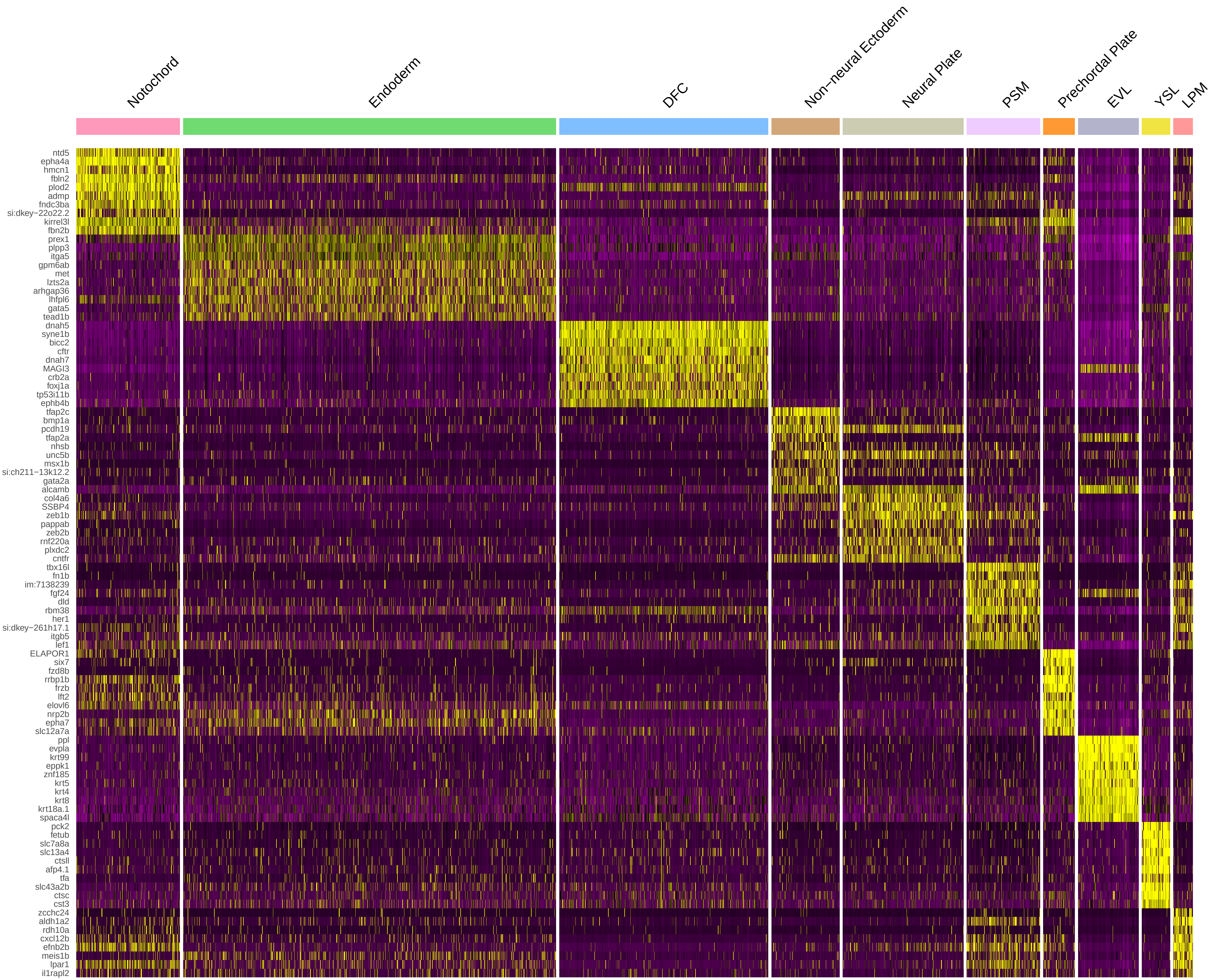
Notochord
Endoderm
DFC
Non−neural Ectoderm
Neural Plate
PSM
Prechordal Plate
EVL
YSL
LPM
ntd5
epha4a
hmcn1
fbln2
plod2
admp
fndc3ba
si:dkey−22o22.2
kirrel3l
fbn2b
prex1
plpp3
itga5
gpm6ab
met
lzts2a
arhgap36
lhfpl6
gata5
tead1b
dnah5
syne1b
bicc2
cftr
dnah7
MAGI3
crb2a
foxj1a
tp53i11b
ephb4b
tfap2c
bmp1a
pcdh19
tfap2a
nhsb
unc5b
msx1b
si:ch211−13k12.2
gata2a
alcamb
col4a6
SSBP4
zeb1b
pappab
zeb2b
rnf220a
plxdc2
cntfr
tbx16l
fn1b
im:7138239
fgf24
dld
rbm38
her1
si:dkey−261h17.1
itgb5
lef1
ELAPOR1
six7
fzd8b
rrbp1b
frzb
lft2
elovl6
nrp2b
epha7
slc12a7a
ppl
evpla
krt99
eppk1
znf185
krt5
krt4
krt8
krt18a.1
spaca4l
pck2
fetub
slc7a8a
slc13a4
ctsll
afp4.1
tfa
slc43a2b
ctsc
cst3
zcchc24
aldh1a2
rdh10a
cxcl12b
efnb2b
meis1b
lpar1
il1rapl2

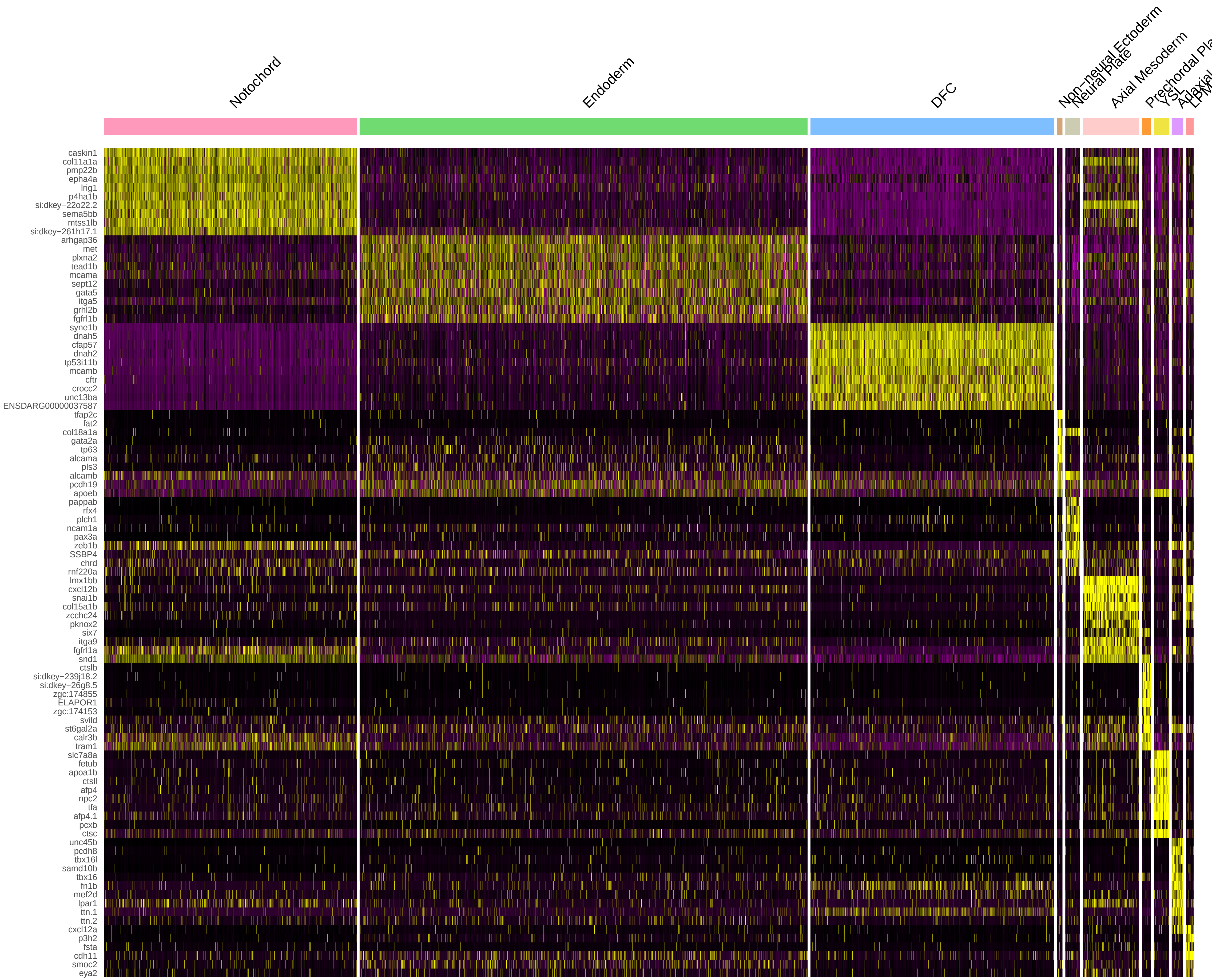
Notochord
Endoderm
DFC
Non−neural Ectoderm
Neural Plate
Axial Mesoderm
YSL
caskin1
col11a1a
pmp22b
epha4a
lrig1
p4ha1b
si:dkey−22o22.2
sema5bb
mtss1lb
si:dkey−261h17.1
arhgap36
met
plxna2
tead1b
mcama
sept12
gata5
itga5
grhl2b
fgfrl1b
syne1b
dnah5
cfap57
dnah2
tp53i11b
mcamb
cftr
crocc2
unc13ba
ENSDARG00000037587
tfap2c
fat2
col18a1a
gata2a
tp63
alcama
pls3
alcamb
pcdh19
apoeb
pappab
rfx4
plch1
ncam1a
pax3a
zeb1b
SSBP4
chrd
rnf220a
lmx1bb
cxcl12b
snai1b
col15a1b
zcchc24
pknox2
six7
itga9
fgfrl1a
snd1
ctslb
si:dkey−239j18.2
si:dkey−26g8.5
zgc:174855
ELAPOR1
zgc:174153
svild
st6gal2a
calr3b
tram1
slc7a8a
fetub
apoa1b
ctsll
afp4
npc2
tfa
afp4.1
pcxb
ctsc
unc45b
pcdh8
tbx16l
samd10b
tbx16
fn1b
mef2d
lpar1
ttn.1
ttn.2
cxcl12a
p3h2
fsta
cdh11
smoc2
eya2

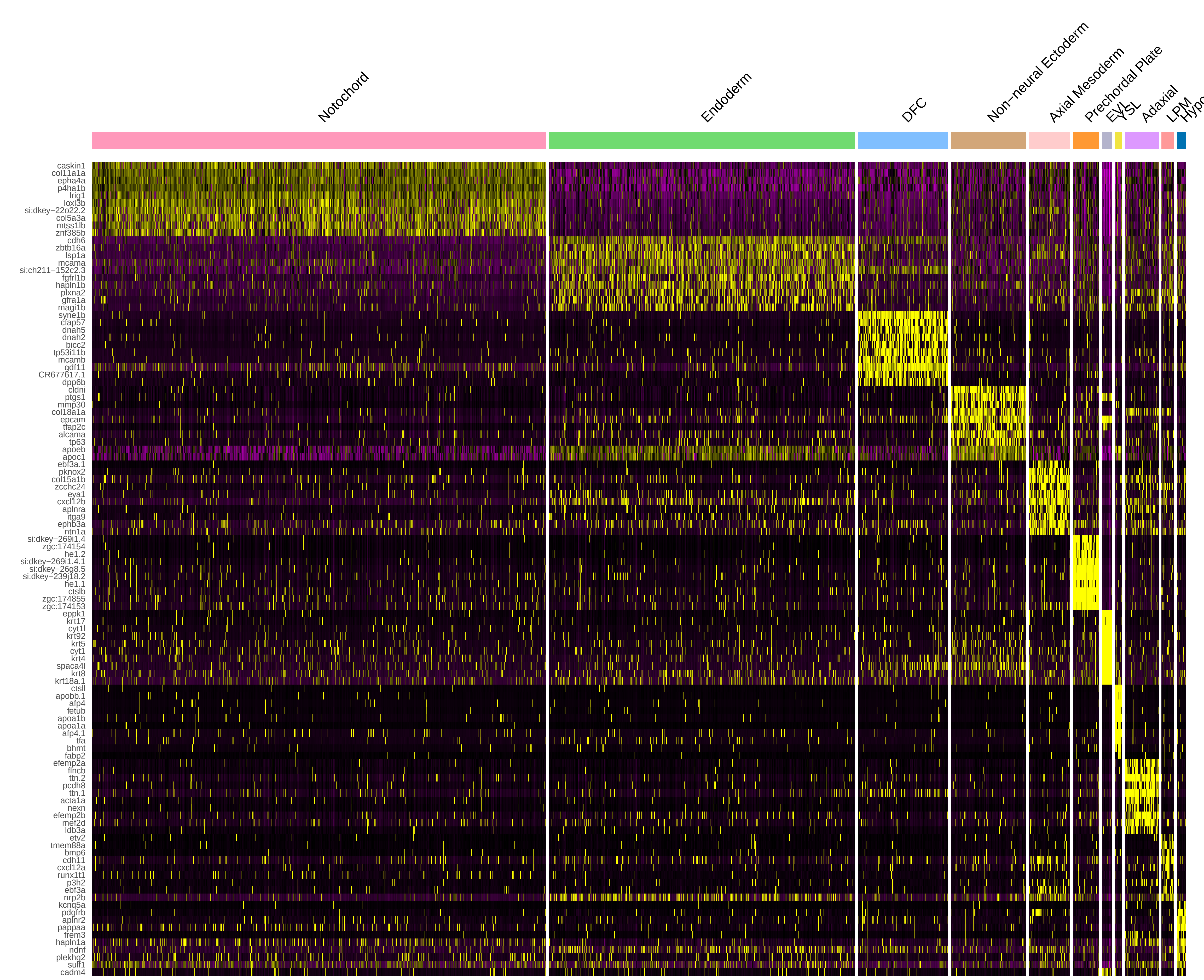
Notochord
Endoderm
DFC
Non−neural Ectoderm
Axial Mesoderm
Prechordal Plate
EVL
YSL
Adaxial
LPM
caskin1
col11a1a
epha4a
p4ha1b
lrig1
loxl3b
si:dkey−22o22.2
col5a3a
mtss1lb
znf385b
cdh6
zbtb16a
lsp1a
mcama
si:ch211−152c2.3
fgfrl1b
hapln1b
plxna2
gfra1a
magi1b
syne1b
cfap57
dnah5
dnah2
bicc2
tp53i11b
mcamb
gdf11
CR677617.1
dpp6b
cldni
ptgs1
mmp30
col18a1a
epcam
tfap2c
alcama
tp63
apoeb
apoc1
ebf3a.1
pknox2
col15a1b
zcchc24
eya1
cxcl12b
aplnra
itga9
ephb3a
ntn1a
si:dkey−269i1.4
zgc:174154
he1.2
si:dkey−269i1.4.1
si:dkey−26g8.5
si:dkey−239j18.2
he1.1
ctslb
zgc:174855
zgc:174153
eppk1
krt17
cyt1l
krt92
krt5
cyt1
krt4
spaca4l
krt8
krt18a.1
ctsll
apobb.1
afp4
fetub
apoa1b
apoa1a
afp4.1
tfa
bhmt
fabp2
efemp2a
flncb
ttn.2
pcdh8
ttn.1
acta1a
nexn
efemp2b
mef2d
ldb3a
etv2
tmem88a
bmp6
cdh11
cxcl12a
runx1t1
p3h2
ebf3a
nrp2b
kcnq5a
pdgfrb
aplnr2
pappaa
frem3
hapln1a
ndnf
plekhg2
sulf1
cadm4

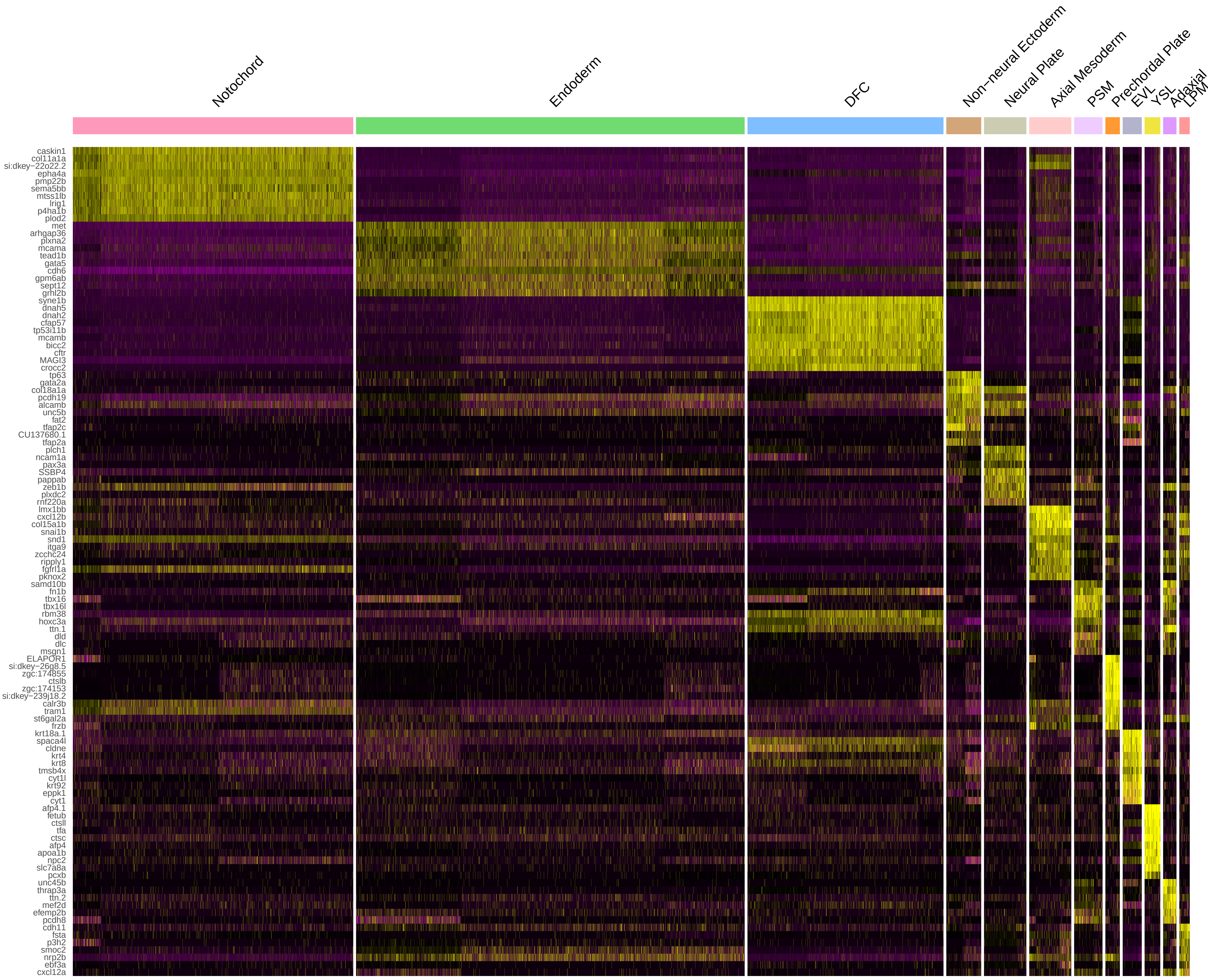

Notochord
Endoderm
DFC
Non−neural Ectoderm
Neural Plate
Axial Mesoderm
PSM
Prechordal Plate
EVL
YSL
Adaxial
LPM
caskin1
col11a1a
si:dkey−22o22.2
epha4a
pmp22b
sema5bb
mtss1lb
lrig1
p4ha1b
plod2
met
arhgap36
plxna2
mcama
tead1b
gata5
cdh6
gpm6ab
sept12
grhl2b
syne1b
dnah5
dnah2
cfap57
tp53i11b
mcamb
bicc2
cftr
MAGI3
crocc2
tp63
gata2a
col18a1a
pcdh19
alcamb
unc5b
fat2
tfap2c
CU137680.1
tfap2a
plch1
ncam1a
pax3a
SSBP4
pappab
zeb1b
plxdc2
rnf220a
lmx1bb
cxcl12b
col15a1b
snai1b
snd1
itga9
zcchc24
ripply1
fgfrl1a
pknox2
samd10b
fn1b
tbx16
tbx16l
rbm38
hoxc3a
ttn.1
dld
dlc
msgn1
ELAPOR1
si:dkey−26g8.5
zgc:174855
ctslb
zgc:174153
si:dkey−239j18.2
calr3b
tram1
st6gal2a
frzb
krt18a.1
spaca4l
cldne
krt4
krt8
tmsb4x
cyt1l
krt92
eppk1
cyt1
afp4.1
fetub
ctsll
tfa
ctsc
afp4
apoa1b
npc2
slc7a8a
pcxb
unc45b
thrap3a
ttn.2
mef2d
efemp2b
pcdh8
cdh11
fsta
p3h2
smoc2
nrp2b
ebf3a
cxcl12a

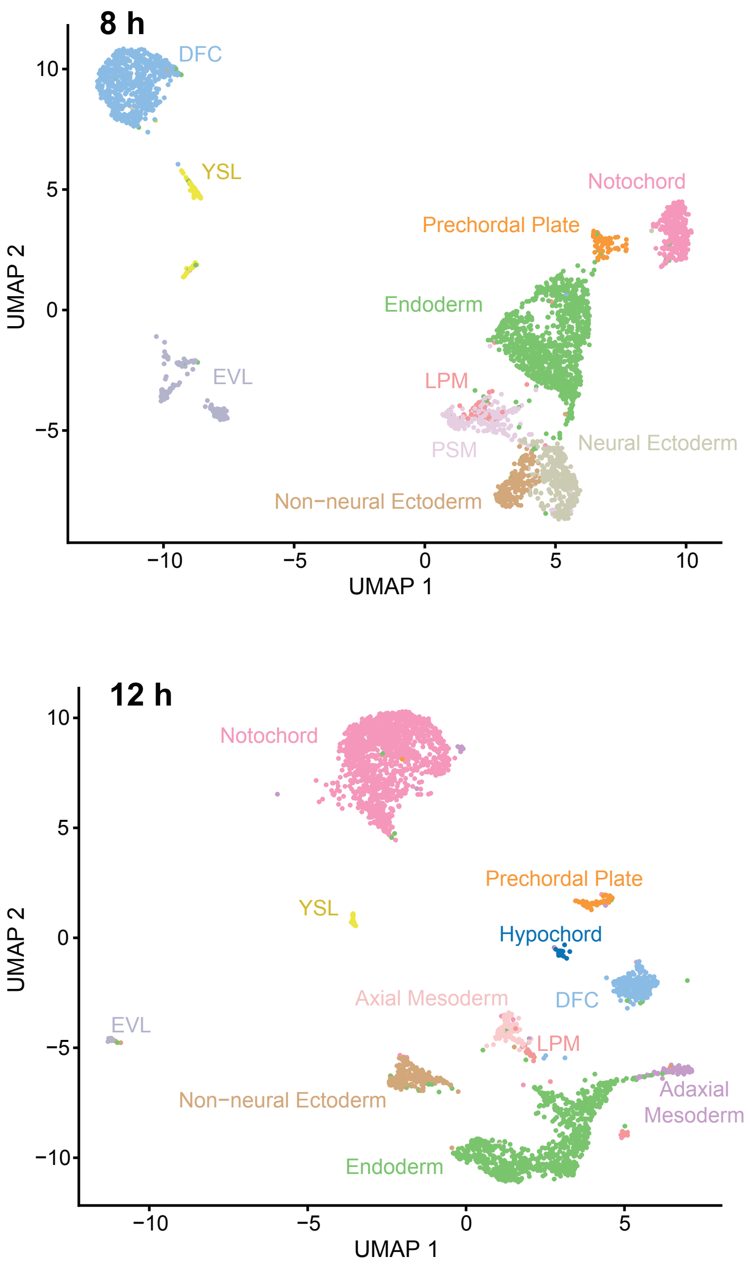


DFC—dorsal forerunner cells
EVL—enveloping layer
LPM—lateral plate mesoderm
PSM—presomitic mesoderm
YSL—yolk syncytial layer

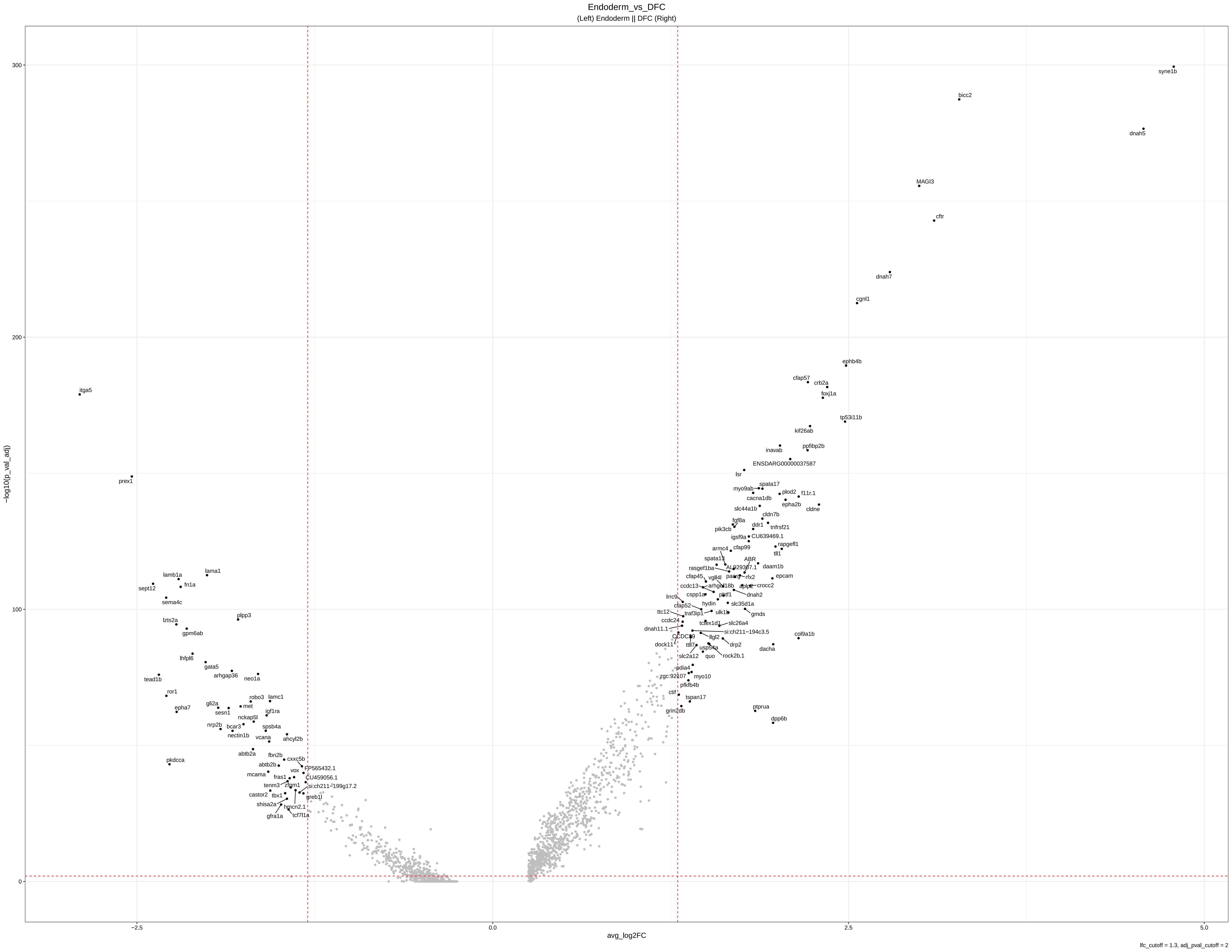
Endoderm_vs_DFC
(Left) Endoderm || DFC (Right)
300
200
100
0
−log10(p_val_adj)
-2.5
0.0
2.5
5.0
avg_log2FC
lfc_cutoff = 1.3, adj_pval_cutoff = 2
syne1b
bicc2
dnah5
MAGI3
cftr
dnah7
cgnl1
ephb4b
cfap57
crb2a
foxj1a
tp53i11b
kif26ab
inavab
ppfibp2b
ENSDARG00000037587
lsr
spata17
myo9ab
plod2
f11r.1
cacna1db
epha2b
cldne
slc44a1b
cldn7b
fgf8a
pik3cb
ddr1
tnfrsf21
igsf9a
CU639469.1
rapgefl1
armc4
cfap99
tll1
spata13
ABR
daam1b
rasgef1ba
AL929307.1
cfap45
vgll4l
pacrg
rfx2
epcam
ccdc13
arhgef18b
aplp2
crocc2
cspp1a
pltf1
dnah2
lrrc9
hydin
slc35d1a
cfap52
ttc12
traf3ip1
ulk1b
gmds
ccdc24
tctex1d1
slc26a4
dnah11.1
si:ch211−194c3.5
CCDC39
llgl2
col9a1b
dock11
ttll7
drp2
usp54a
dacha
slc2a12
quo
rock2b.1
pdia4
zgc:92107
myo10
pfkfb4b
ctif
tspan17
grin2db
ptprua
dpp6b
itga5
prex1
lamb1a
lama1
sept12
fn1a
sema4c
plpp3
lzts2a
gpm6ab
lhfpl6
gata5
arhgap36
neo1a
tead1b
ror1
robo3
lamc1
gli2a
met
sesn1
epha7
igf1ra
nckap5l
nrp2b
bcar3
spsb4a
nectin1b
vcana
ahcyl2b
abtb2a
fbn2b
cxxc5b
pkdcca
abtb2b
FP565432.1
vox
mcama
fras1
CU459056.1
tenm3
zfpm1
si:ch211−199g17.2
castor2
tbx1
greb1l
shisa2a
hmcn2.1
gfra1a
tcf7l1a

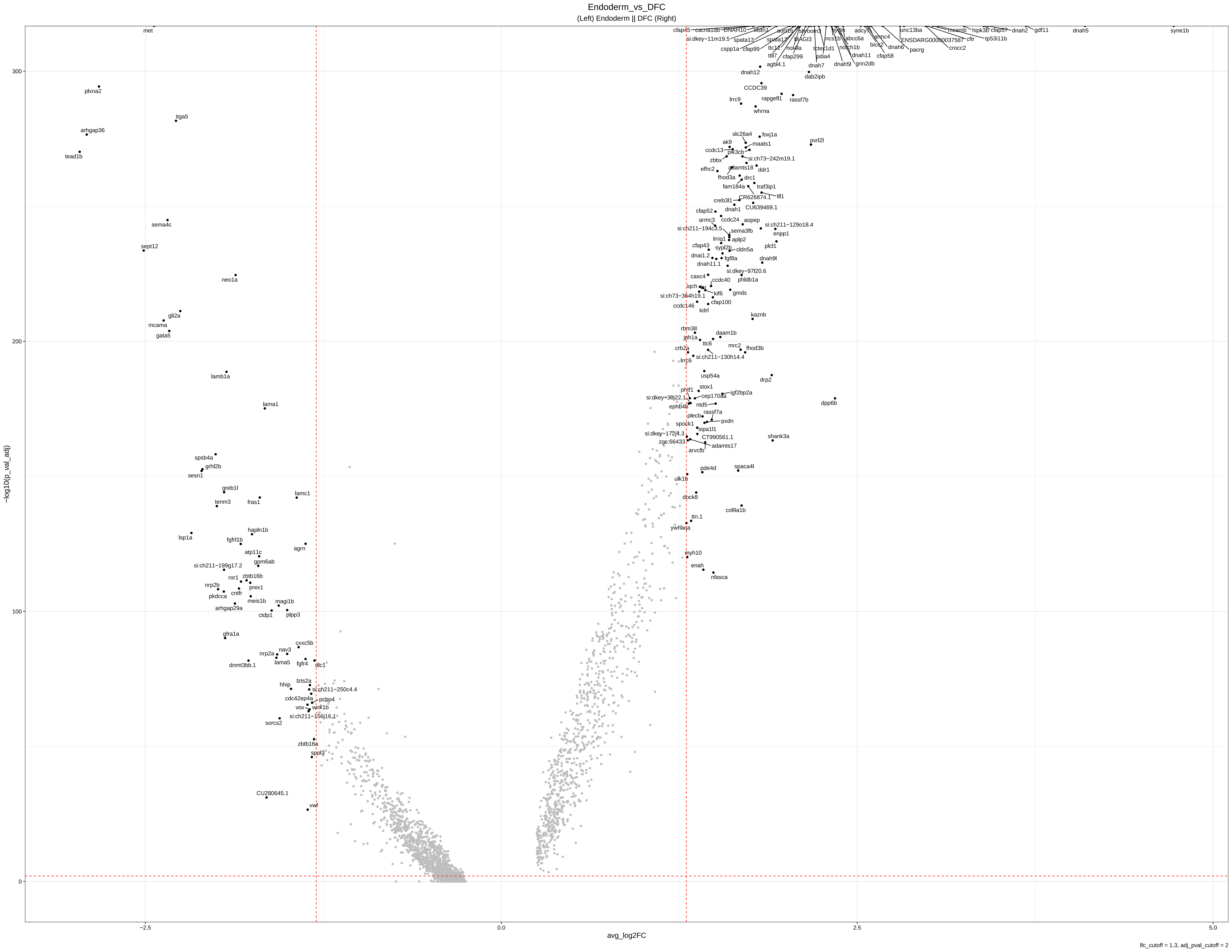

Endoderm_vs_DFC
(Left) Endoderm || DFC (Right)
−log10(p_val_adj)
avg_log2FC
lfc_cutoff = 1.3, adj_pval_cutoff = 2
300
200
100
0
−2.5
0.0
2.5
5.0
met
plxna2
itga5
arhgap36
tead1b
sema4c
sept12
neo1a
gli2a
mcama
gata5
lamb1a
lama1
spsb4a
grhl2b
sesn1
greb1l
tenm3
fras1
lamc1
lsp1a
hapln1b
fgfrl1b
agrn
atp11c
si:ch211−199g17.2
gpm6ab
ror1
zbtb16b
nrp2b
prex1
pkdcca
cntfr
meis1b
magi1b
arhgap29a
ctdp1
plpp3
gfra1a
cxxc5b
nrp2a
nav3
lama5
fgfr4
dlc1
dnmt3bb.1
lzts2a
hhip
si:ch211−250c4.4
cdc42ep4a
pcbp4
vox
wnk1b
si:ch211−156j16.1
sorcs2
zbtb16a
sppl3
CU280645.1
vwf
cfap45
cacna1db
DNAH10
elsm1
si:dkey−11m19.5
spata13
spata17
shroom3
MAGI3
cspp1a
cfap99
ttc12
nol4la
ttll7
cfap299
tctex1d1
agbl4.1
pdia4
ncs1b
abcc6a
notch1b
dnah11
dnah5l
grin2db
armc4
bicc2
dnah6
cfap58
pacrg
unc13ba
ENSDARG00000037587
cftr
crocc2
mcamb
hipk3b
cfap57
dnah2
gdf11
tp53i11b
dnah5
syne1b
dnah7
dab2ipb
dnah12
CCDC39
rapgefl1
rassf7b
lrrc9
whrna
slc26a4
foxj1a
ak9
maats1
pvrl2l
ccdc13
pik3cb
zbbx
si:ch73−242m19.1
efhc2
adamts18
ddr1
fhod3a
drc1
fam184a
traf3ip1
creb3l1
CR626874.1
tll1
dnah1
CU639469.1
cfap52
ccdc24
armc3
aopep
si:ch211−129o18.4
si:ch211−194c3.5
sema3fb
enpp1
lrrig1
aplp2
cfap43
sypl2b
cldn5a
plcl1
dnai1.2
fgf8a
dnah9l
dnah11.1
si:dkey−97l20.6
casc4
ccdc40
phldb1a
iqch
kif6
gmds
si:ch73−364h19.1
cfap100
ccdc146
kdrl
kaznb
rbm38
daam1b
jph1a
ttc6
mrc2
fhod3b
crb2a
si:ch211−130h14.4
lrrc6
usp54a
drp2
stox1
phtf1
igf2bp2a
si:dkey−30j22.1
cep170aa
dpp6b
ephb4b
ntd5
plecb
rassf7a
pxdn
spock1
sipa1l1
si:dkey−172j4.3
CT990561.1
shank3a
zgc:66433
adamts17
arvcfb
pde4d
spaca4l
ulk1b
dock8
col9a1b
ttn.1
ywhaqa
myh10
enah
nfasca

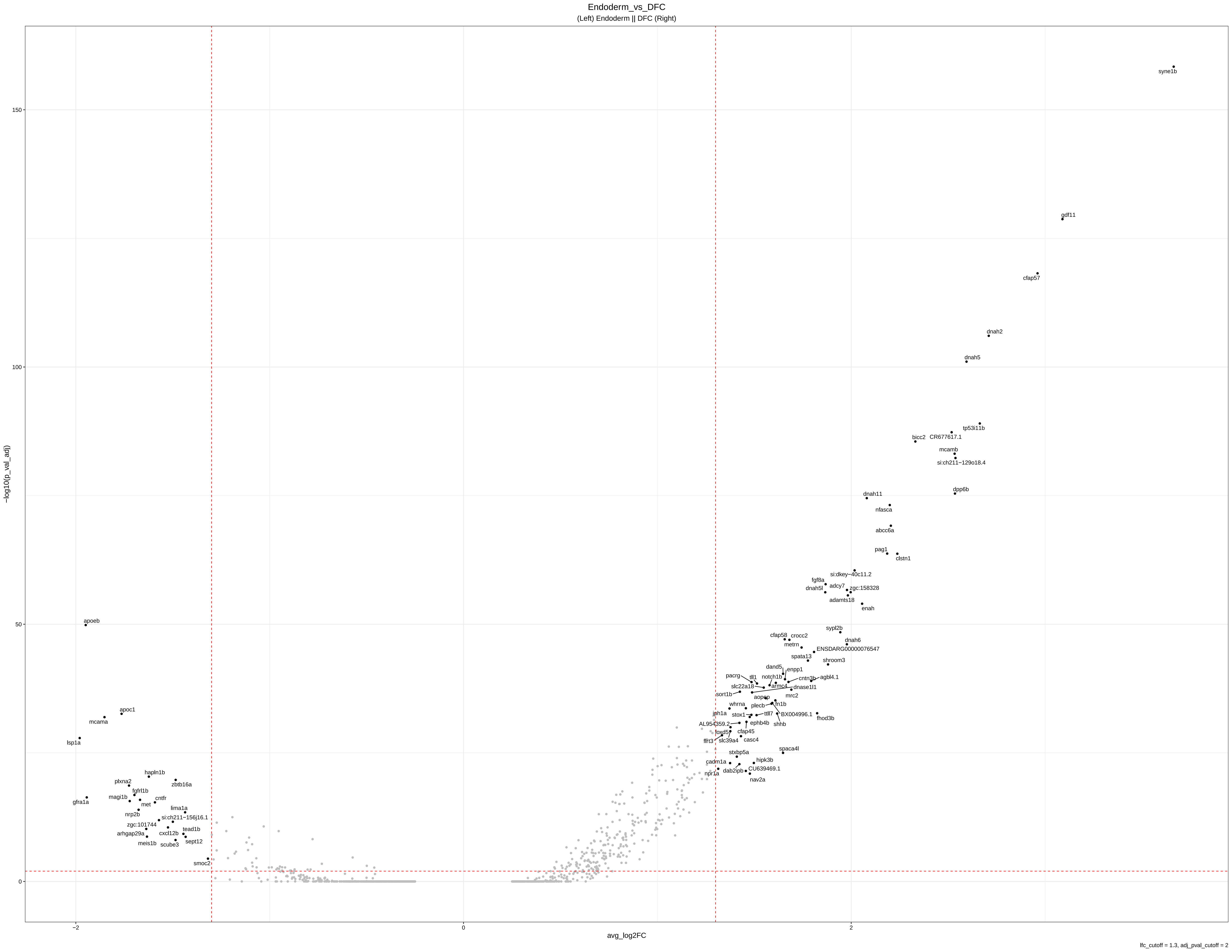
Endoderm_vs_DFC
(Left) Endoderm || DFC (Right)
−log10(p_val_adj)
avg_log2FC
150
100
50
0
−2
0
2
syne1b
gdf11
cfap57
dnah2
dnah5
tp53i11b
CR677617.1
bicc2
mcamb
si:ch211−129o18.4
dpp6b
dnah11
nfasca
abcc6a
pag1
clstn1
si:dkey−40c11.2
fgf8a
adcy7
dnah5l
zgc:158328
adamts18
enah
sypl2b
cfap58
crocc2
dnah6
metrn
ENSDARG00000076547
spata13
shroom3
dand5
enpp1
pacrg
tll1
notch1b
cntn3b
agbl4.1
slc22a18
armc4
dnase1l1
sort1b
aopep
mrc2
whrna
plecb
fn1b
jph1a
stox1
ttll7
BX004996.1
fhod3b
shhb
AL954359.2
ephb4b
foxd5
cfap45
flrt3
slc39a4
casc4
spaca4l
stxbp5a
hipk3b
cadm1a
dab2ipb
CU639469.1
npr1a
nav2a
apoeb
apoc1
mcama
lsp1a
hapln1b
plxna2
zbtb16a
fgfrl1b
magi1b
cntfr
gfra1a
met
lima1a
nrp2b
si:ch211−156j16.1
zgc:101744
tead1b
arhgap29a
cxcl12b
meis1b
scube3
sept12
smoc2
lfc_cutoff = 1.3, adj_pval_cutoff = 2

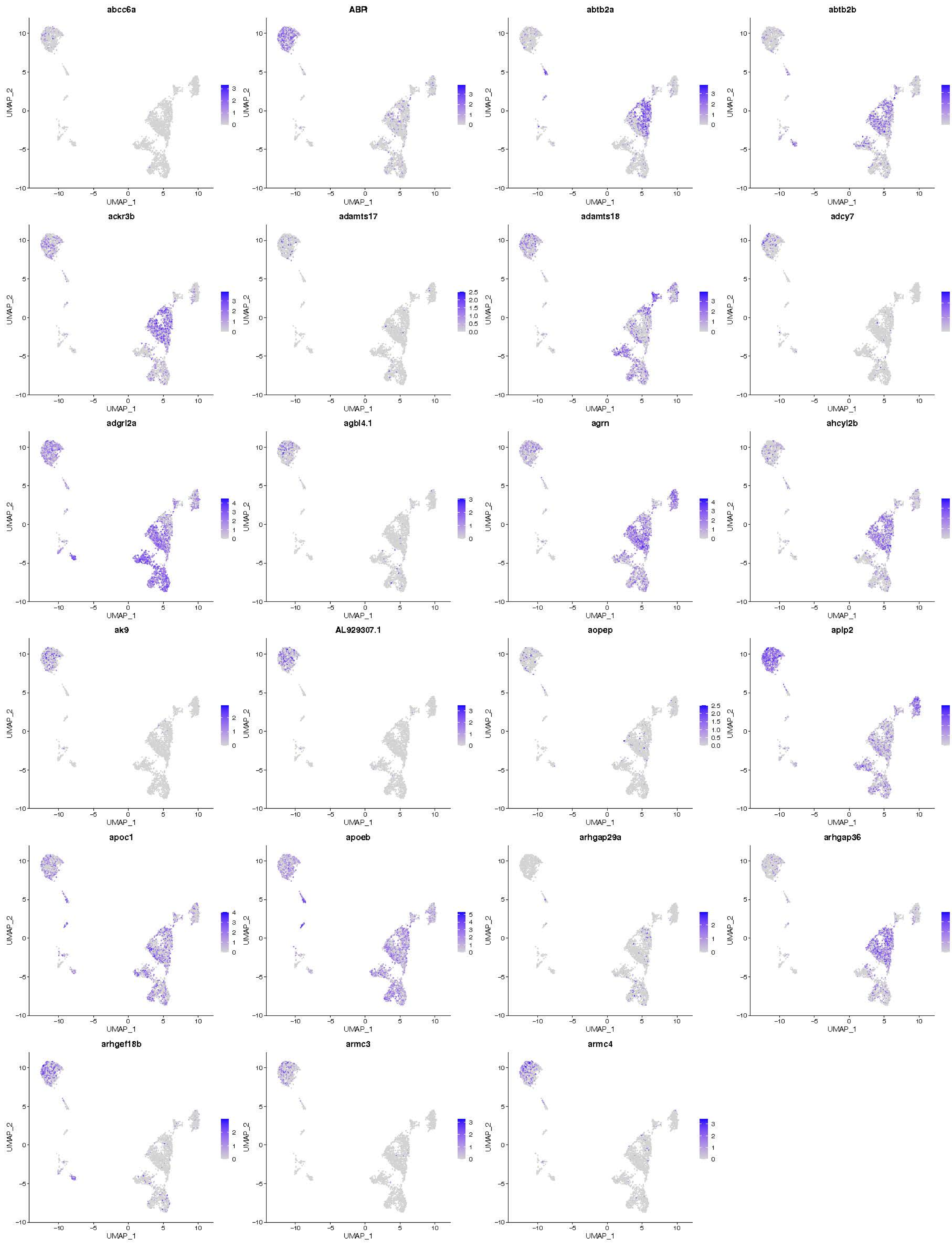

abcc6a
ABR
abtb2a
abtb2b
ackr3b
adamts17
adamts18
adcy7
adgrl2a
agbl4.1
agrn
ahcyl2b
ak9
AL929307.1
aopep
aplp2
apoc1
apoeb
arhgap29a
arhgap36
arhgef18b
armc3
armc4
UMAP_1
UMAP_2

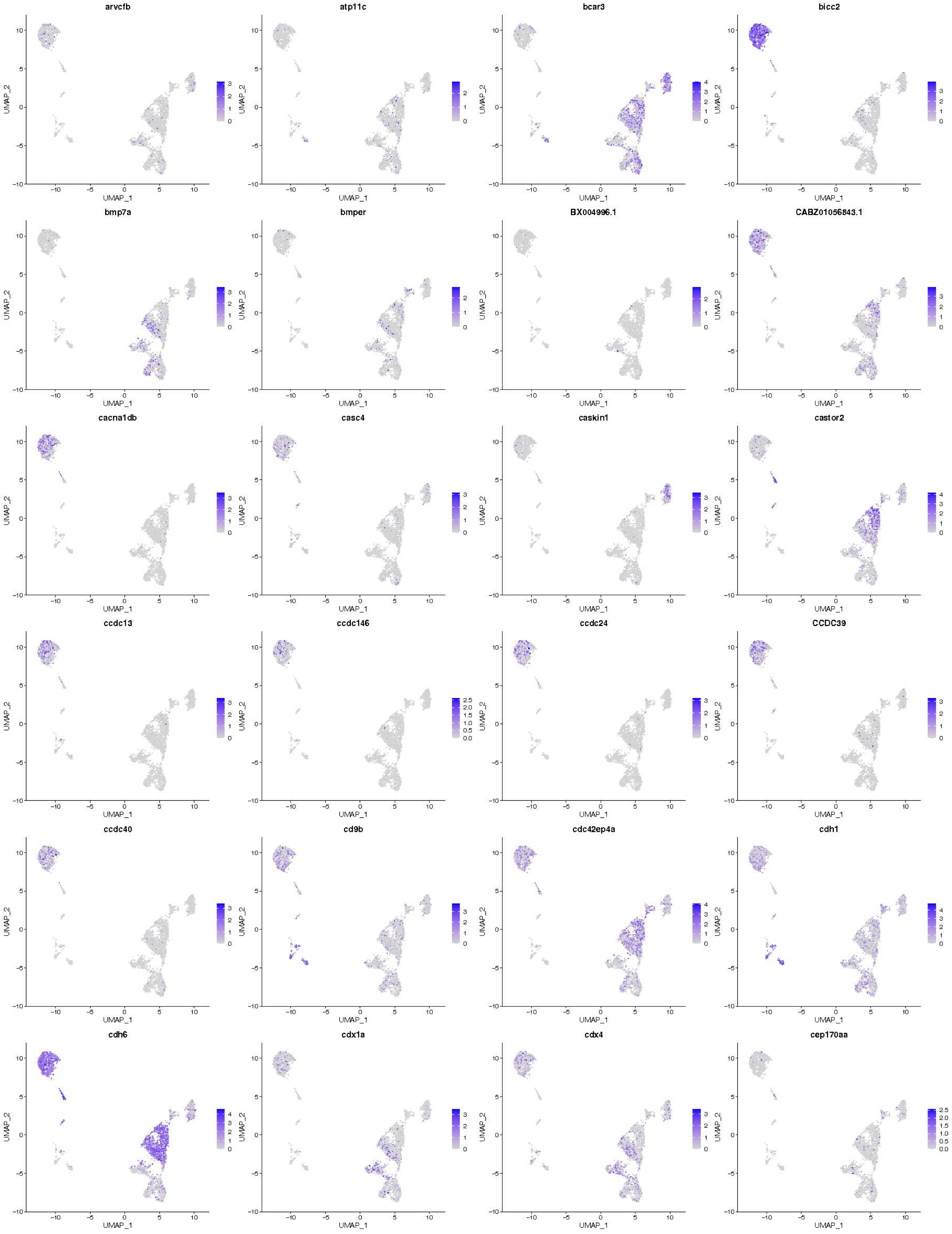
arvcfb
atp11c
bcar3
bicc2
bmp7a
bmper
BX004996.1
CABZ01056843.1
cacna1db
casc4
caskin1
castor2
ccdc13
ccdc146
ccdc24
CCDC39
ccdc40
cd9b
cdc42ep4a
cdh1
cdh6
cdx1a
cdx4
cep170aa
UMAP_1
UMAP_2

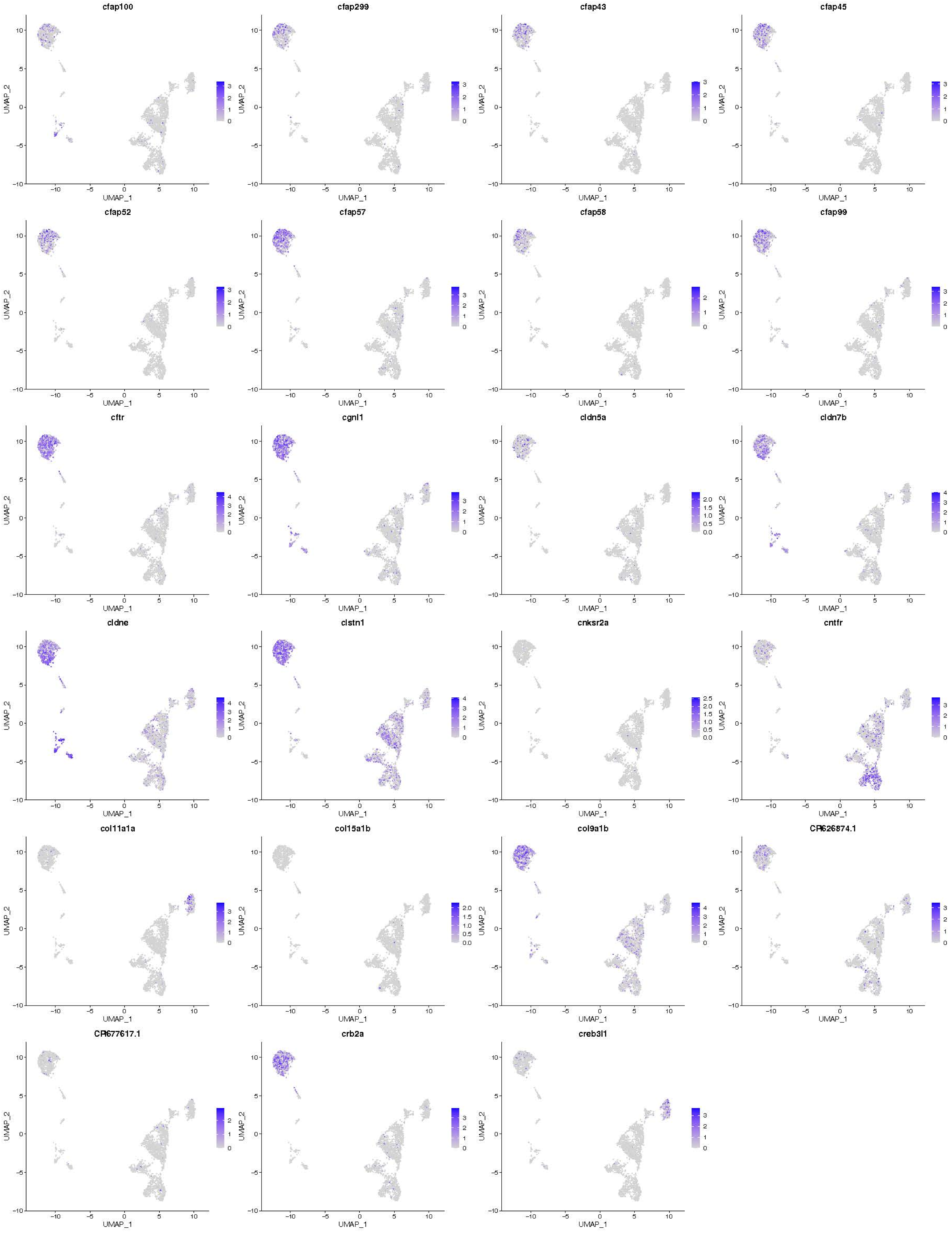
cfap100
cfap299
cfap43
cfap45
cfap52
cfap57
cfap58
cfap99
cftr
cgnl1
cldn5a
cldn7b
cldne
clstn1
cnksr2a
cntfr
col11a1a
col15a1b
col9a1b
CR626874.1
CR677617.1
crb2a
creb3l1
UMAP_1
UMAP_2

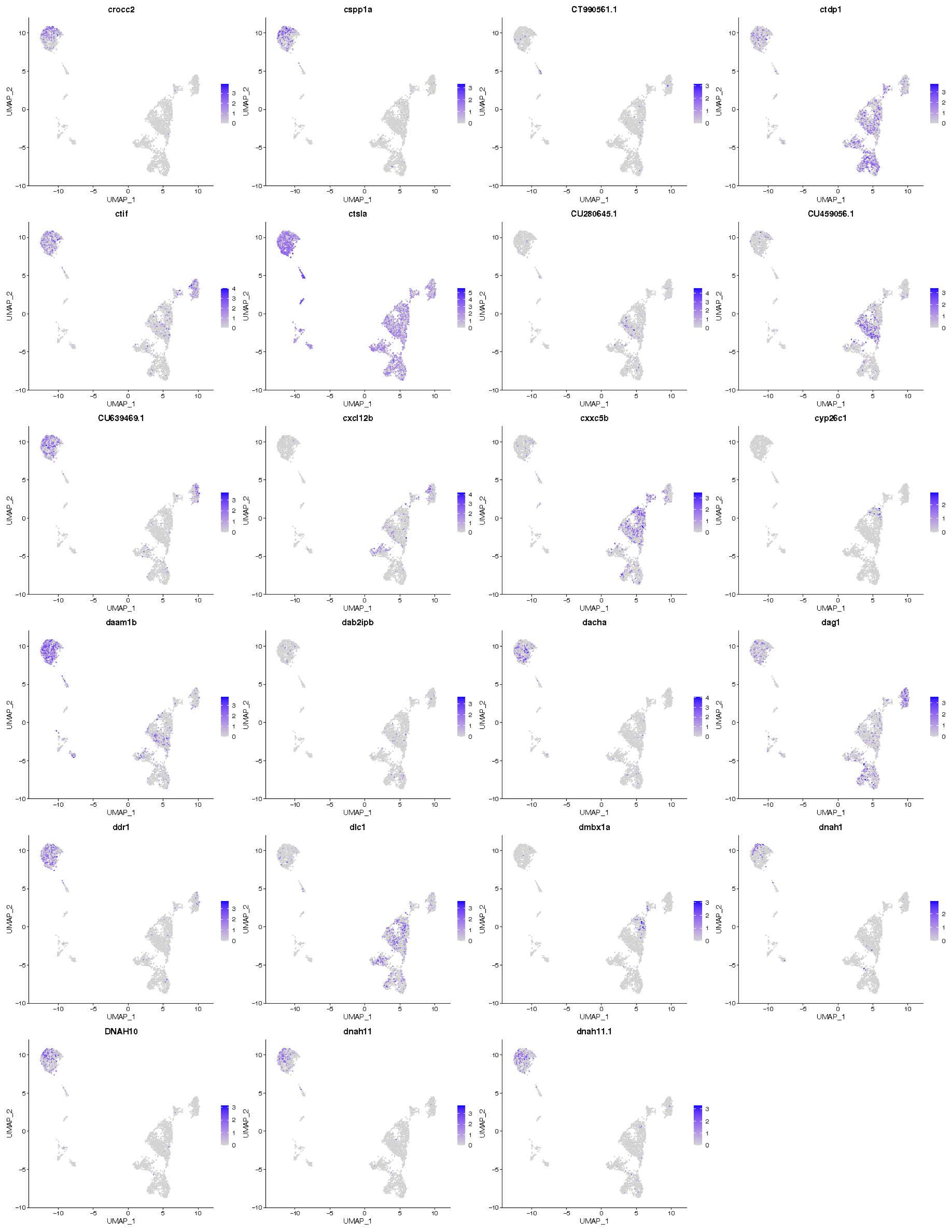
crocc2
cspp1a
CT990561.1
ctdp1
ctif
ctsla
CU280645.1
CU459056.1
CU639469.1
cxcl12b
cxxc5b
cyp26c1
daam1b
dab2ipb
dacha
dag1
ddr1
dlc1
dmbx1a
dnah1
DNAH10
dnah11
dnah11.1
UMAP_1
UMAP_2

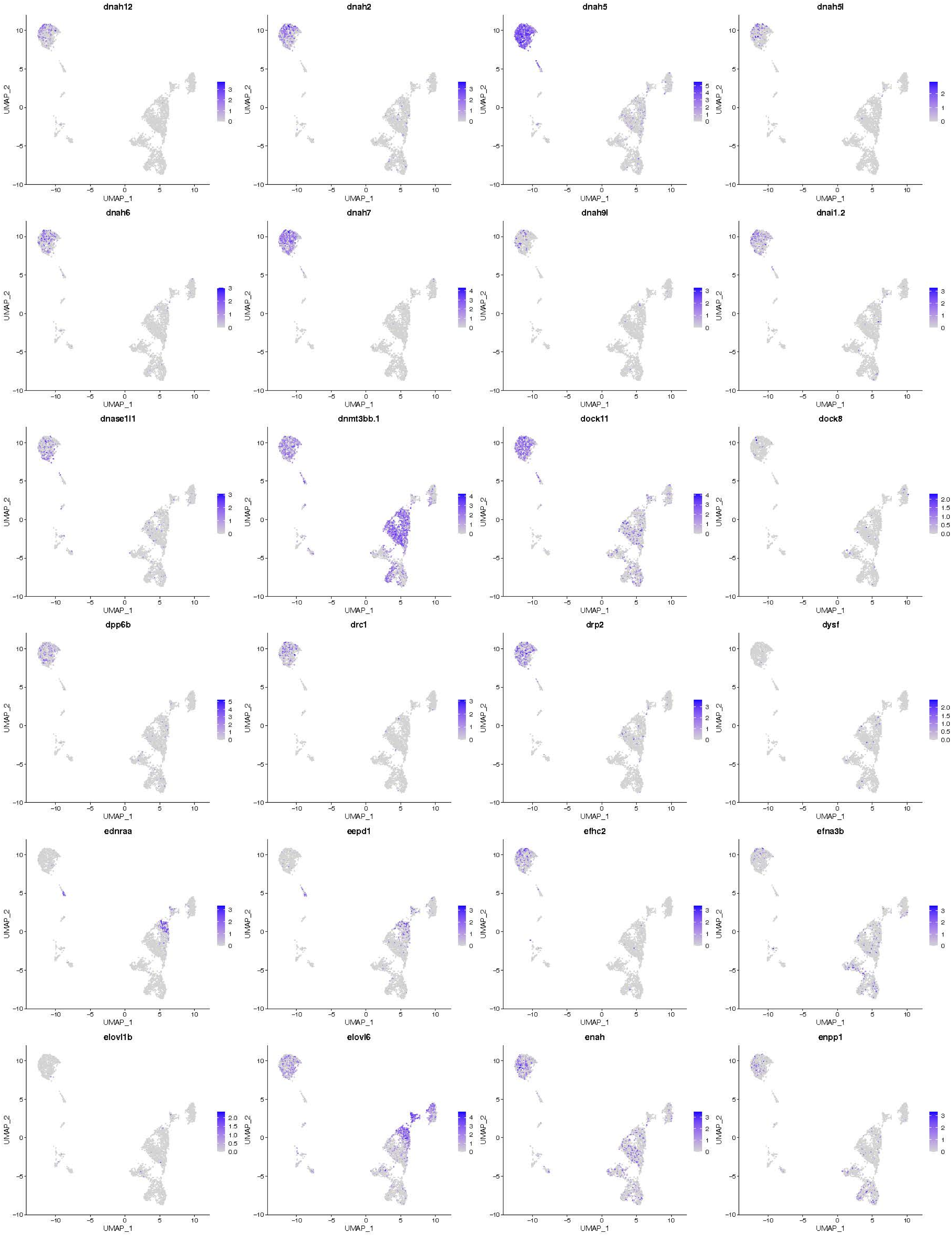
dnah12
dnah2
dnah5
dnah5l
dnah6
dnah7
dnah9l
dnai1.2
dnase1l1
dnmt3bb.1
dock11
dock8
dpp6b
drc1
drp2
dysf
ednraa
eepd1
efhc2
efna3b
elovl1b
elovl6
enah
enpp1
UMAP_1
UMAP_2

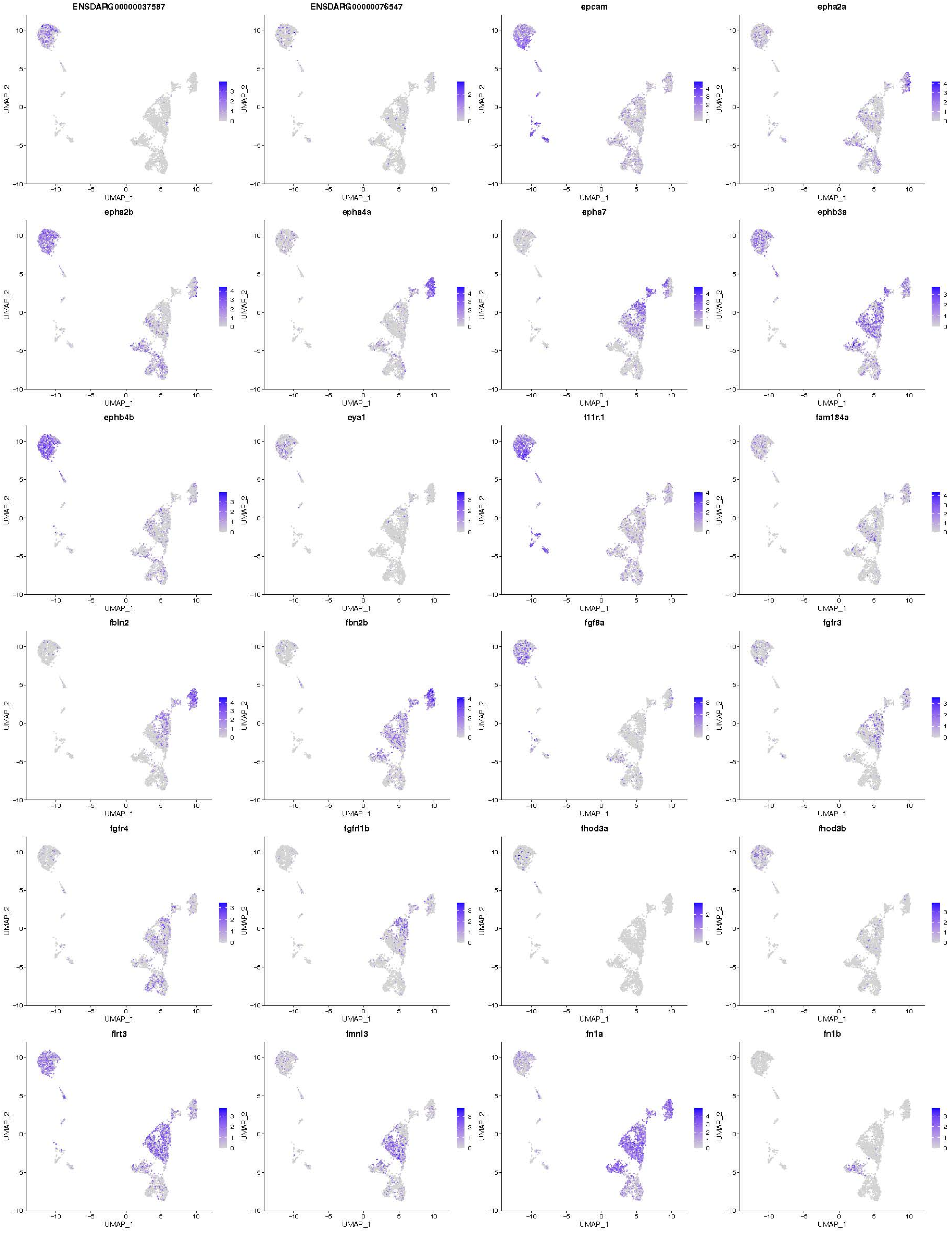
ENSDARG00000037587
ENSDARG00000076547
epcam
epha2a
epha2b
epha4a
epha7
ephb3a
ephb4b
eya1
f11r.1
fam184a
fbln2
fbn2b
fgf8a
fgfr3
fgfr4
fgfrl1b
fhod3a
fhod3b
flrt3
fmnl3
fn1a
fn1b
UMAP_1
UMAP_2

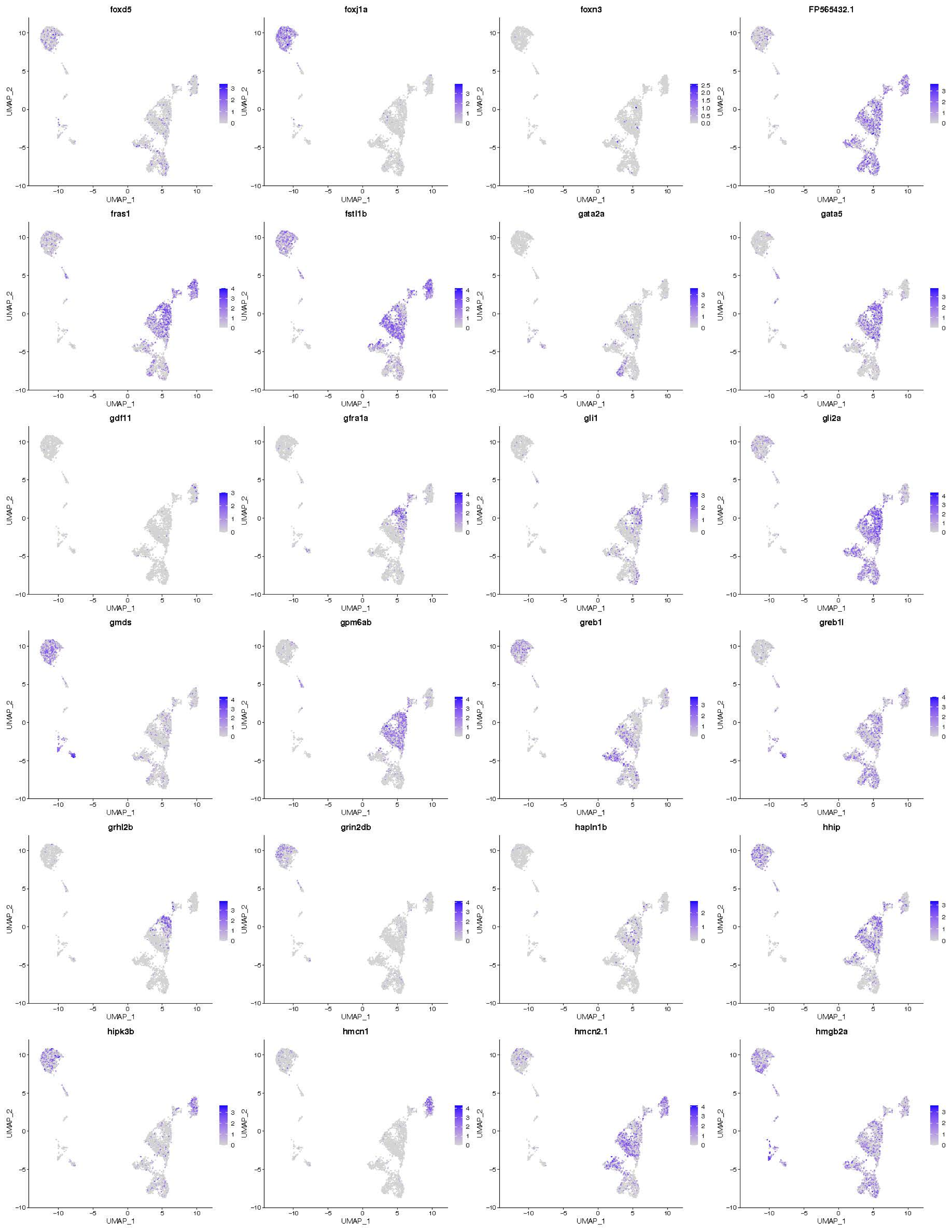
foxd5
foxj1a
foxn3
FP565432.1
fras1
fstl1b
gata2a
gata5
gdf11
gfra1a
gli1
gli2a
gmds
gpm6ab
greb1
greb1l
grhl2b
grin2db
hapln1b
hhip
hipk3b
hmcn1
hmcn2.1
hmgb2a
UMAP_1
UMAP_2

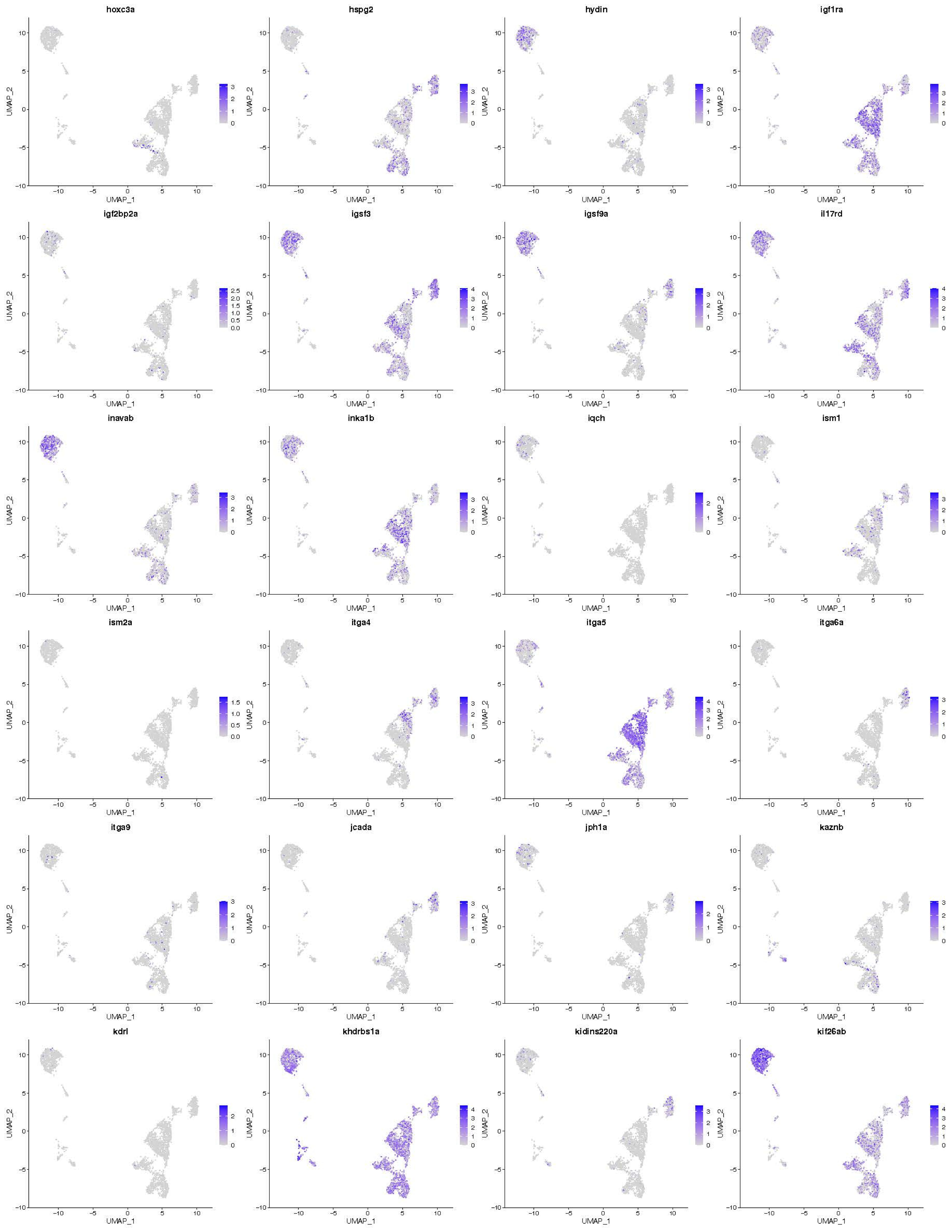

hoxc3a
hspg2
hydin
igf1ra
igf2bp2a
igsf3
igsf9a
il17rd
inavab
inka1b
iqch
ism1
ism2a
itga4
itga5
itga6a
itga9
jcada
jph1a
kaznb
kdrl
khdrbs1a
kidins220a
kif26ab
UMAP_1
UMAP_2

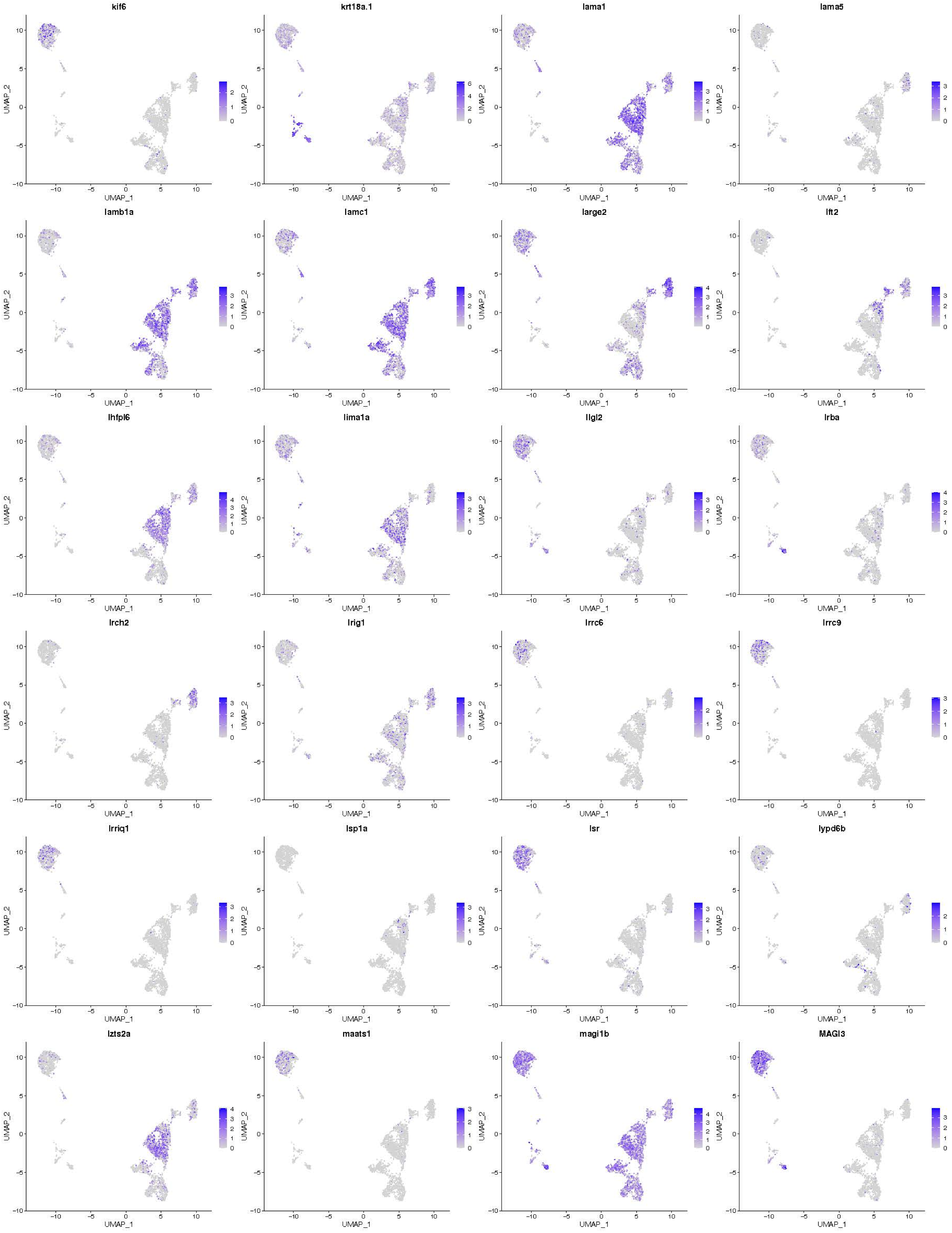
kif6
krt18a.1
lama1
lama5
lamb1a
lamc1
large2
lft2
lhfpl6
lima1a
llgl2
lrba
lrch2
lrig1
lrrc6
lrrc9
lrriq1
lsp1a
lsr
lypd6b
lzts2a
maats1
magi1b
MAGI3
UMAP_1
UMAP_2

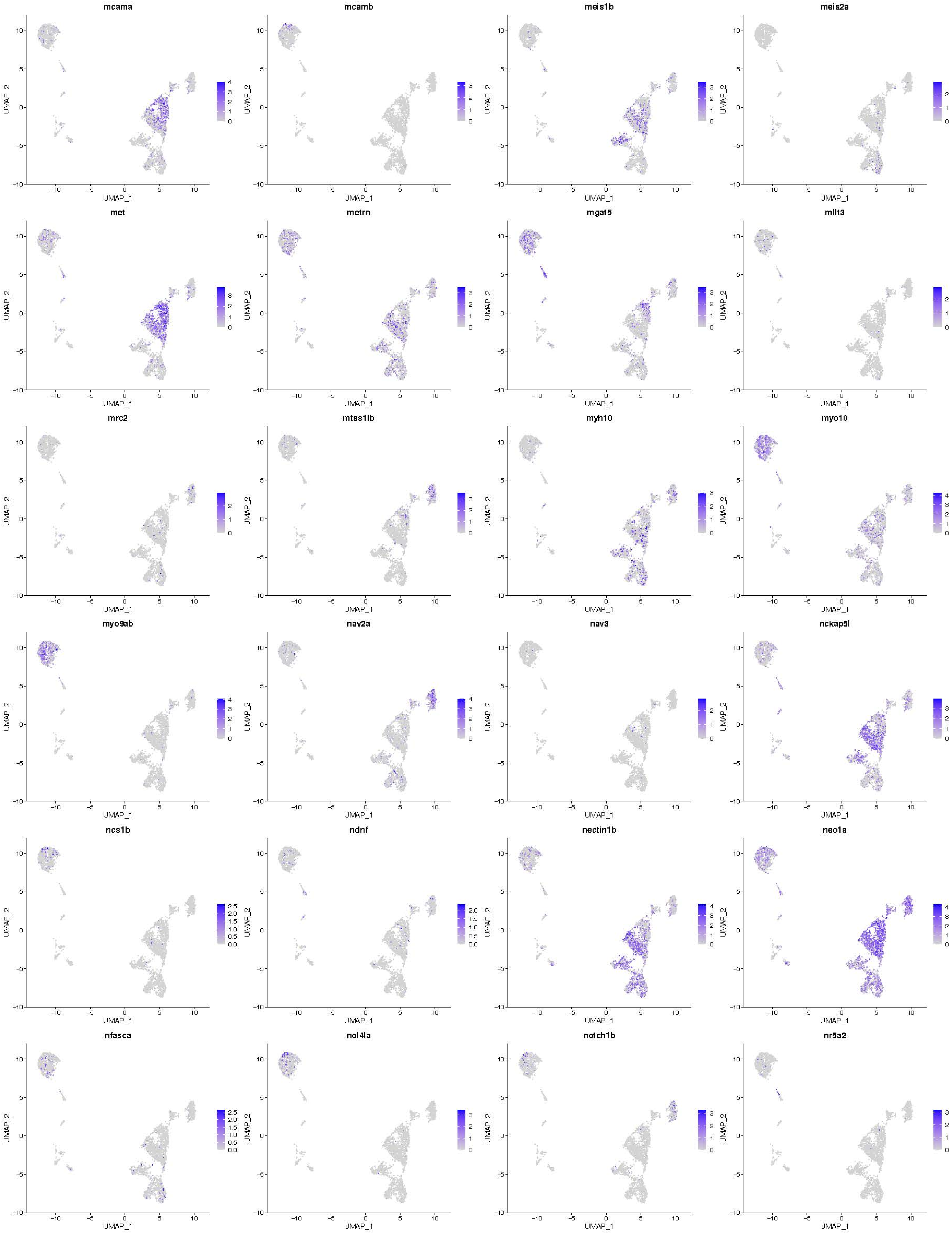
mcama
mcamb
meis1b
meis2a
met
metrn
mgat5
mllt3
mrc2
mtss1lb
myh10
myo10
myo9ab
nav2a
nav3
nckap5l
ncs1b
ndnf
nectin1b
neo1a
nfasca
nol4la
notch1b
nr5a2
UMAP_1
UMAP_2

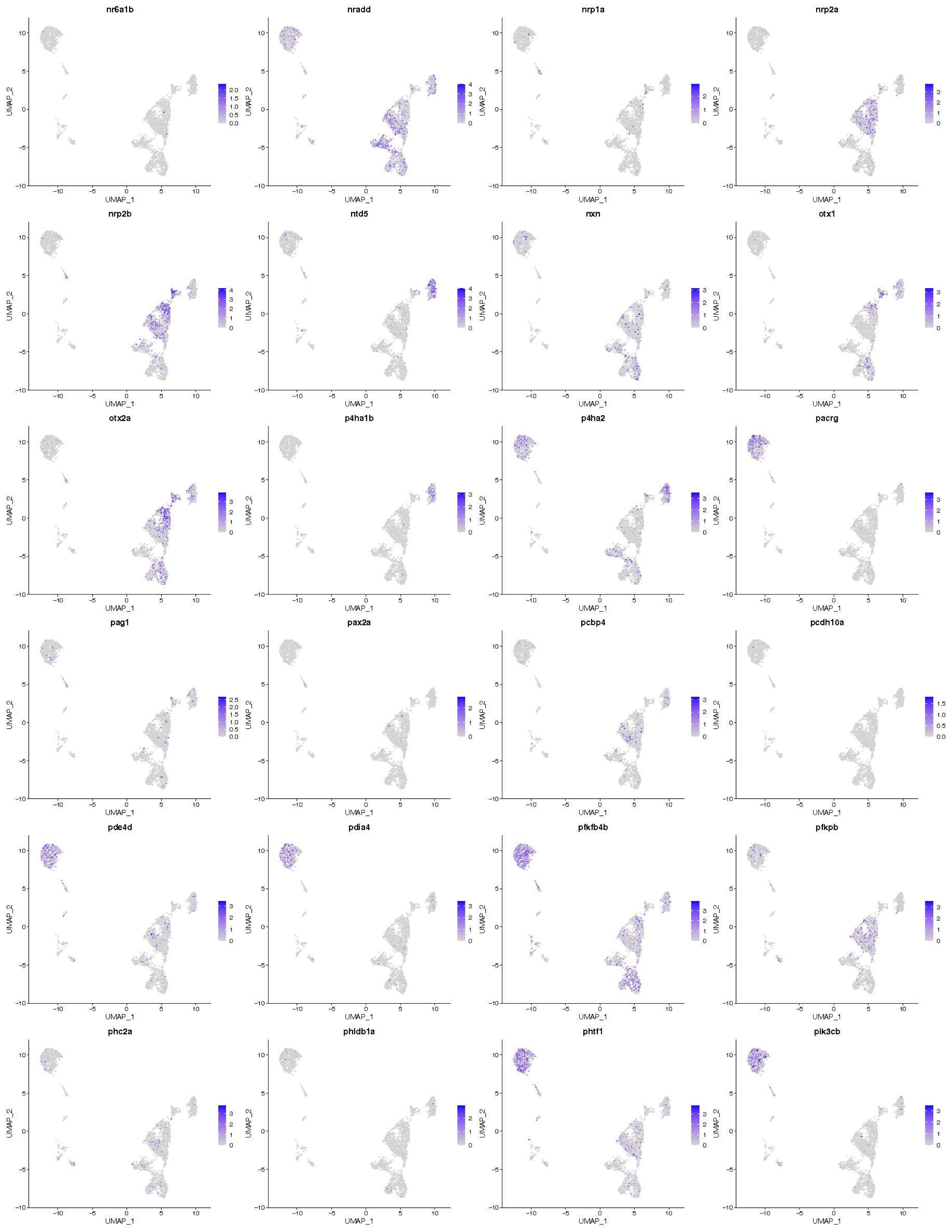
nr6a1b
nradd
nrp1a
nrp2a
nrp2b
ntd5
nxn
otx1
otx2a
p4ha1b
p4ha2
pacrg
pag1
pax2a
pcbp4
pcdh10a
pde4d
pdia4
pfkfb4b
pfkpb
phc2a
phldb1a
phtf1
pik3cb
UMAP_1
UMAP_2

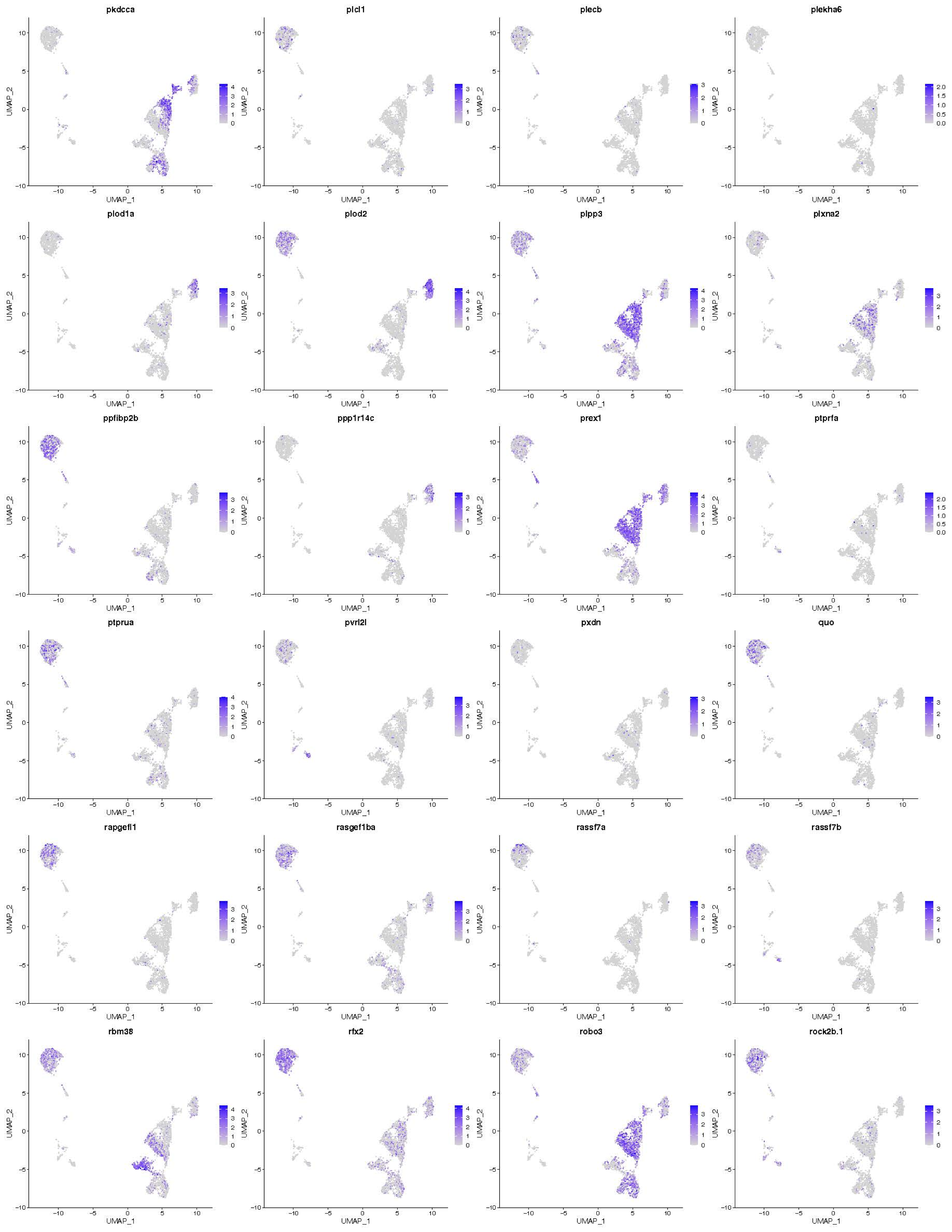
pkdcca
plcl1
plecb
plekha6
plod1a
plod2
plpp3
plxna2
ppfibp2b
ppp1r14c
prex1
ptprfa
ptprua
pvrl2l
pxdn
quo
rapgefl1
rasgef1ba
rassf7a
rassf7b
rbm38
rfx2
robo3
rock2b.1
UMAP_1
UMAP_2

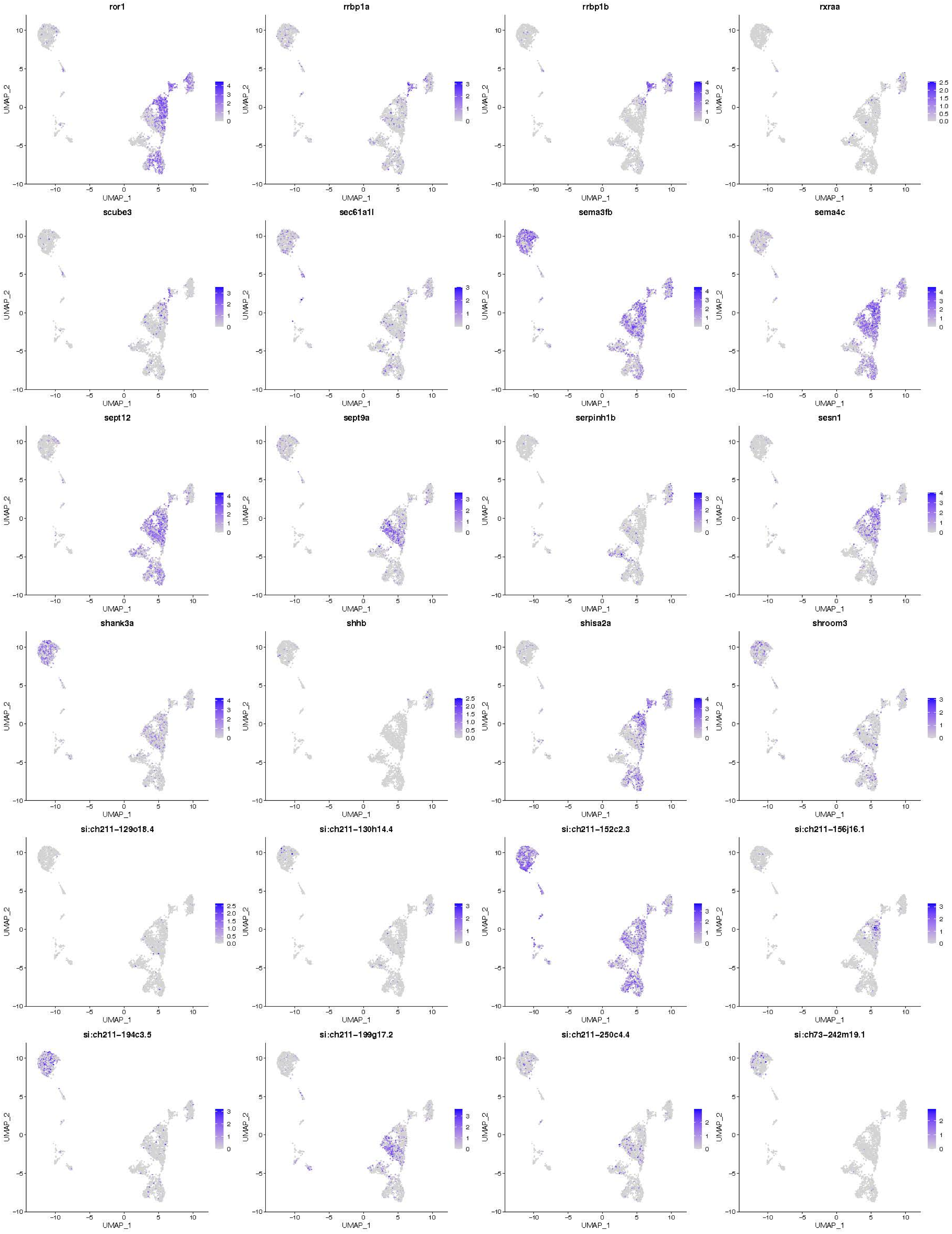

ror1
rrbp1a
rrbp1b
rxraa
scube3
sec61a1l
sema3fb
sema4c
sept12
sept9a
serpinh1b
sesn1
shank3a
shhb
shisa2a
shroom3
si:ch211-129o18.4
si:ch211-130h14.4
si:ch211-152c2.3
si:ch211-156j16.1
si:ch211-194c3.5
si:ch211-199g17.2
si:ch211-250c4.4
si:ch73-242m19.1
UMAP_1
UMAP_2

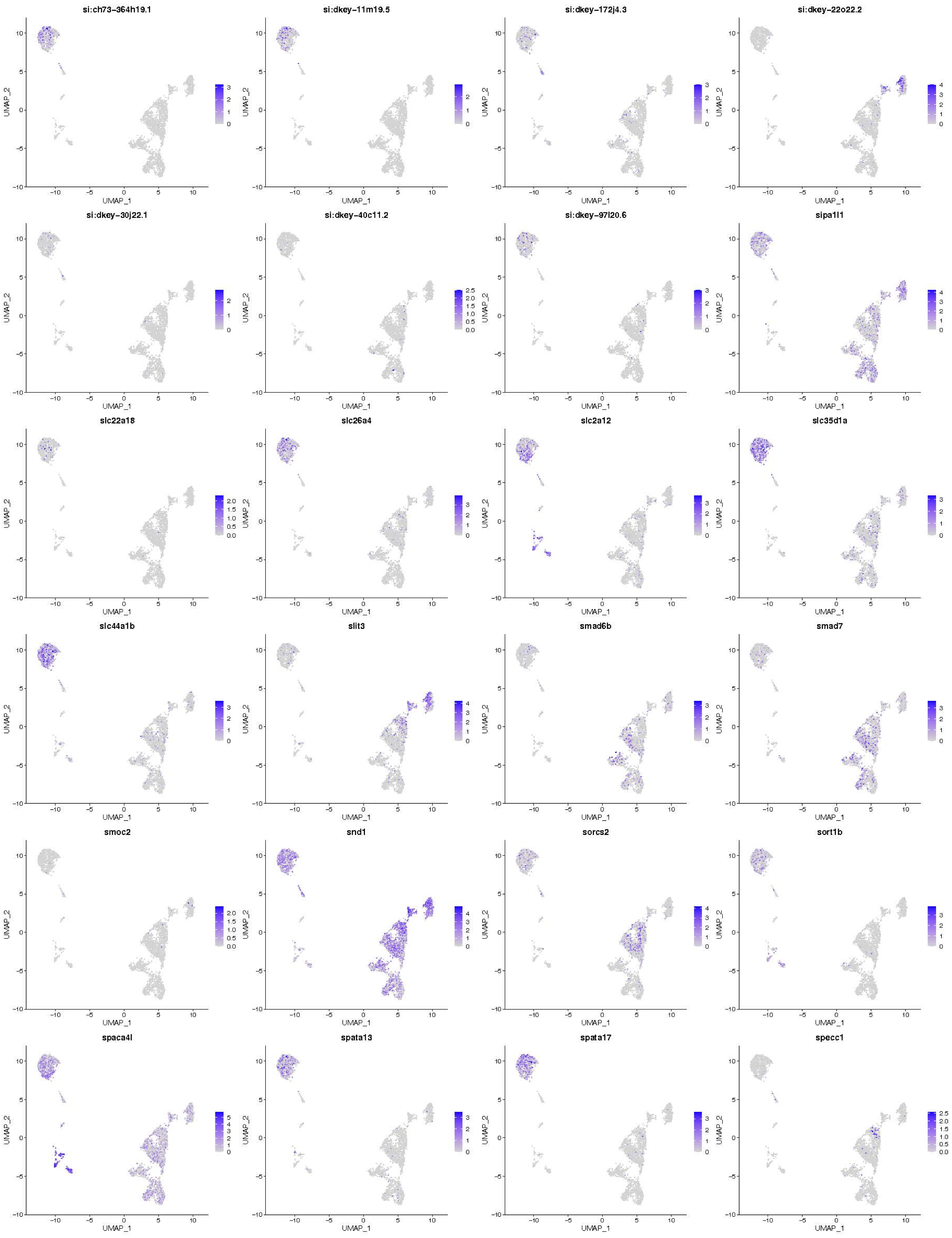
si:ch73-364h19.1
si:dkey-11m19.5
si:dkey-172j4.3
si:dkey-22o22.2
si:dkey-30j22.1
si:dkey-40c11.2
si:dkey-97l20.6
sipa1l1
slc22a18
slc26a4
slc2a12
slc35d1a
slc44a1b
slit3
smad6b
smad7
smoc2
snd1
sorcs2
sort1b
spaca4l
spata13
spata17
specc1
UMAP_1
UMAP_2

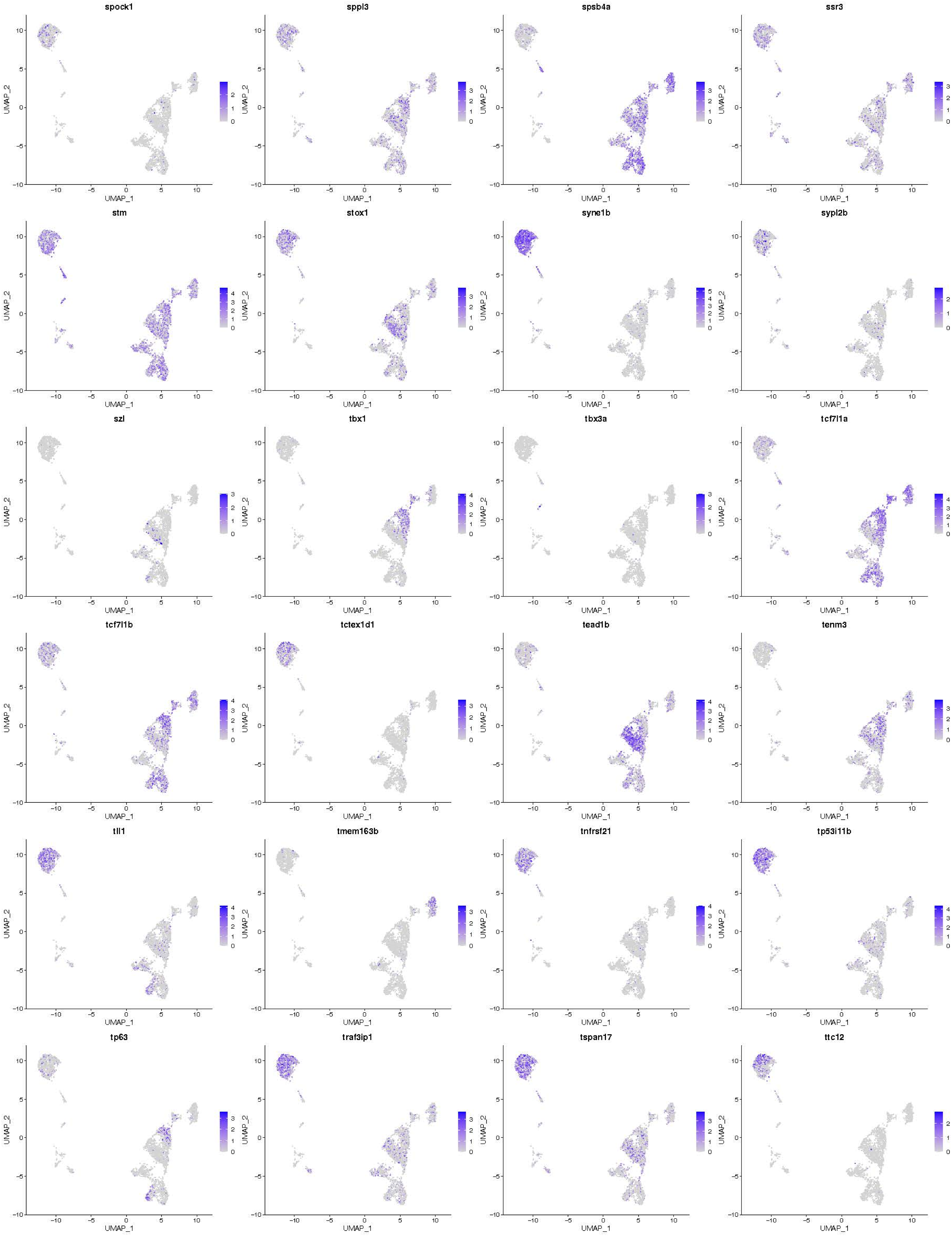
spock1
sppl3
spsb4a
ssr3
stm
stox1
syne1b
sypl2b
szl
tbx1
tbx3a
tcf7l1a
tcf7l1b
tctex1d1
tead1b
tenm3
tll1
tmem163b
tnfrsf21
tp53i11b
tp63
traf3ip1
tspan17
ttc12
UMAP_1
UMAP_2

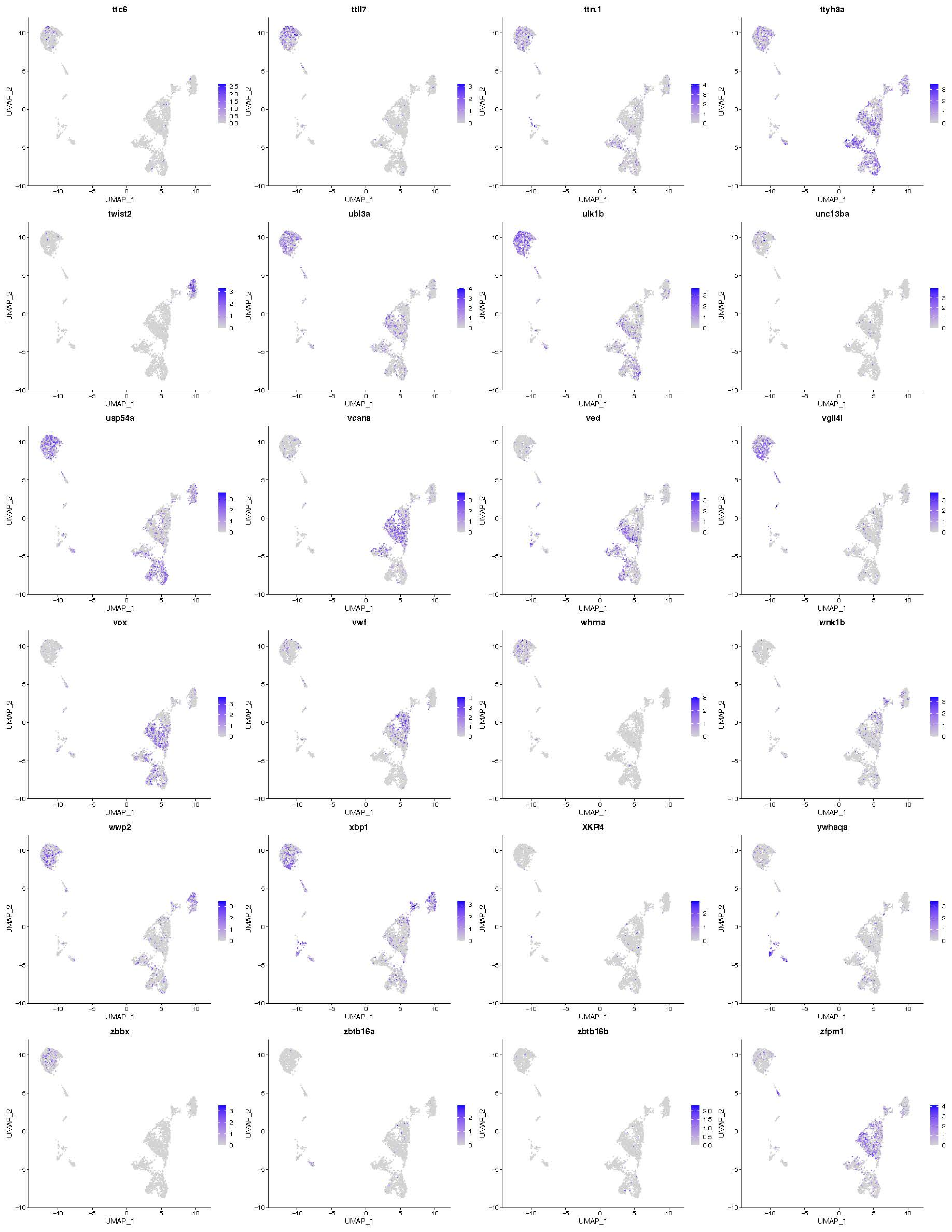

ttc6
ttll7
ttn.1
ttyh3a
twist2
ubl3a
ulk1b
unc13ba
usp54a
vcana
ved
vgll4l
vox
vwf
whrna
wnk1b
wwp2
xbp1
XKR4
ywhaqa
zbbx
zbtb16a
zbtb16b
zfpm1
UMAP_1
UMAP_2

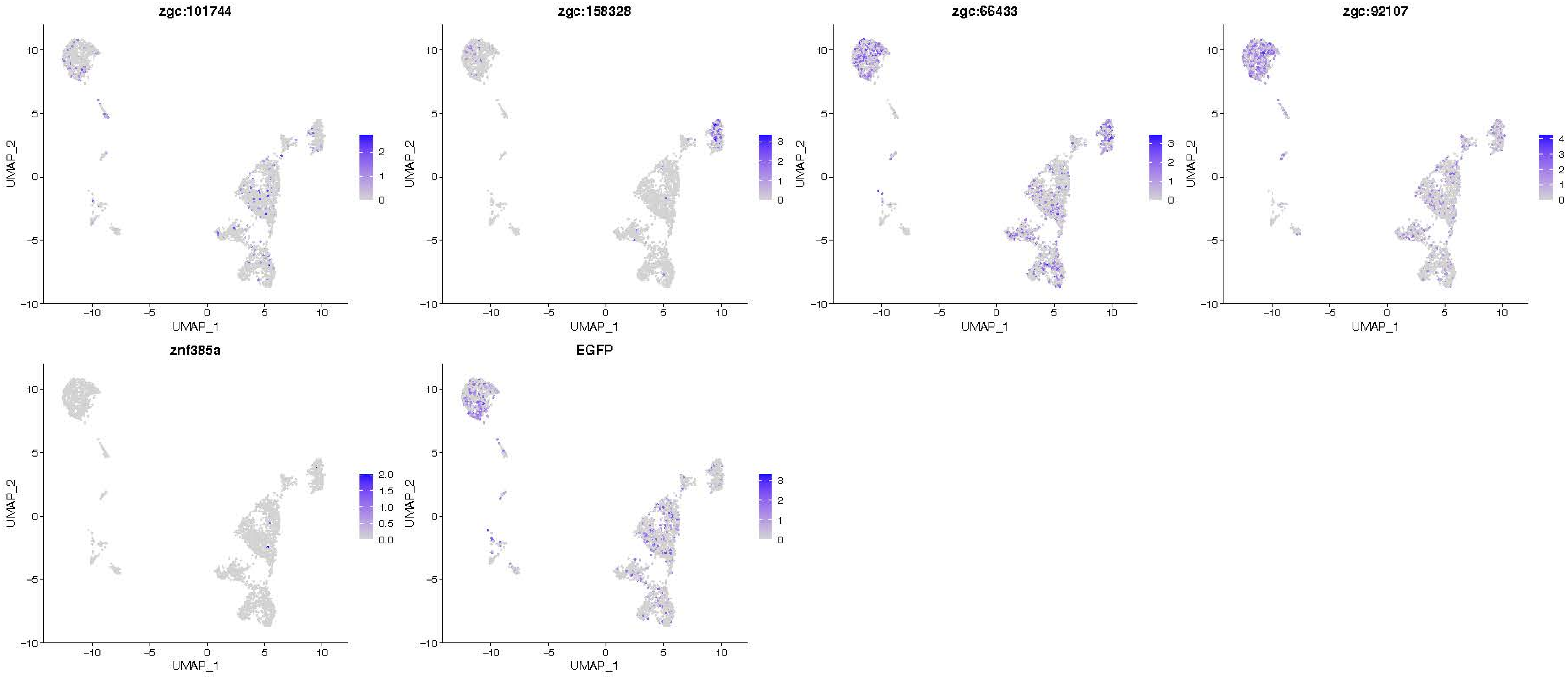
zgc:101744
zgc:158328
zgc:66433
zgc:92107
znf385a
EGFP
UMAP_1
UMAP_2

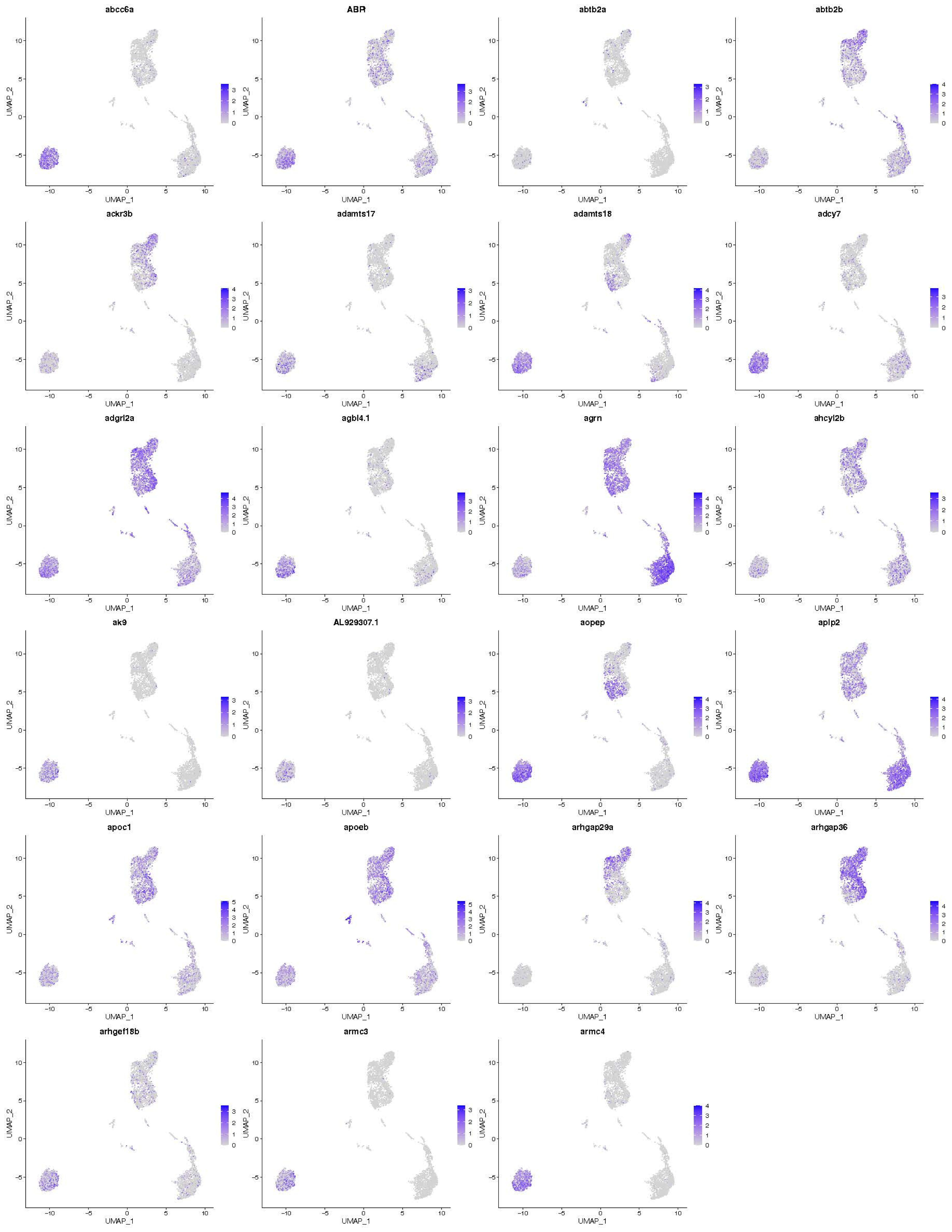
abcc6a
ABR
abtb2a
abtb2b
ackr3b
adamts17
adamts18
adcy7
adgrl2a
agbl4.1
agrn
ahcyl2b
ak9
AL929307.1
aopep
aplp2
apoc1
apoeb
arhgap29a
arhgap36
arhgef18b
armc3
armc4
UMAP_1
UMAP_2

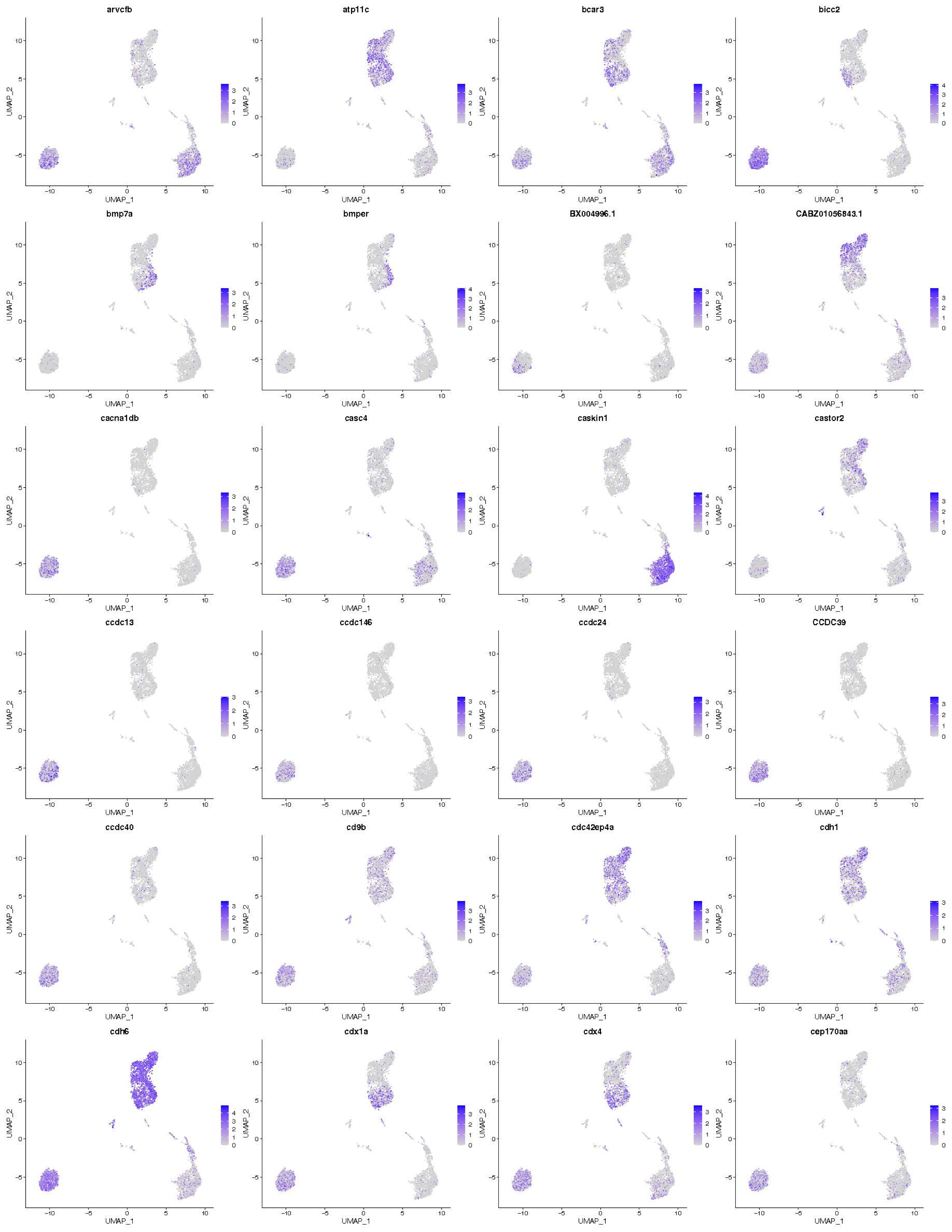
arvcfb
atp11c
bcar3
bicc2
bmp7a
bmper
BX004996.1
CABZ01056843.1
cacna1db
casc4
caskin1
castor2
ccdc13
ccdc146
ccdc24
CCDC39
ccdc40
cd9b
cdc42ep4a
cdh1
cdh6
cdx1a
cdx4
cep170aa
UMAP_1
UMAP_2

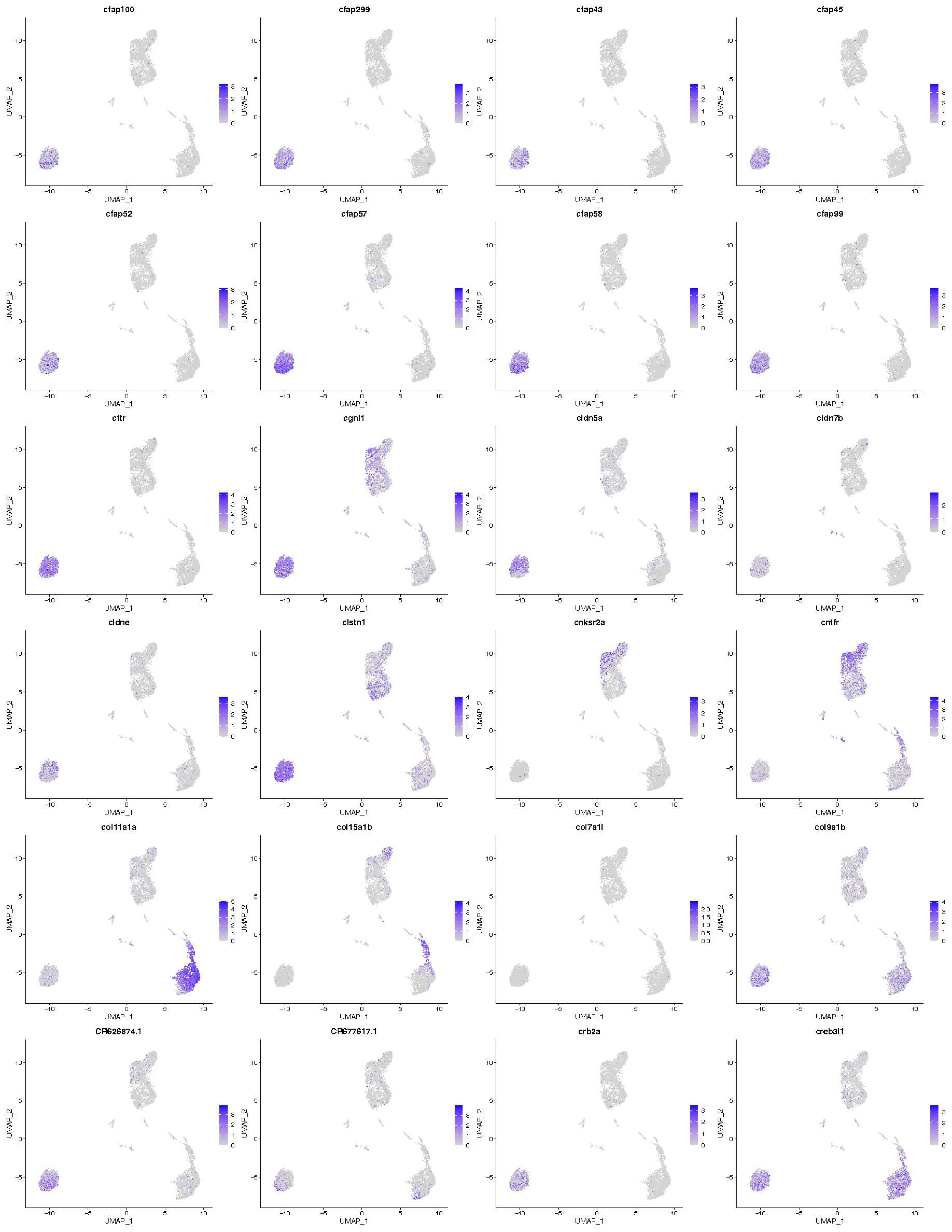

cfap100
cfap299
cfap43
cfap45
cfap52
cfap57
cfap58
cfap99
cftr
cgnl1
cldn5a
cldn7b
cldne
clstn1
cnksr2a
cntfr
col11a1a
col15a1b
col7a1l
col9a1b
CR626874.1
CR677617.1
crb2a
creb3l1
UMAP_1
UMAP_2

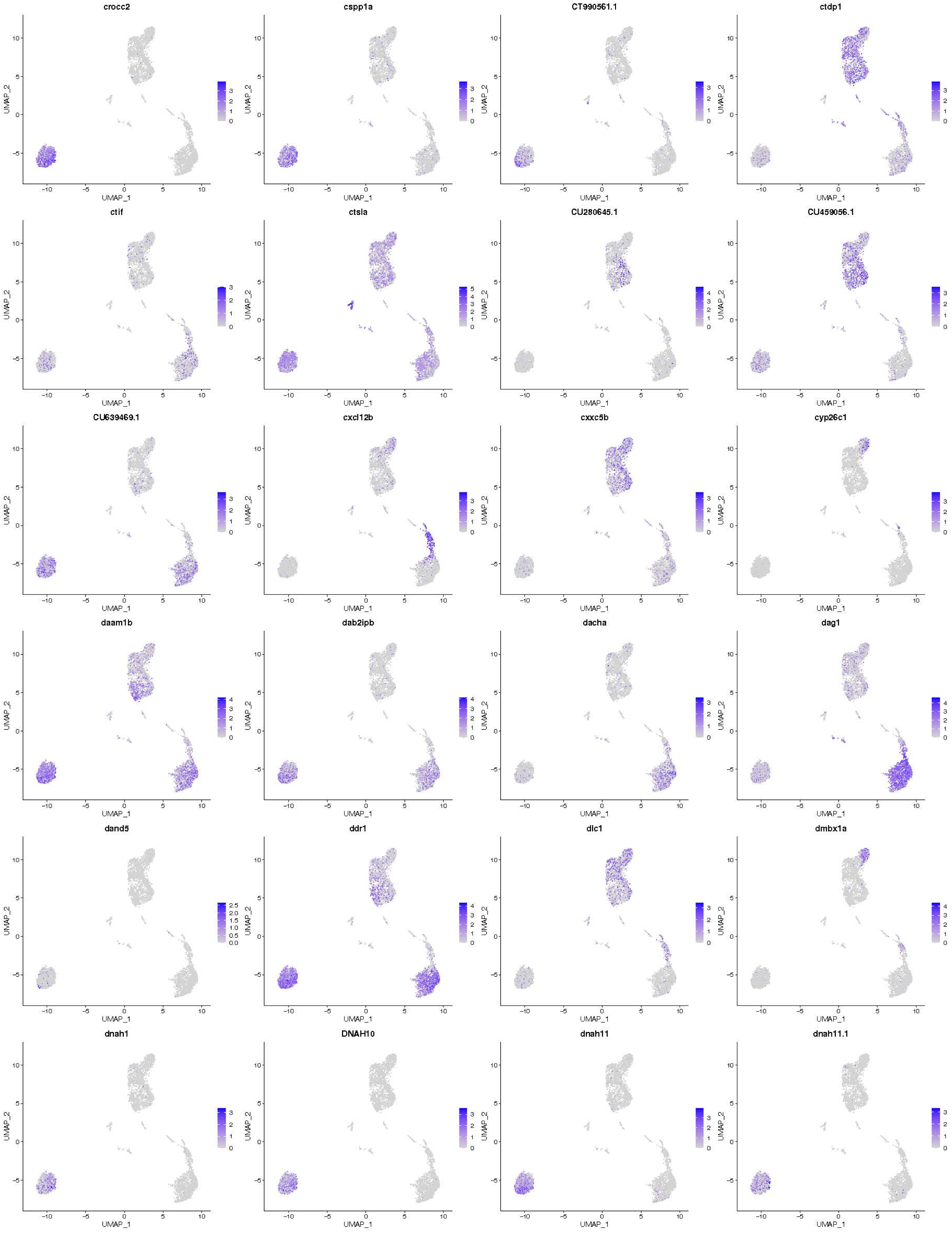

crocc2
cspp1a
CT990561.1
ctdp1
ctif
ctsla
CU280645.1
CU459056.1
CU639469.1
cxcl12b
cxxc5b
cyp26c1
daam1b
dab2ipb
dacha
dag1
dand5
ddr1
dlc1
dmbx1a
dnah1
DNAH10
dnah11
dnah11.1
UMAP_1
UMAP_2

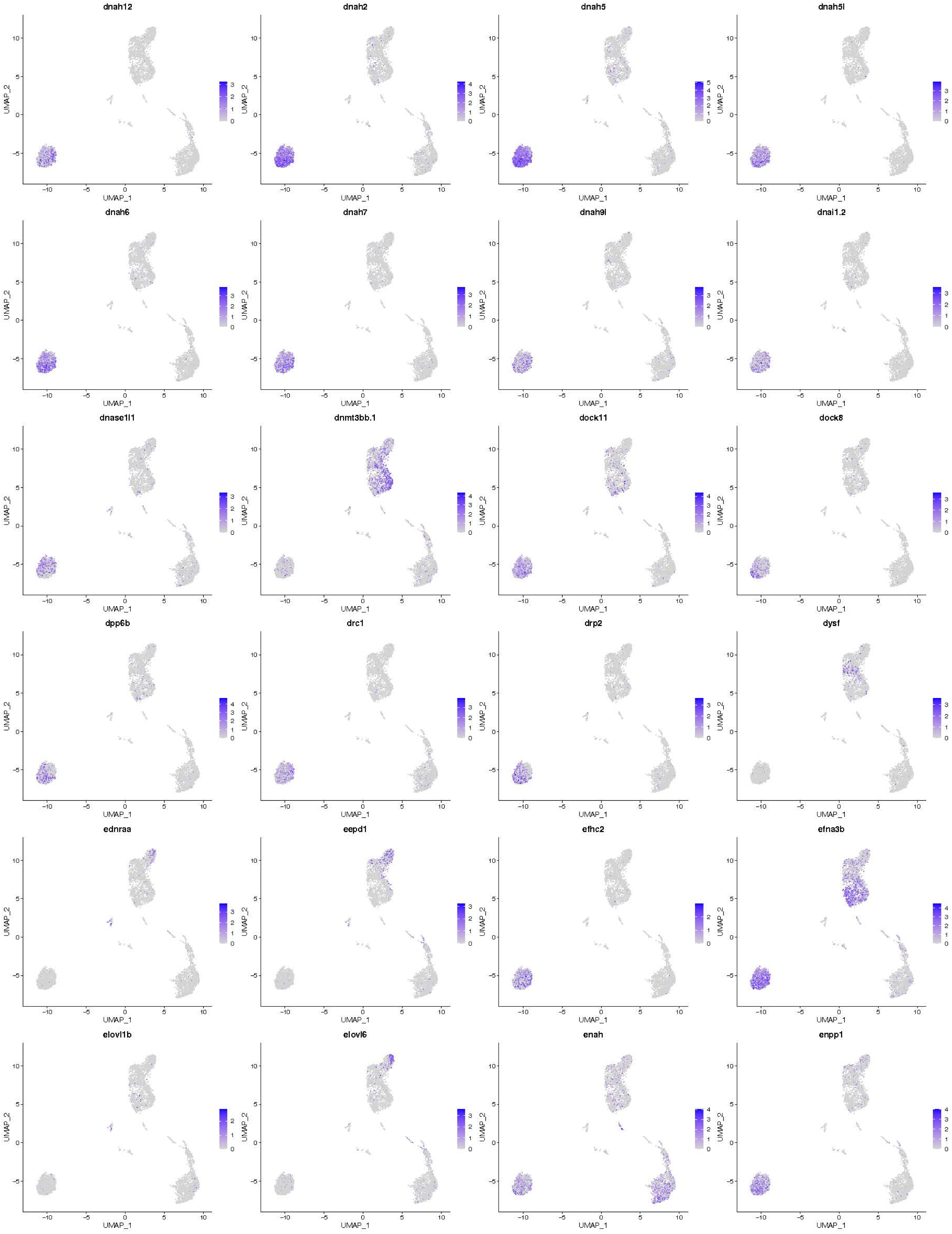
dnah12
dnah2
dnah5
dnah5l
dnah6
dnah7
dnah9l
dnai1.2
dnase1l1
dnmt3bb.1
dock11
dock8
dpp6b
drc1
drp2
dysf
ednraa
eepd1
efhc2
efna3b
elovl1b
elovl6
enah
enpp1
UMAP_1
UMAP_2

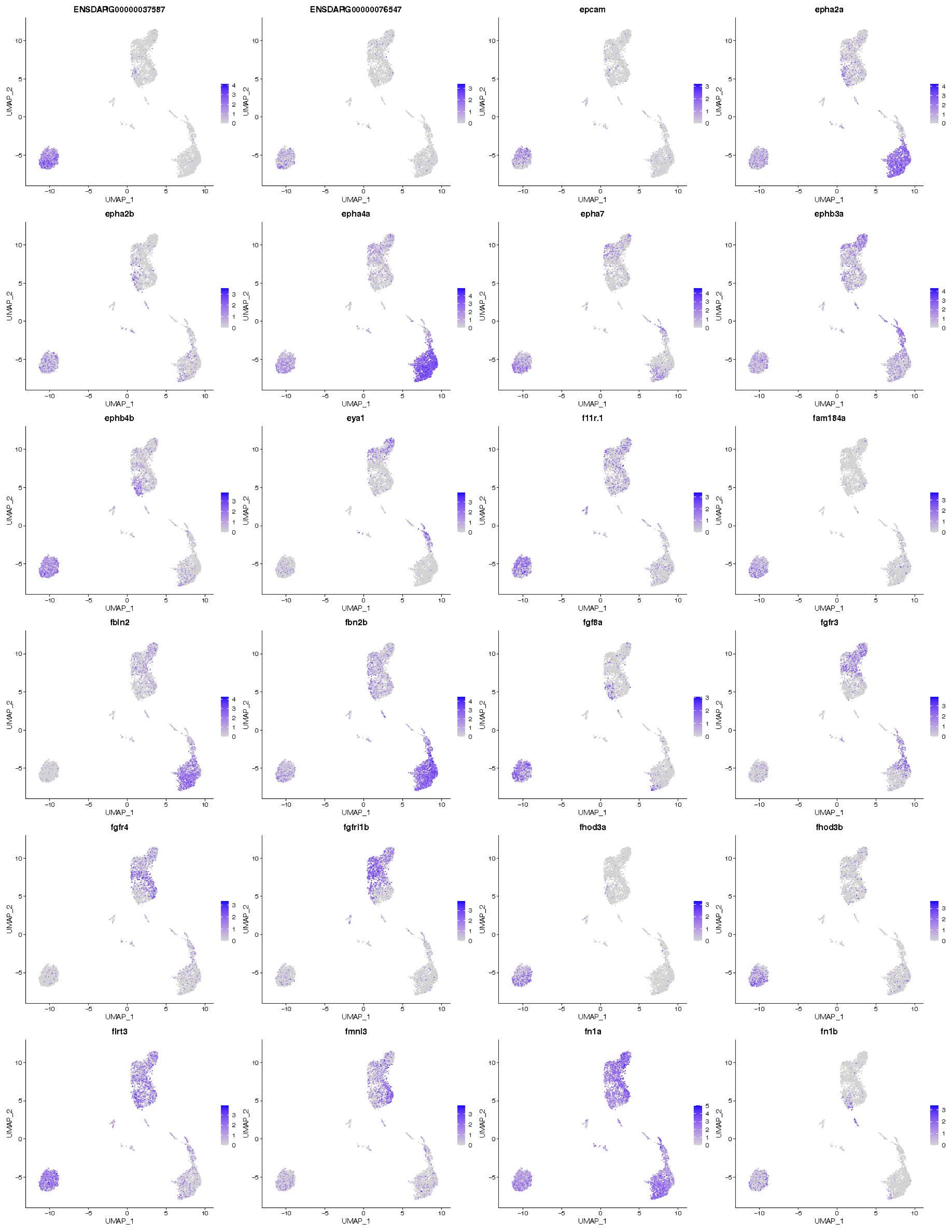
ENSDARG00000037587
ENSDARG00000076547
epcam
epha2a
epha2b
epha4a
epha7
ephb3a
ephb4b
eya1
f11r.1
fam184a
fbln2
fbn2b
fgf8a
fgfr3
fgfr4
fgfrl1b
fhod3a
fhod3b
flrt3
fmnl3
fn1a
fn1b
UMAP_1
UMAP_2

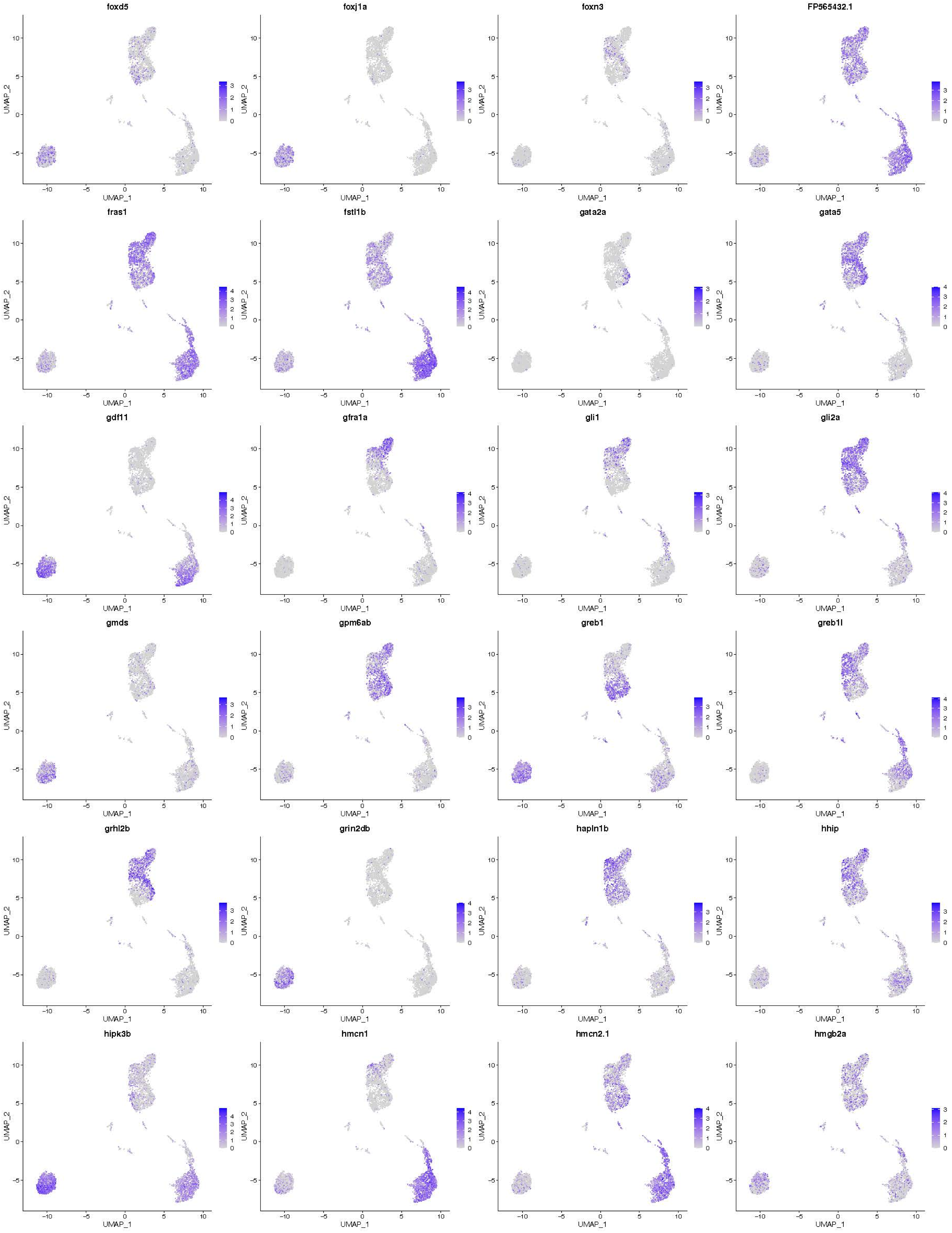
foxd5
foxj1a
foxn3
FP565432.1
fras1
fstl1b
gata2a
gata5
gdf11
gfra1a
gli1
gli2a
gmds
gpm6ab
greb1
greb1l
grhl2b
grin2db
hapln1b
hhip
hipk3b
hmcn1
hmcn2.1
hmgb2a
UMAP_1
UMAP_2

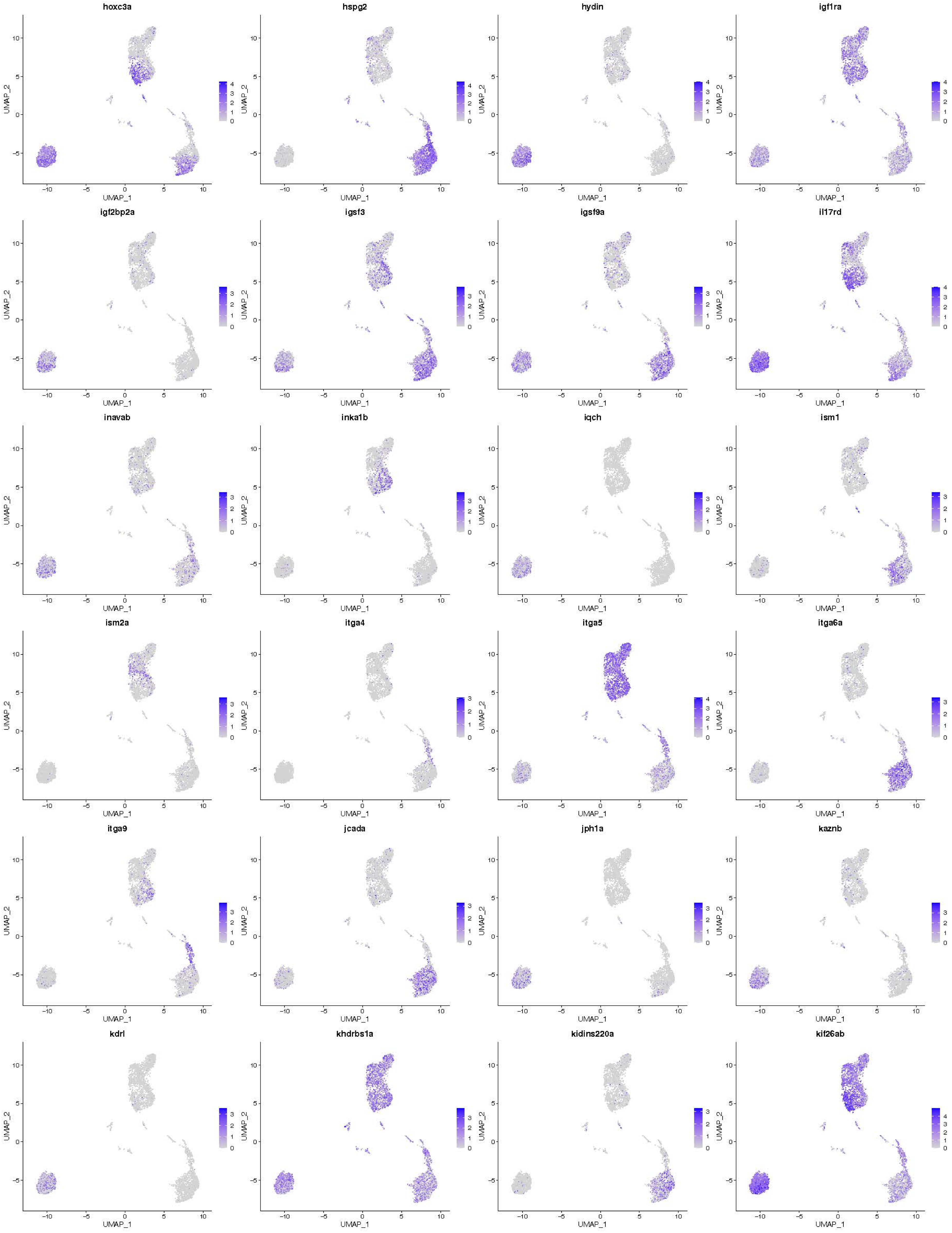
hoxc3a
hspg2
hydin
igf1ra
igf2bp2a
igsf3
igsf9a
il17rd
inavab
inka1b
iqch
ism1
ism2a
itga4
itga5
itga6a
itga9
jcada
jph1a
kaznb
kdrl
khdrbs1a
kidins220a
kif26ab
UMAP_1
UMAP_2

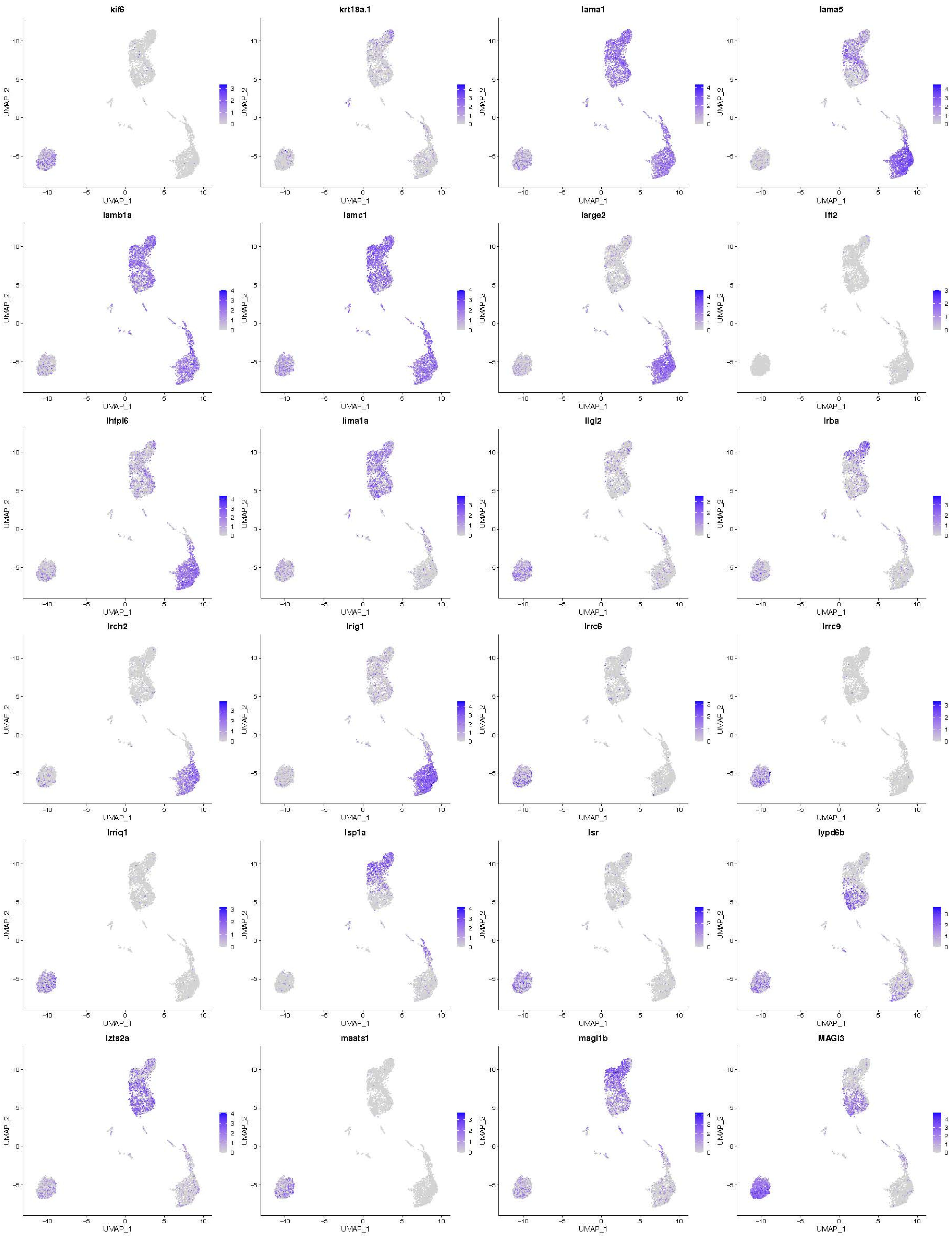
kif6
krt18a.1
lama1
lama5
lamb1a
lamc1
large2
lft2
lhfpl6
lima1a
llgl2
lrba
lrch2
lrig1
lrrc6
lrrc9
lrriq1
lsp1a
lsr
lypd6b
lzts2a
maats1
magi1b
MAGI3
UMAP_1
UMAP_2

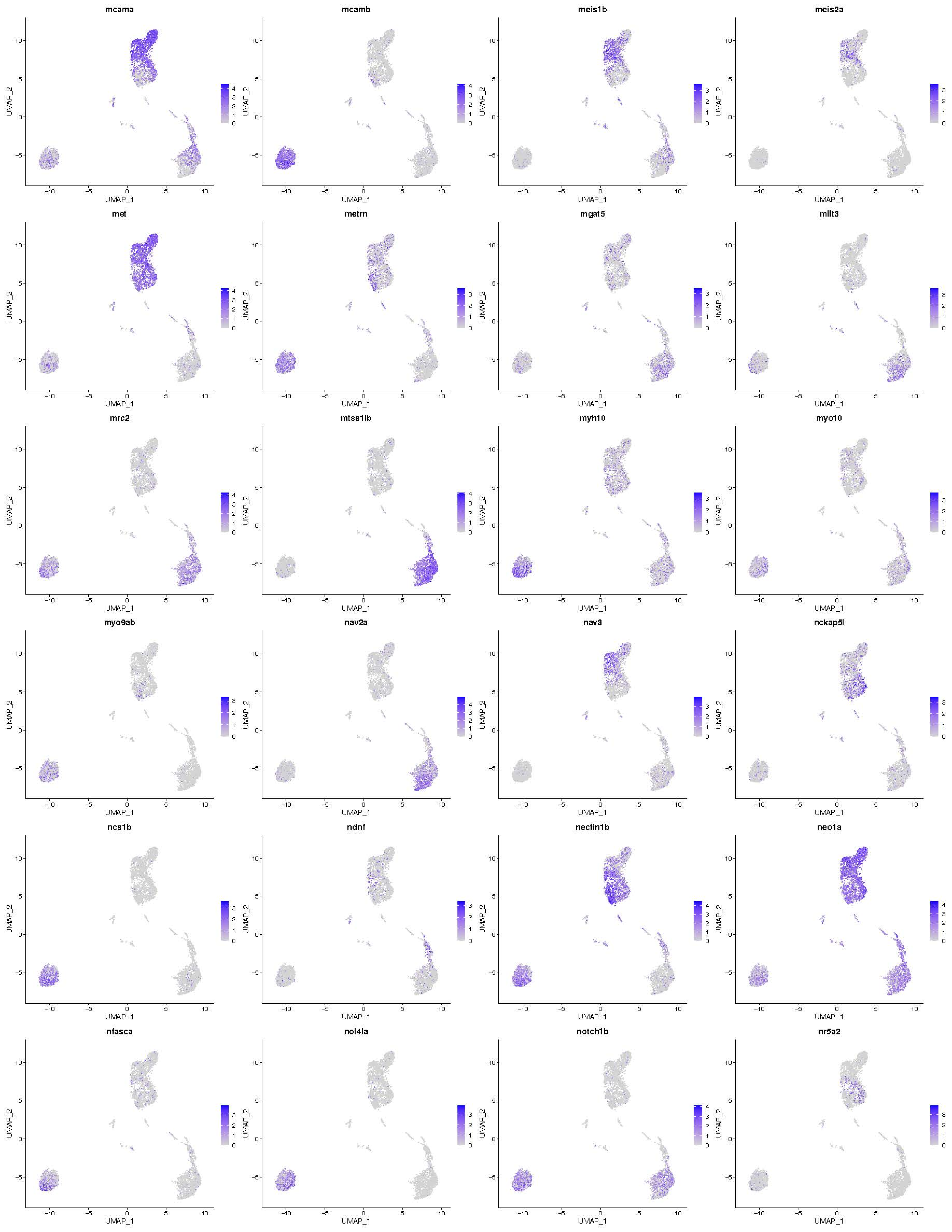
mcama
mcamb
meis1b
meis2a
met
metrn
mgat5
mllt3
mrc2
mtss1lb
myh10
myo10
myo9ab
nav2a
nav3
nckap5l
ncs1b
ndnf
nectin1b
neo1a
nfasca
nol4la
notch1b
nr5a2
UMAP_1
UMAP_2

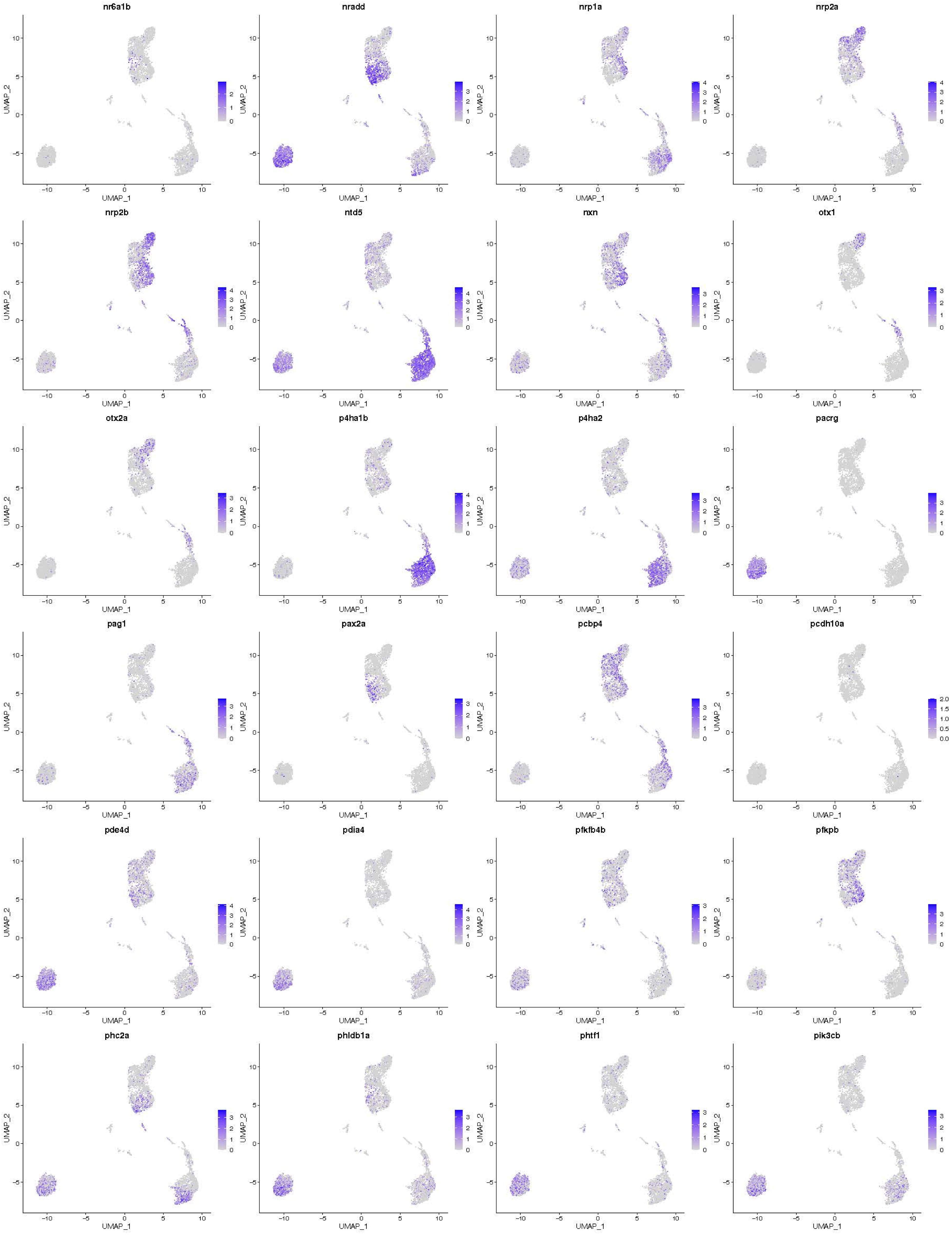
nr6a1b
nradd
nrp1a
nrp2a
nrp2b
ntd5
nxn
otx1
otx2a
p4ha1b
p4ha2
pacrg
pag1
pax2a
pcbp4
pcdh10a
pde4d
pdia4
pfkfb4b
pfkpb
phc2a
phldb1a
phtf1
pik3cb
UMAP_1
UMAP_2

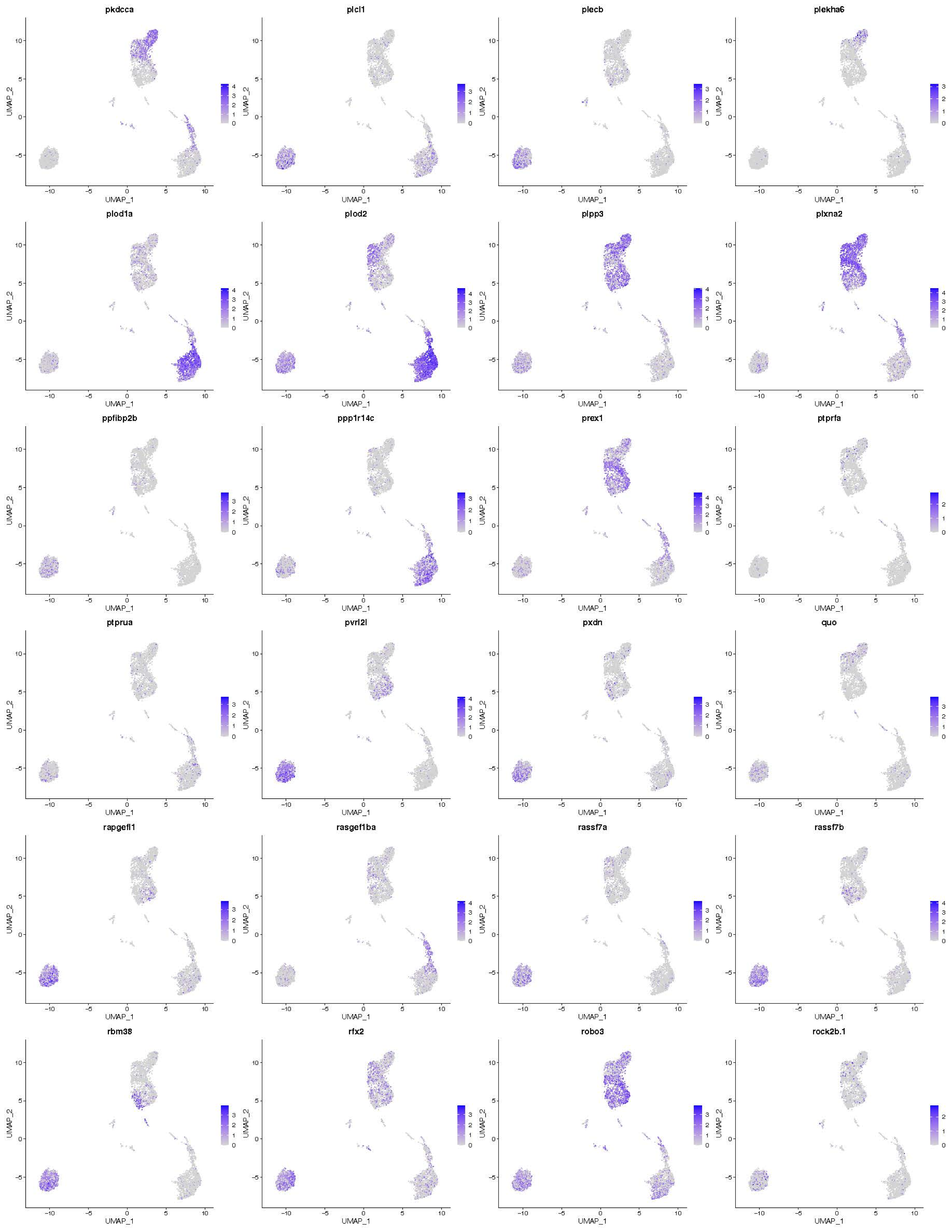
pkdcca
plcl1
plecb
plekha6
plod1a
plod2
plpp3
plxna2
ppfibp2b
ppp1r14c
prex1
ptprfa
ptprua
pvrl2l
pxdn
quo
rapgefl1
rasgef1ba
rassf7a
rassf7b
rbm38
rfx2
robo3
rock2b.1
UMAP_1
UMAP_2

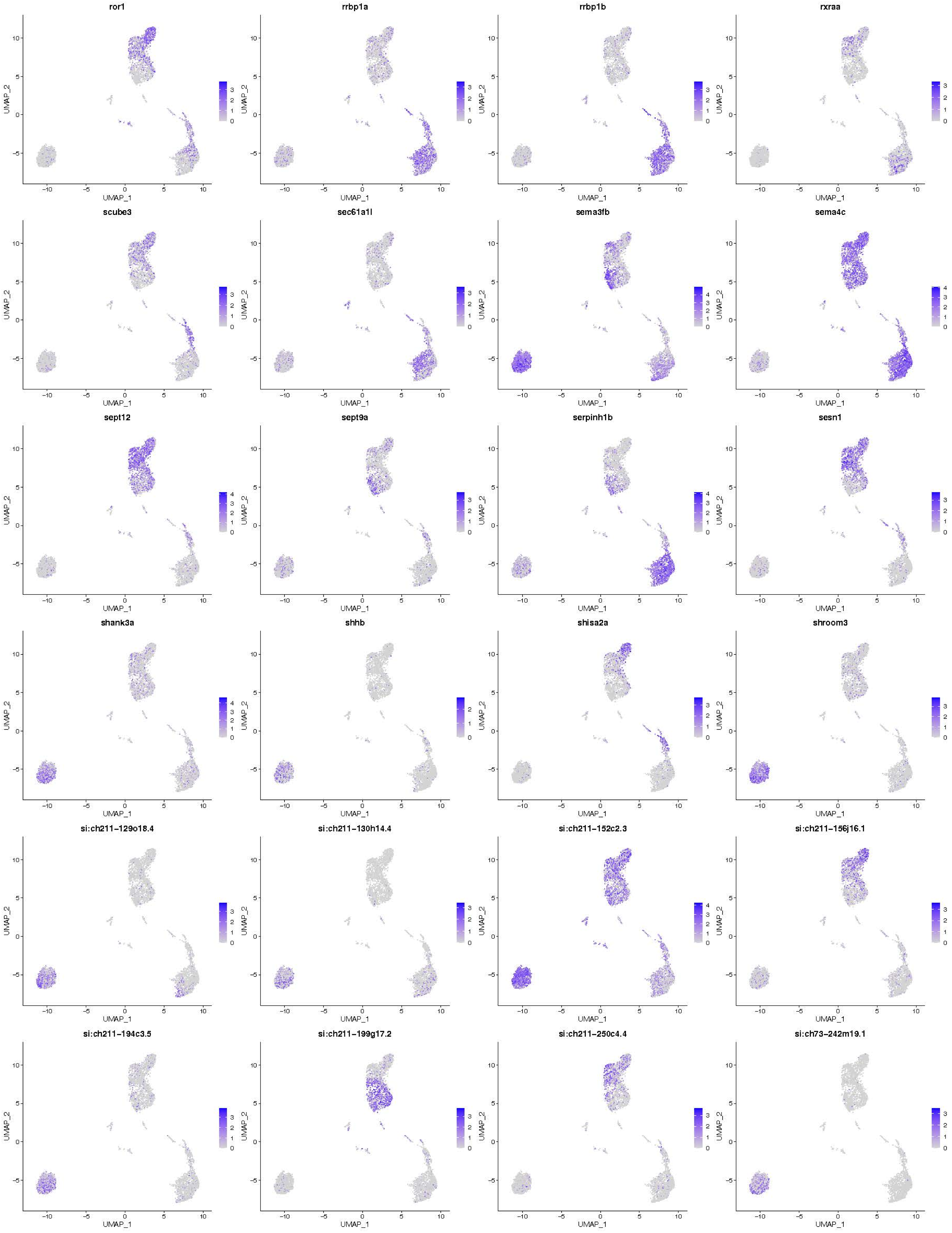

ror1
rrbp1a
rrbp1b
rxraa
scube3
sec61a1l
sema3fb
sema4c
sept12
sept9a
serpinh1b
sesn1
shank3a
shhb
shisa2a
shroom3
si:ch211-129o18.4
si:ch211-130h14.4
si:ch211-152c2.3
si:ch211-156j16.1
si:ch211-194c3.5
si:ch211-199g17.2
si:ch211-250c4.4
si:ch73-242m19.1
UMAP_1
UMAP_2

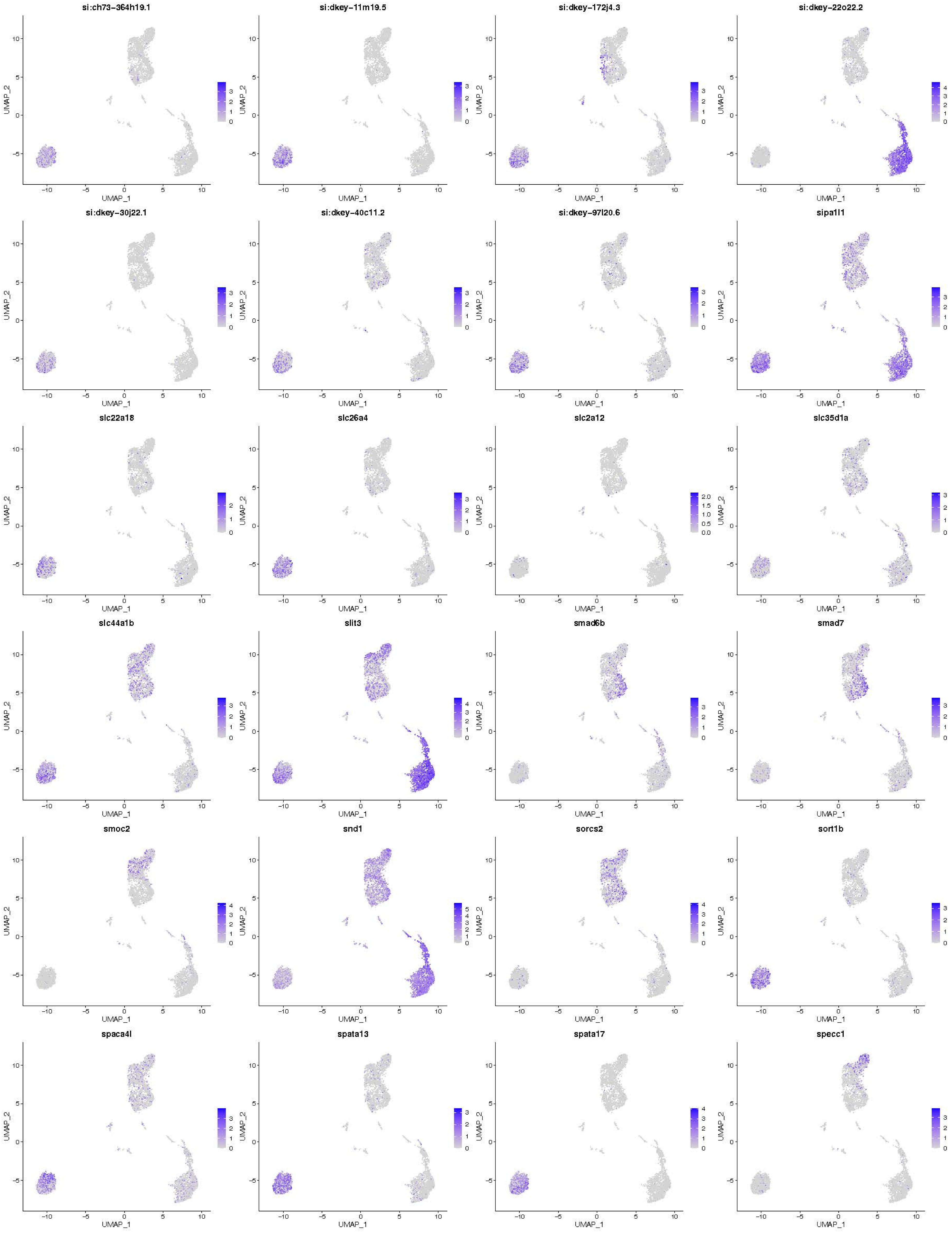

si:ch73-364h19.1
si:dkey-11m19.5
si:dkey-172j4.3
si:dkey-22o22.2
si:dkey-30j22.1
si:dkey-40c11.2
si:dkey-97l20.6
sipa1l1
slc22a18
slc26a4
slc2a12
slc35d1a
slc44a1b
slit3
smad6b
smad7
smoc2
snd1
sorcs2
sort1b
spaca4l
spata13
spata17
specc1
UMAP_1
UMAP_2

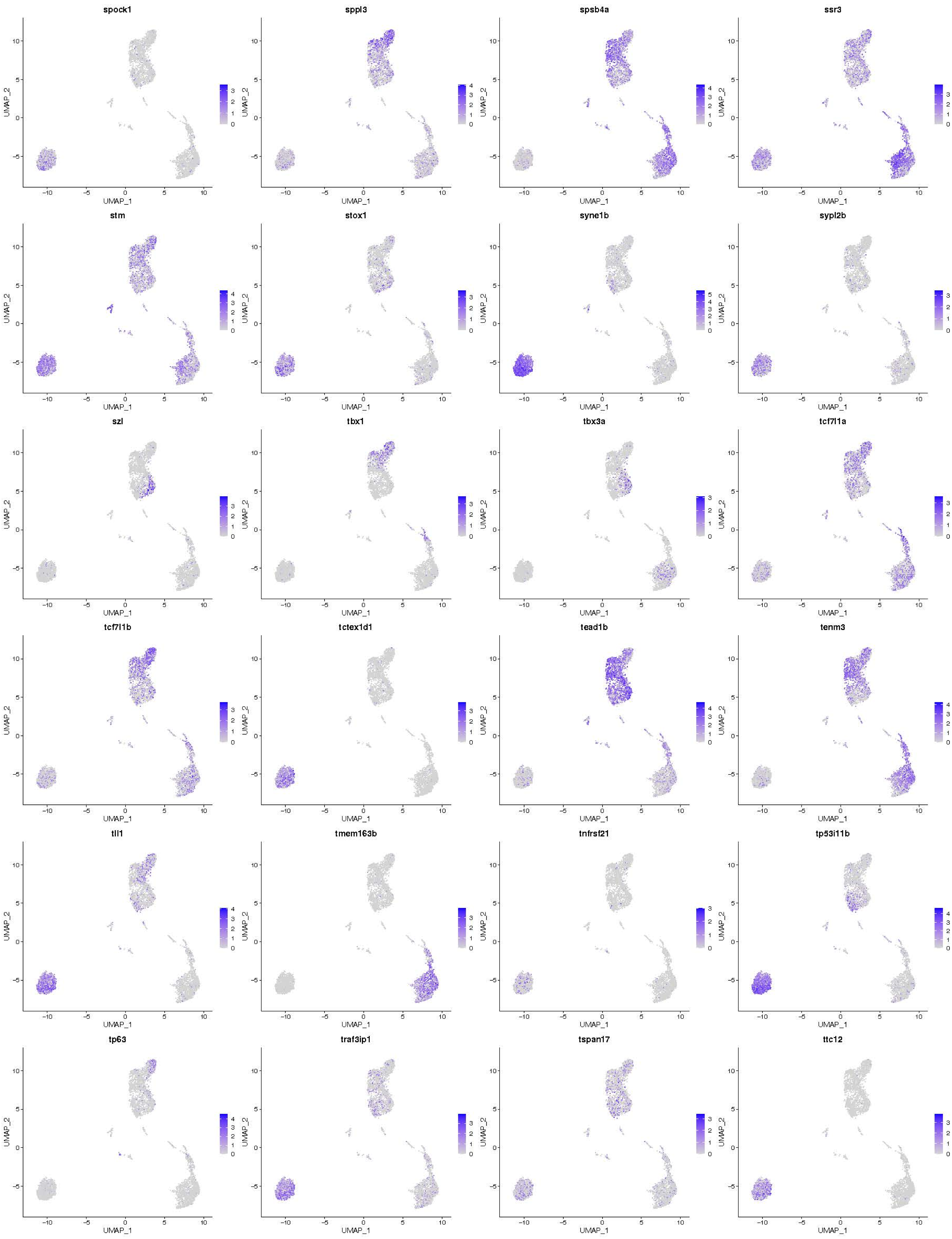

spock1
sppl3
spsb4a
ssr3
stm
stox1
syne1b
sypl2b
szl
tbx1
tbx3a
tcf7l1a
tcf7l1b
tctex1d1
tead1b
tenm3
tll1
tmem163b
tnfrsf21
tp53i11b
tp63
traf3ip1
tspan17
ttc12
UMAP_1
UMAP_2

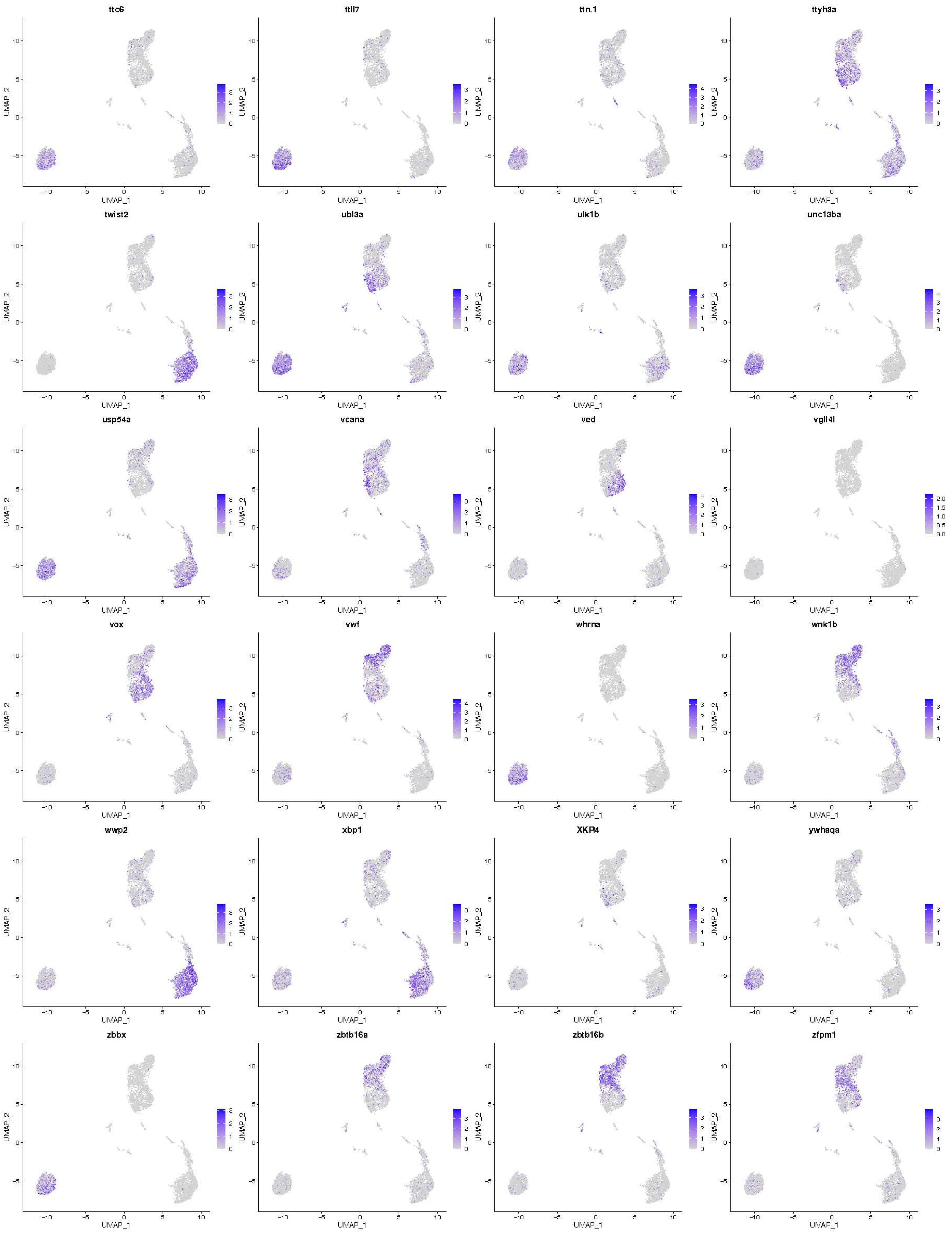
ttc6
ttll7
ttn.1
ttyh3a
twist2
ubl3a
ulk1b
unc13ba
usp54a
vcana
ved
vgll4l
vox
vwf
whrna
wnk1b
wwp2
xbp1
XKR4
ywhaqa
zbbx
zbtb16a
zbtb16b
zfpm1
UMAP_1
UMAP_2

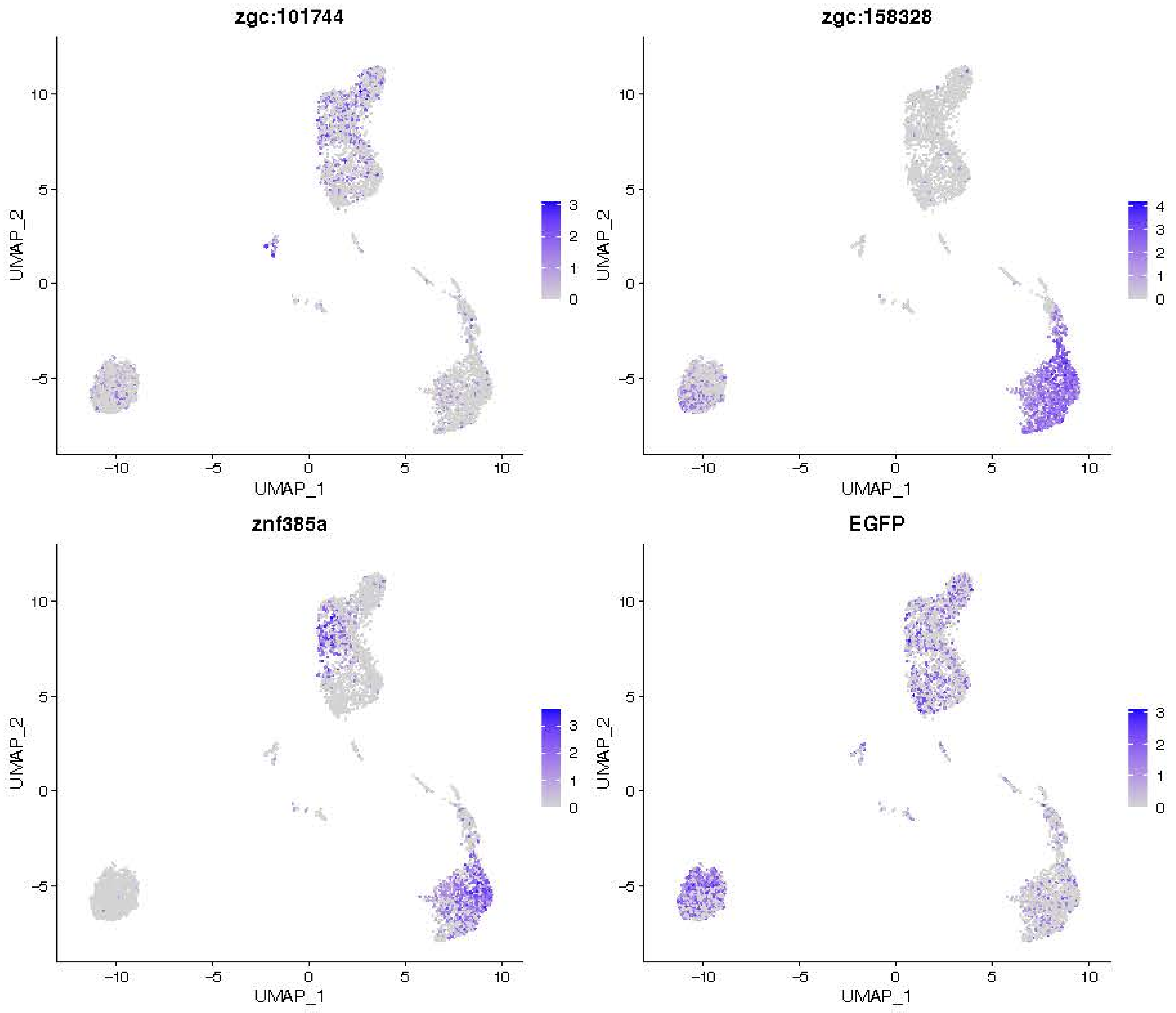
zgc:101744
zgc:158328
zgc:66433
zgc:92107
znf385a
EGFP
UMAP_1
UMAP_2

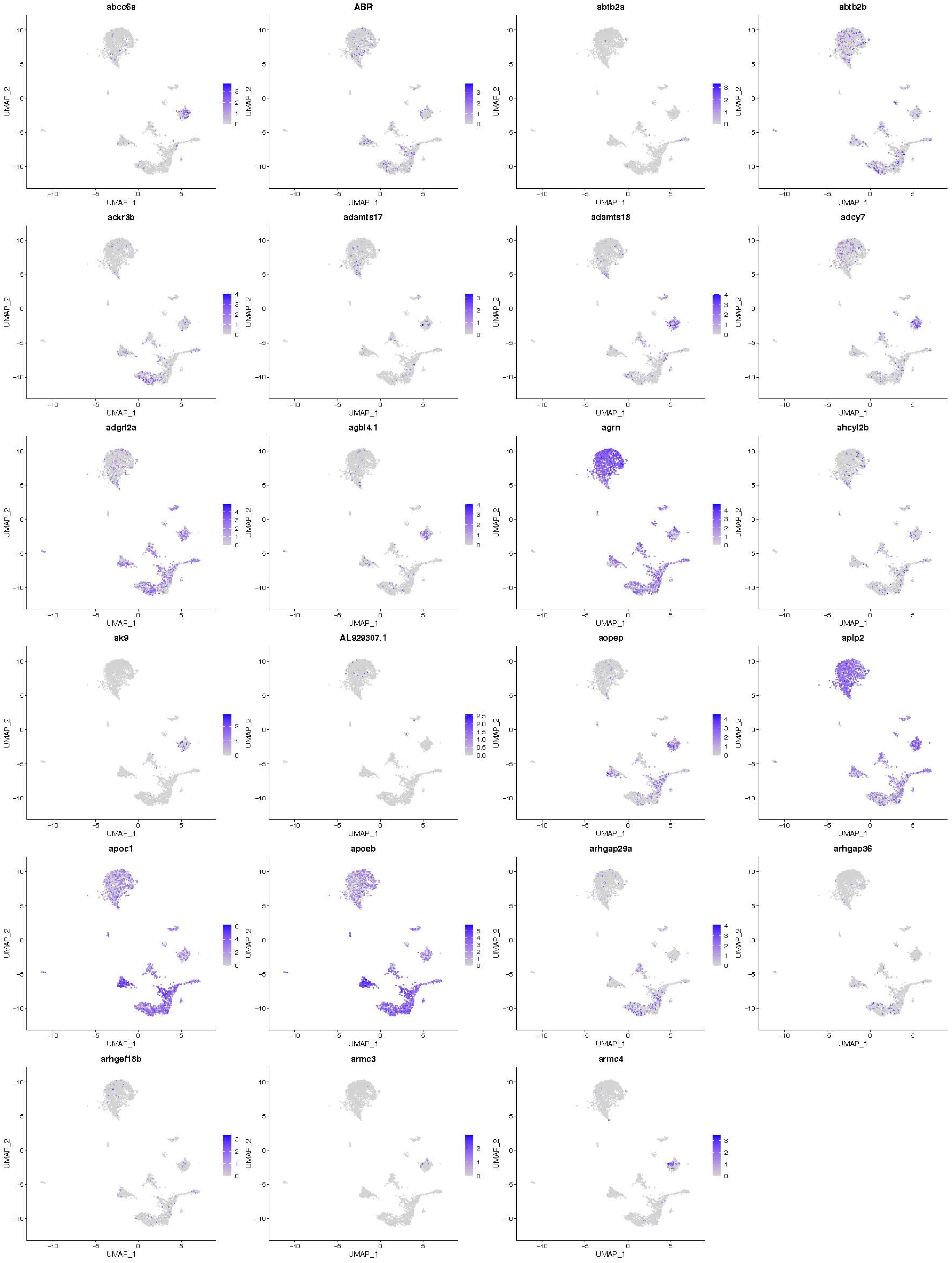

abcc6a
ABR
abtb2a
abtb2b
ackr3b
adamts17
adamts18
adcy7
adgrl2a
agbl4.1
agrn
ahcyl2b
ak9
AL929307.1
aopep
aplp2
apoc1
apoeb
arhgap29a
arhgap36
arhgef18b
armc3
armc4
UMAP_1
UMAP_2

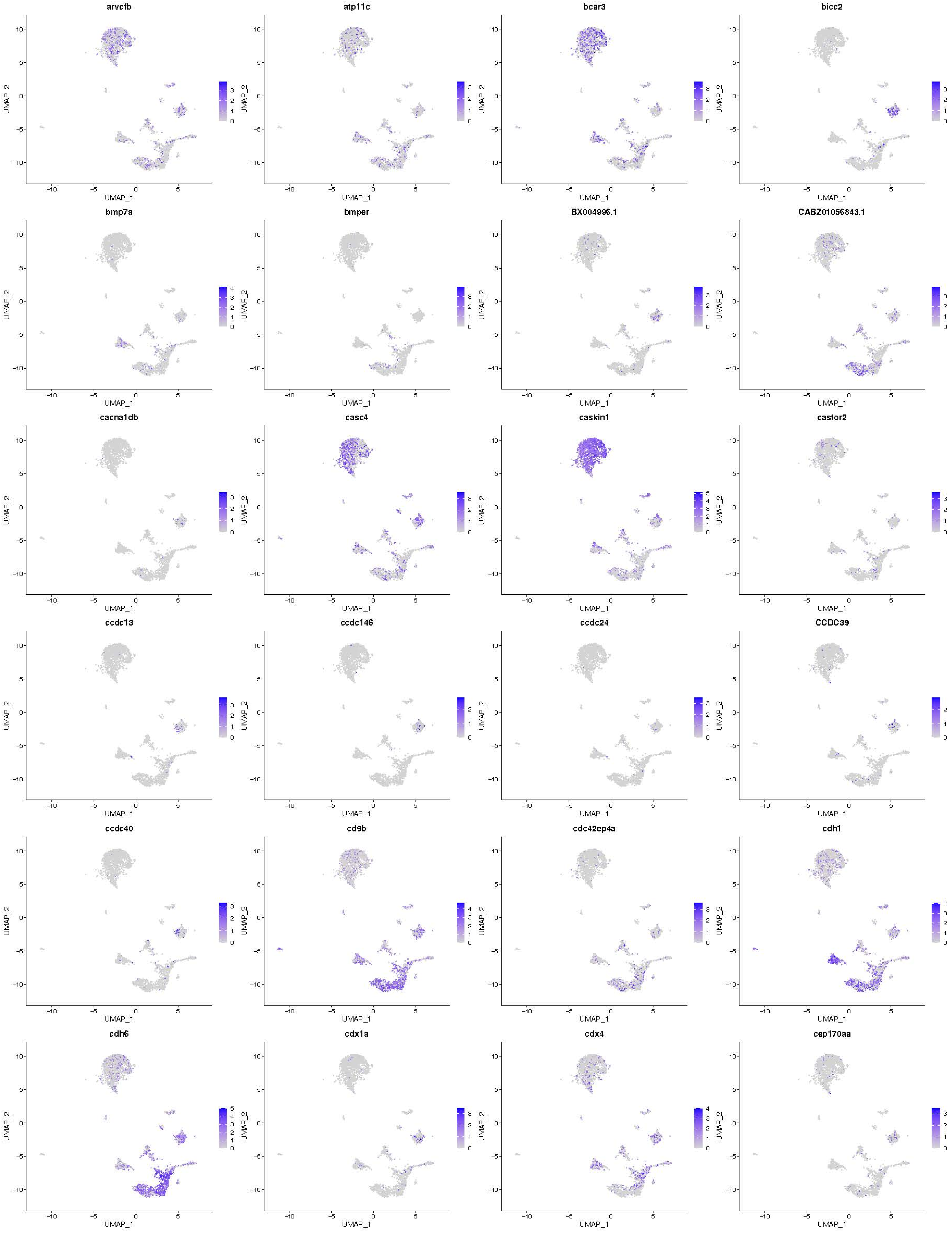
arvcfb
atp11c
bcar3
bicc2
bmp7a
bmper
BX004996.1
CABZ01056843.1
cacna1db
casc4
caskin1
castor2
ccdc13
ccdc146
ccdc24
CCDC39
ccdc40
cd9b
cdc42ep4a
cdh1
cdh6
cdx1a
cdx4
cep170aa
UMAP_1
UMAP_2

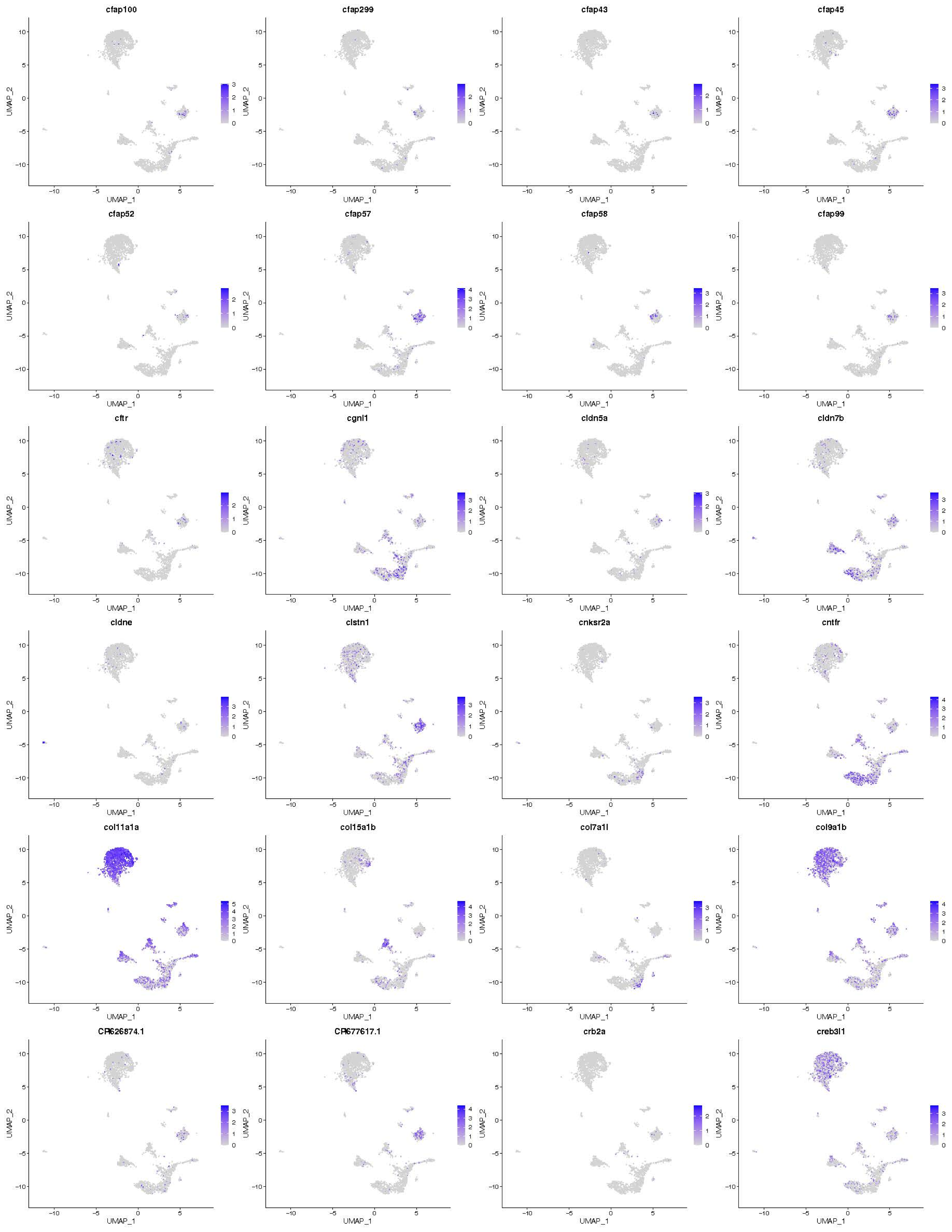

cfap100
cfap299
cfap43
cfap45
cfap52
cfap57
cfap58
cfap99
cftr
cgnl1
cldn5a
cldn7b
cldne
clstn1
cnksr2a
cntfr
col11a1a
col15a1b
col7a1l
col9a1b
CR626874.1
CR677617.1
crb2a
creb3l1
UMAP_1
UMAP_2

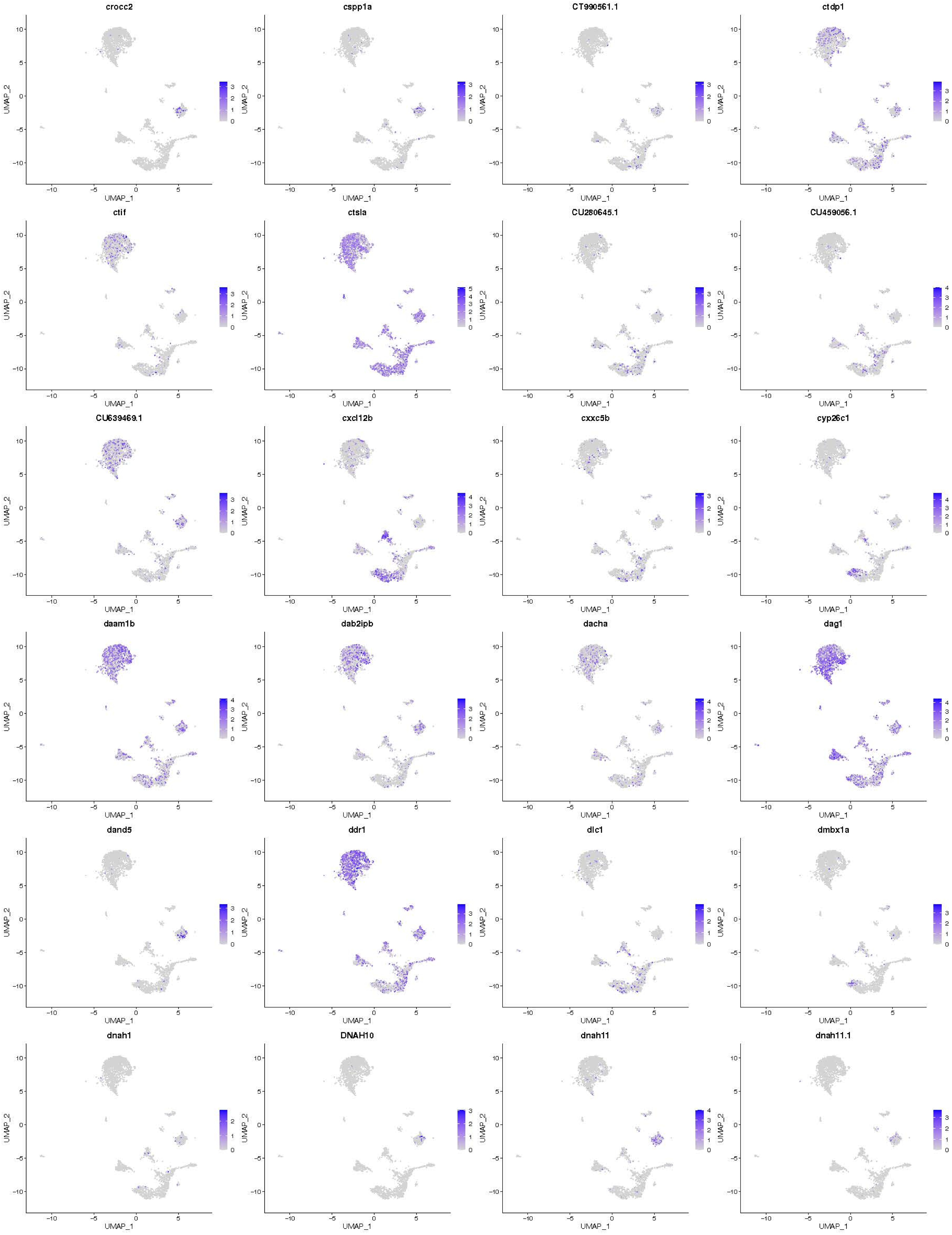
crocc2
cspp1a
CT990561.1
ctdp1
ctif
ctsla
CU280645.1
CU459056.1
CU639469.1
cxcl12b
cxxc5b
cyp26c1
daam1b
dab2ipb
dacha
dag1
dand5
ddr1
dlc1
dmbx1a
dnah1
DNAH10
dnah11
dnah11.1
UMAP_1
UMAP_2

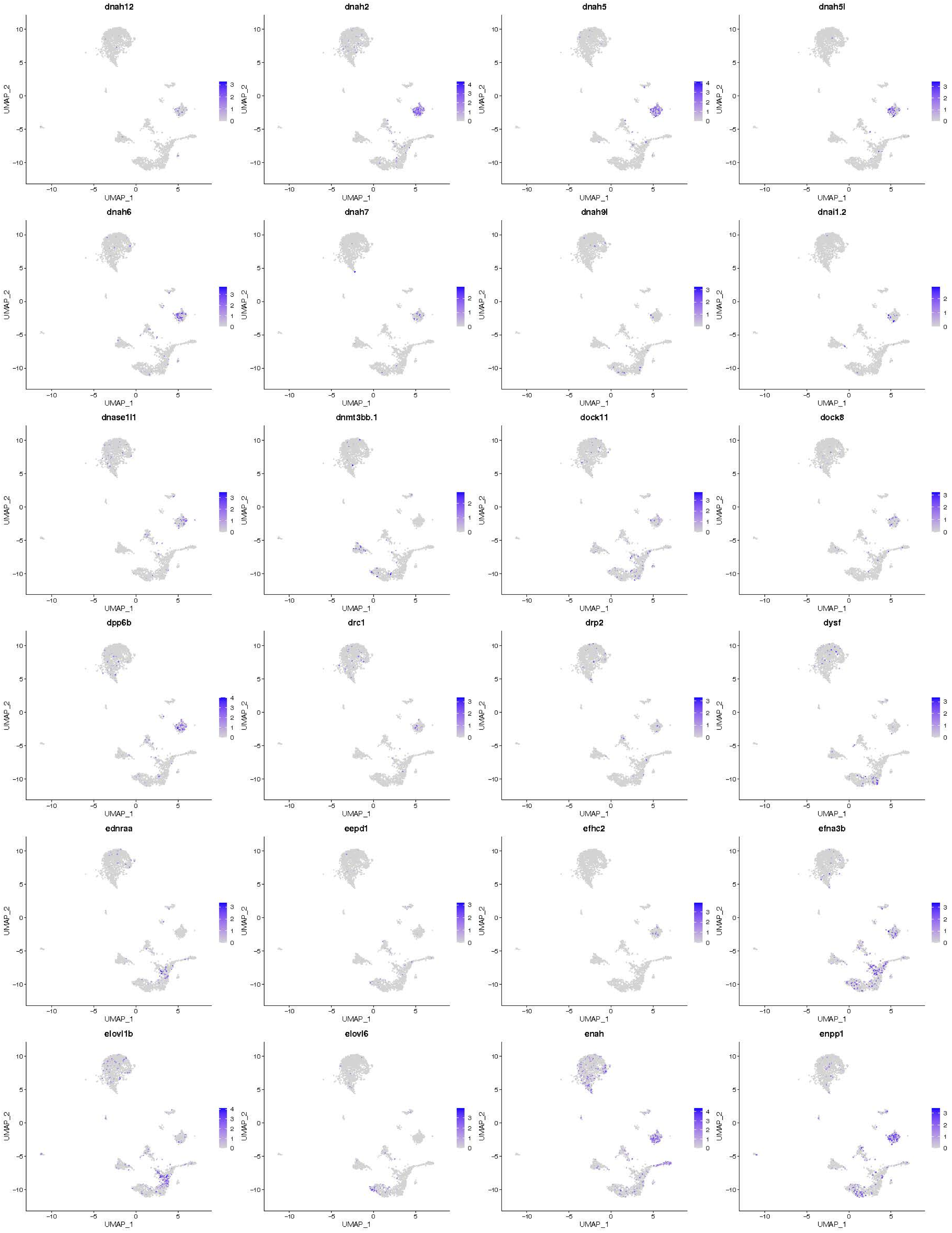
dnah12
dnah2
dnah5
dnah5l
dnah6
dnah7
dnah9l
dnai1.2
dnase1l1
dnmt3bb.1
dock11
dock8
dpp6b
drc1
drp2
dysf
ednraa
eepd1
efhc2
efna3b
elovl1b
elovl6
enah
enpp1
UMAP_1
UMAP_2

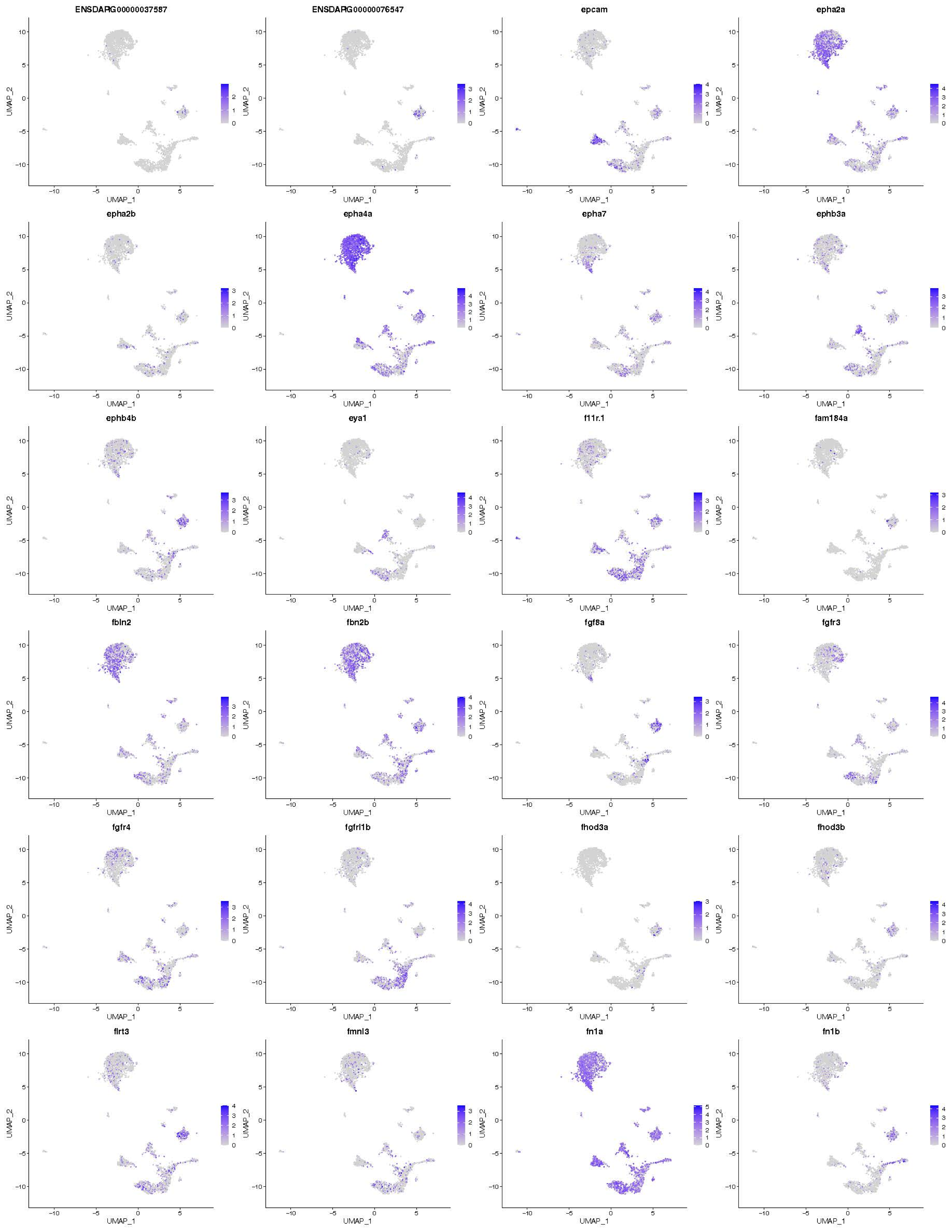

ENSDARG00000037587
ENSDARG00000076547
epcam
epha2a
epha2b
epha4a
epha7
ephb3a
ephb4b
eya1
f11r.1
fam184a
fbln2
fbn2b
fgf8a
fgfr3
fgfr4
fgfrl1b
fhod3a
fhod3b
flrt3
fmnl3
fn1a
fn1b
UMAP_1
UMAP_2

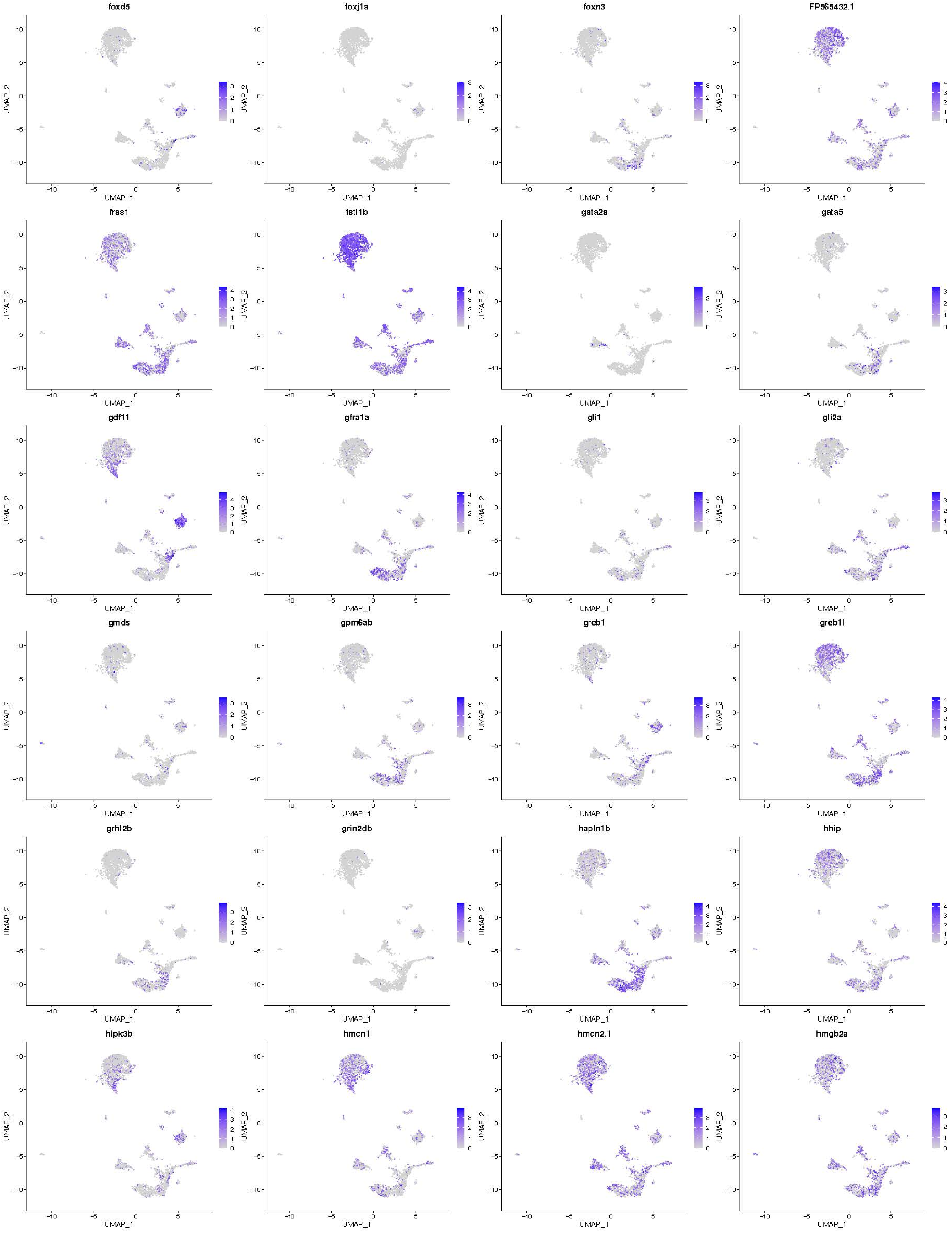
foxd5
foxj1a
foxn3
FP565432.1
fras1
fstl1b
gata2a
gata5
gdf11
gfra1a
gli1
gli2a
gmds
gpm6ab
greb1
greb1l
grhl2b
grin2db
hapln1b
hhip
hipk3b
hmcn1
hmcn2.1
hmgb2a
UMAP_1
UMAP_2

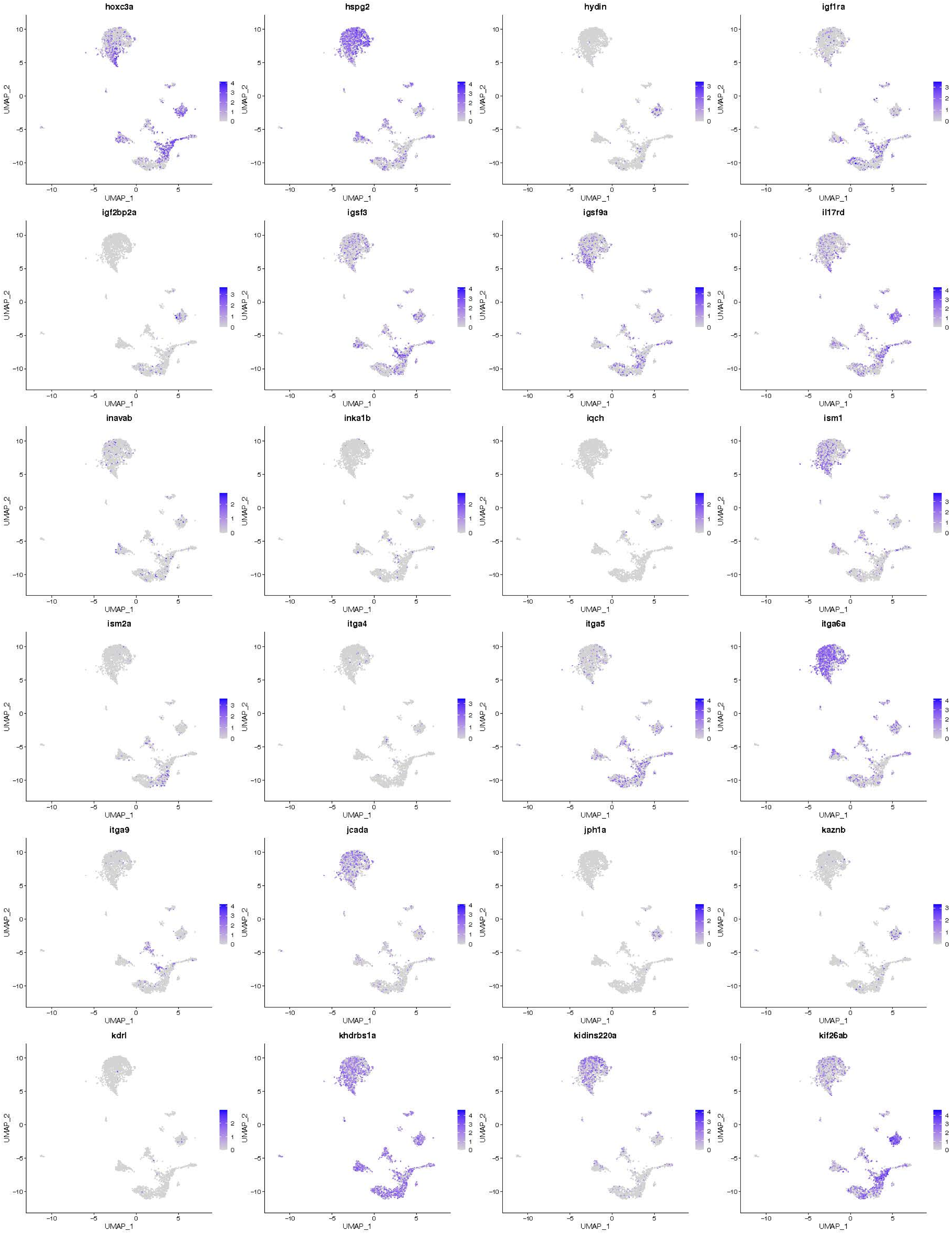

hoxc3a
hspg2
hydin
igf1ra
igf2bp2a
igsf3
igsf9a
il17rd
inavab
inka1b
iqch
ism1
ism2a
itga4
itga5
itga6a
itga9
jcada
jph1a
kaznb
kdrl
khdrbs1a
kidins220a
kif26ab
UMAP_1
UMAP_2

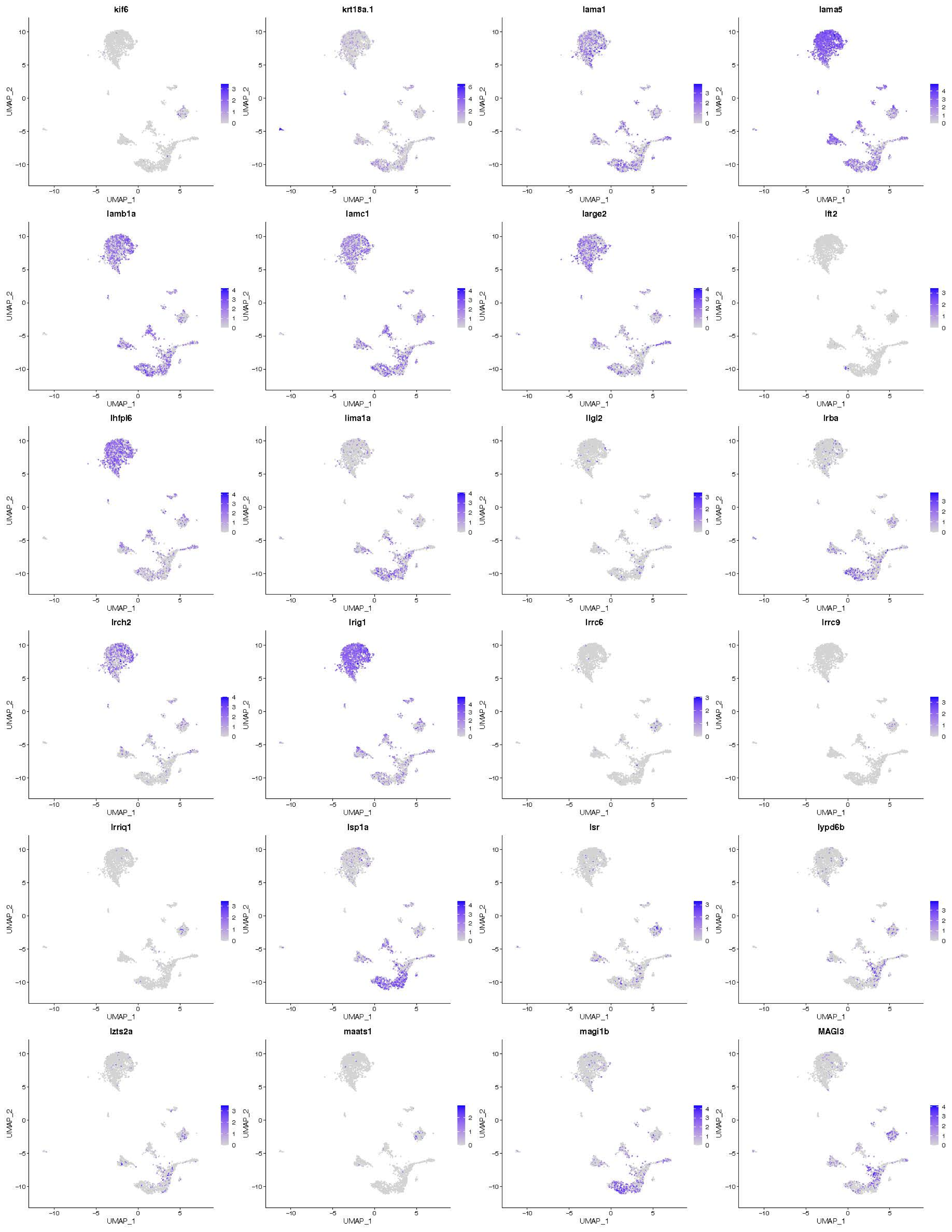

kif6
krt18a.1
lama1
lama5
lamb1a
lamc1
large2
lft2
lhfpl6
lima1a
llgl2
lrba
lrch2
lrig1
lrrc6
lrrc9
lrriq1
lsp1a
lsr
lypd6b
lzts2a
maats1
magi1b
MAGI3
UMAP_1
UMAP_2

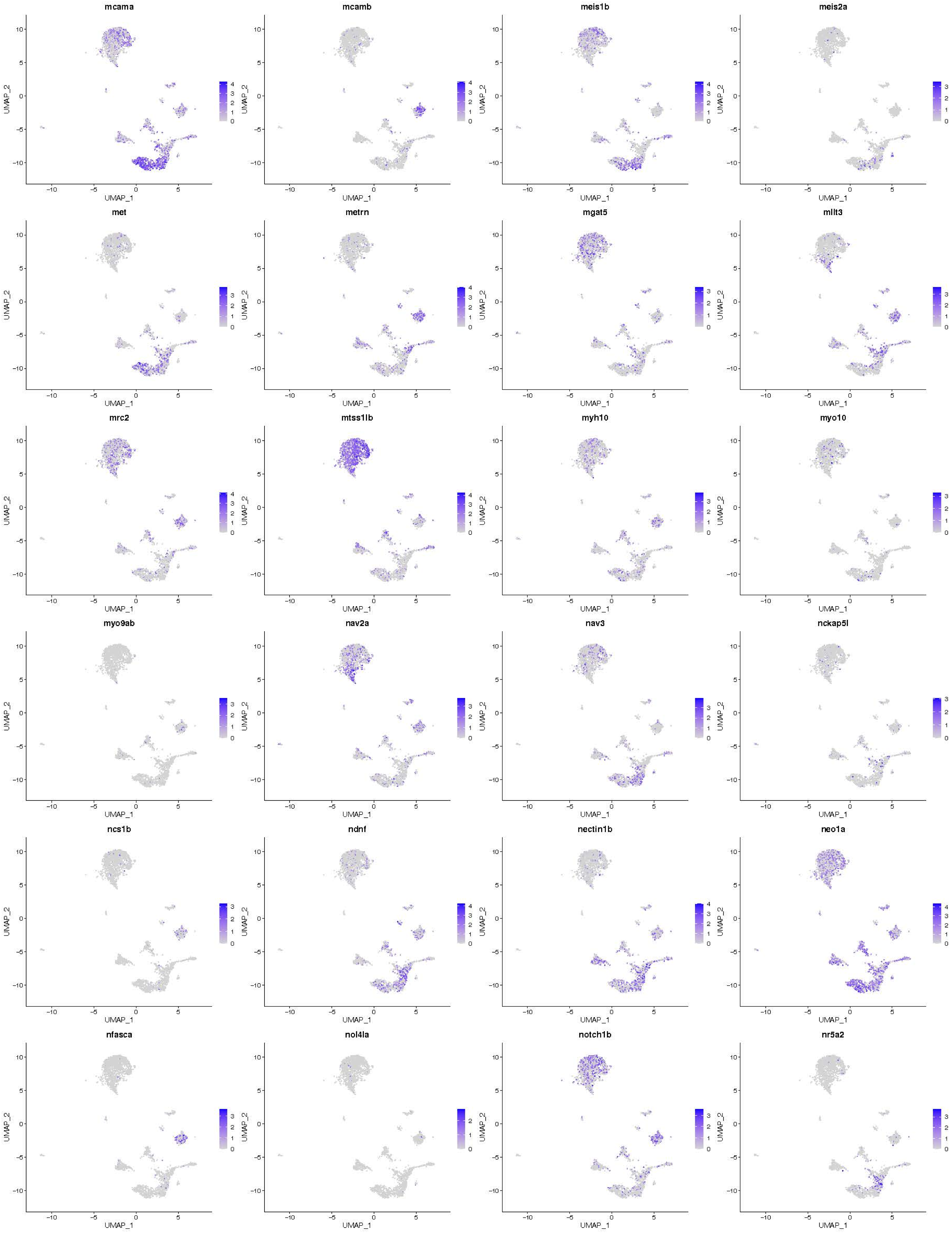
mcama
mcamb
meis1b
meis2a
met
metrn
mgat5
mllt3
mrc2
mtss1lb
myh10
myo10
myo9ab
nav2a
nav3
nckap5l
ncs1b
ndnf
nectin1b
neo1a
nfasca
nol4la
notch1b
nr5a2
UMAP_1
UMAP_2

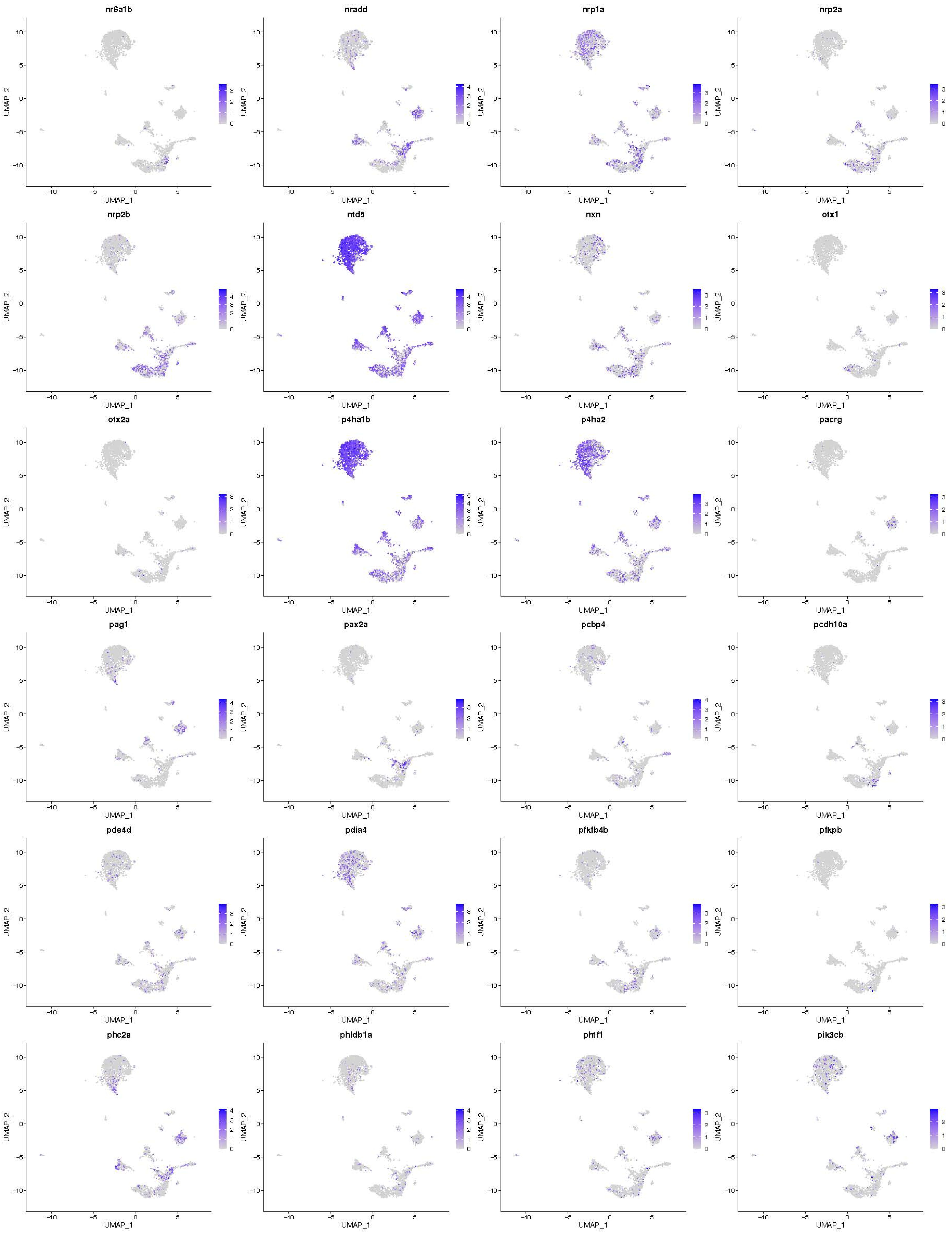
nr6a1b
nradd
nrp1a
nrp2a
nrp2b
ntd5
nxn
otx1
otx2a
p4ha1b
p4ha2
pacrg
pag1
pax2a
pcbp4
pcdh10a
pde4d
pdia4
pfkfb4b
pfkpb
phc2a
phldb1a
phtf1
pik3cb
UMAP_1
UMAP_2

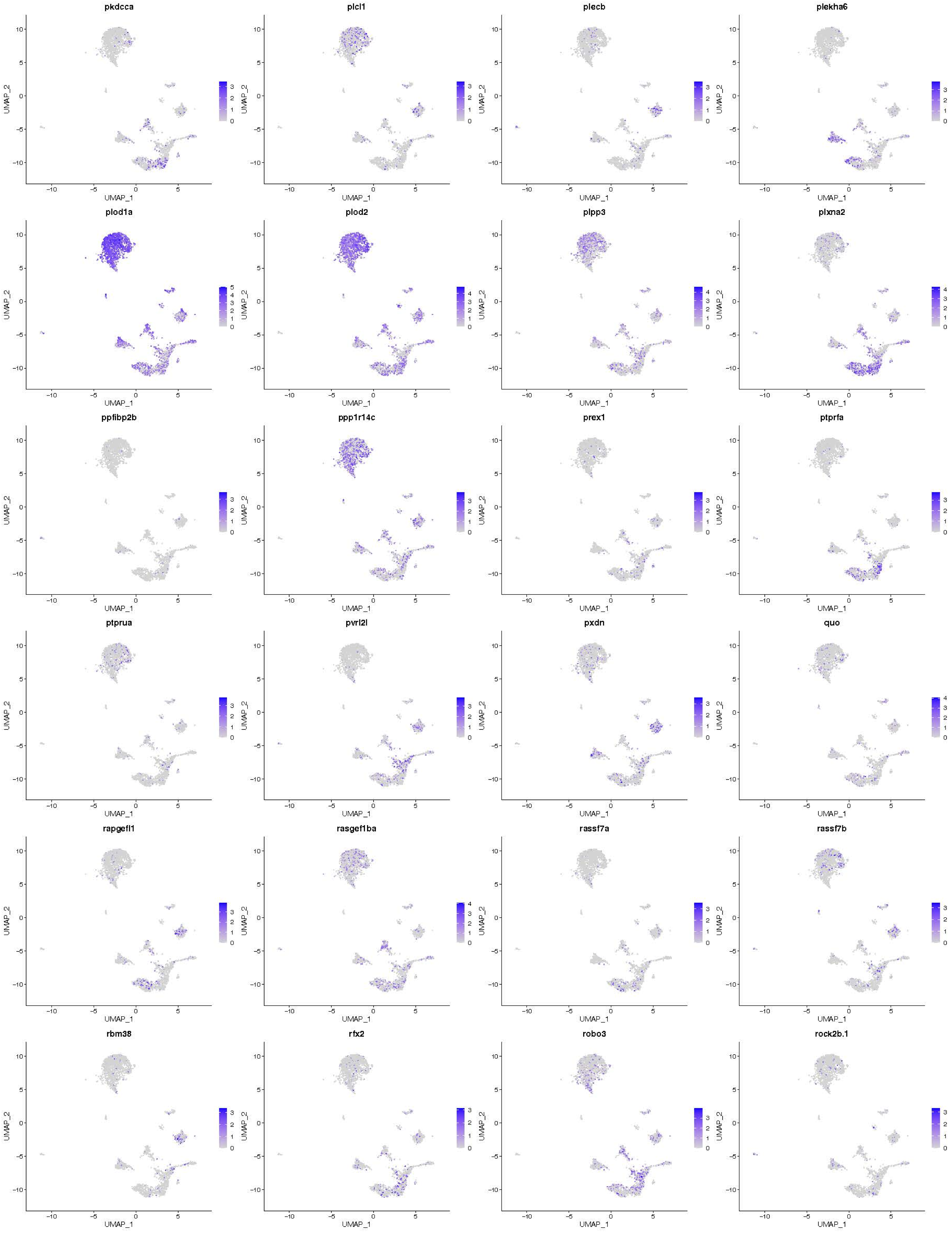
pkdcca
plcl1
plecb
plekha6
plod1a
plod2
plpp3
plxna2
ppfibp2b
ppp1r14c
prex1
ptprfa
ptprua
pvrl2l
pxdn
quo
rapgefl1
rasgef1ba
rassf7a
rassf7b
rbm38
rfx2
robo3
rock2b.1
UMAP_1
UMAP_2

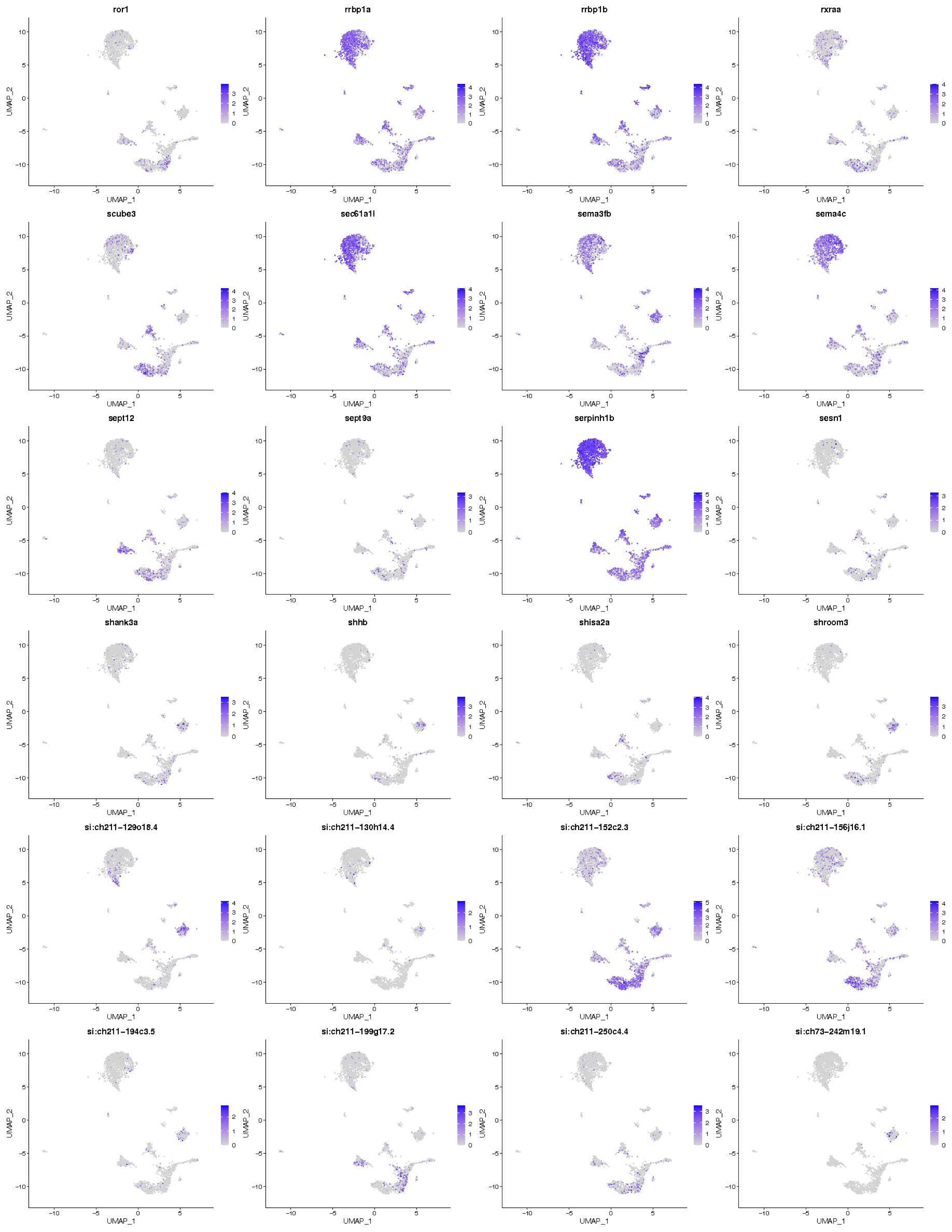
ror1
rrbp1a
rrbp1b
rxraa
scube3
sec61a1l
sema3fb
sema4c
sept12
sept9a
serpinh1b
sesn1
shank3a
shhb
shisa2a
shroom3
si:ch211-129o18.4
si:ch211-130h14.4
si:ch211-152c2.3
si:ch211-156j16.1
si:ch211-194c3.5
si:ch211-199g17.2
si:ch211-250c4.4
si:ch73-242m19.1
UMAP_1
UMAP_2

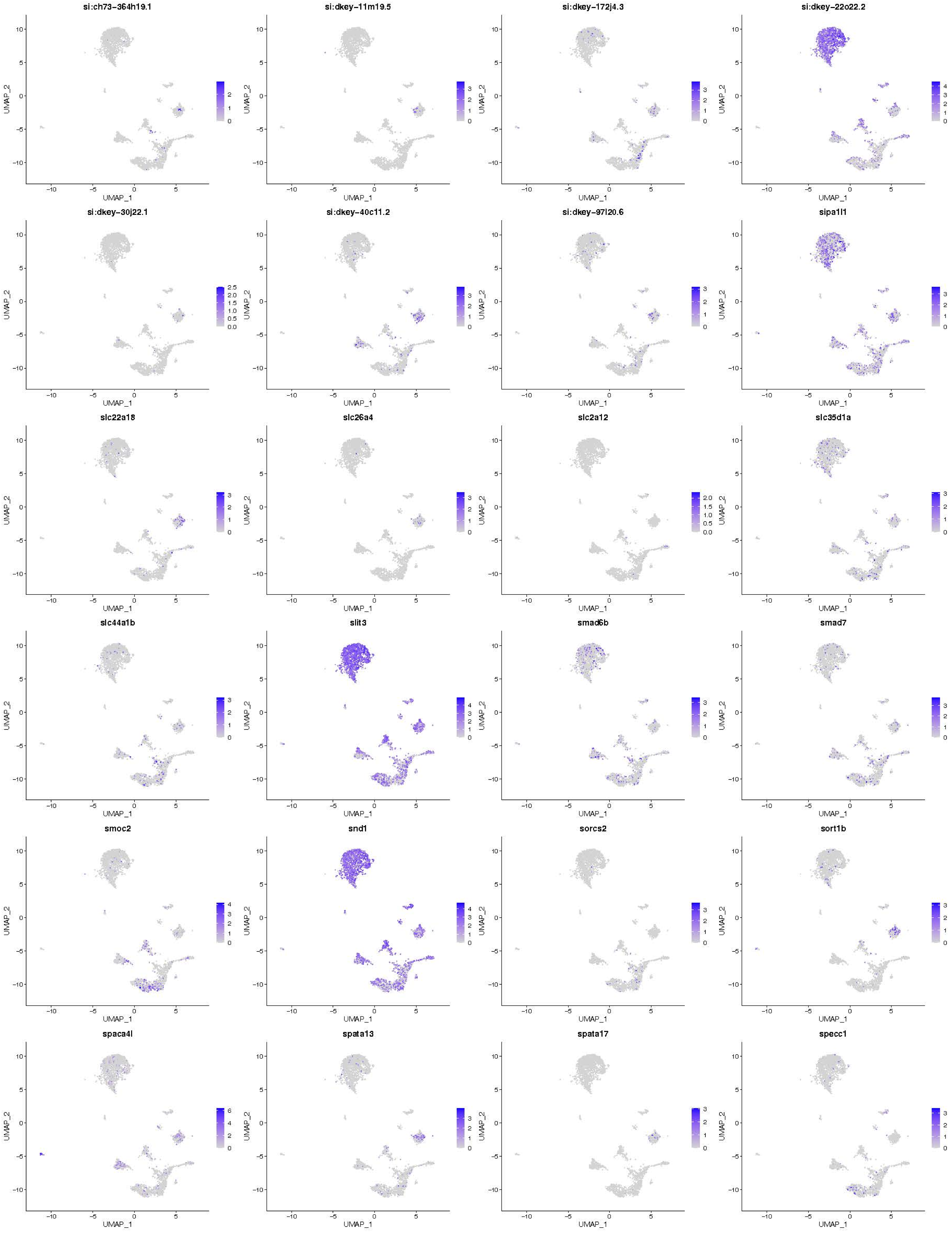
si:ch73-364h19.1
si:dkey-11m19.5
si:dkey-172j4.3
si:dkey-22o22.2
si:dkey-30j22.1
si:dkey-40c11.2
si:dkey-97l20.6
sipa1l1
slc22a18
slc26a4
slc2a12
slc35d1a
slc44a1b
slit3
smad6b
smad7
smoc2
snd1
sorcs2
sort1b
spaca4l
spata13
spata17
specc1
UMAP_1
UMAP_2

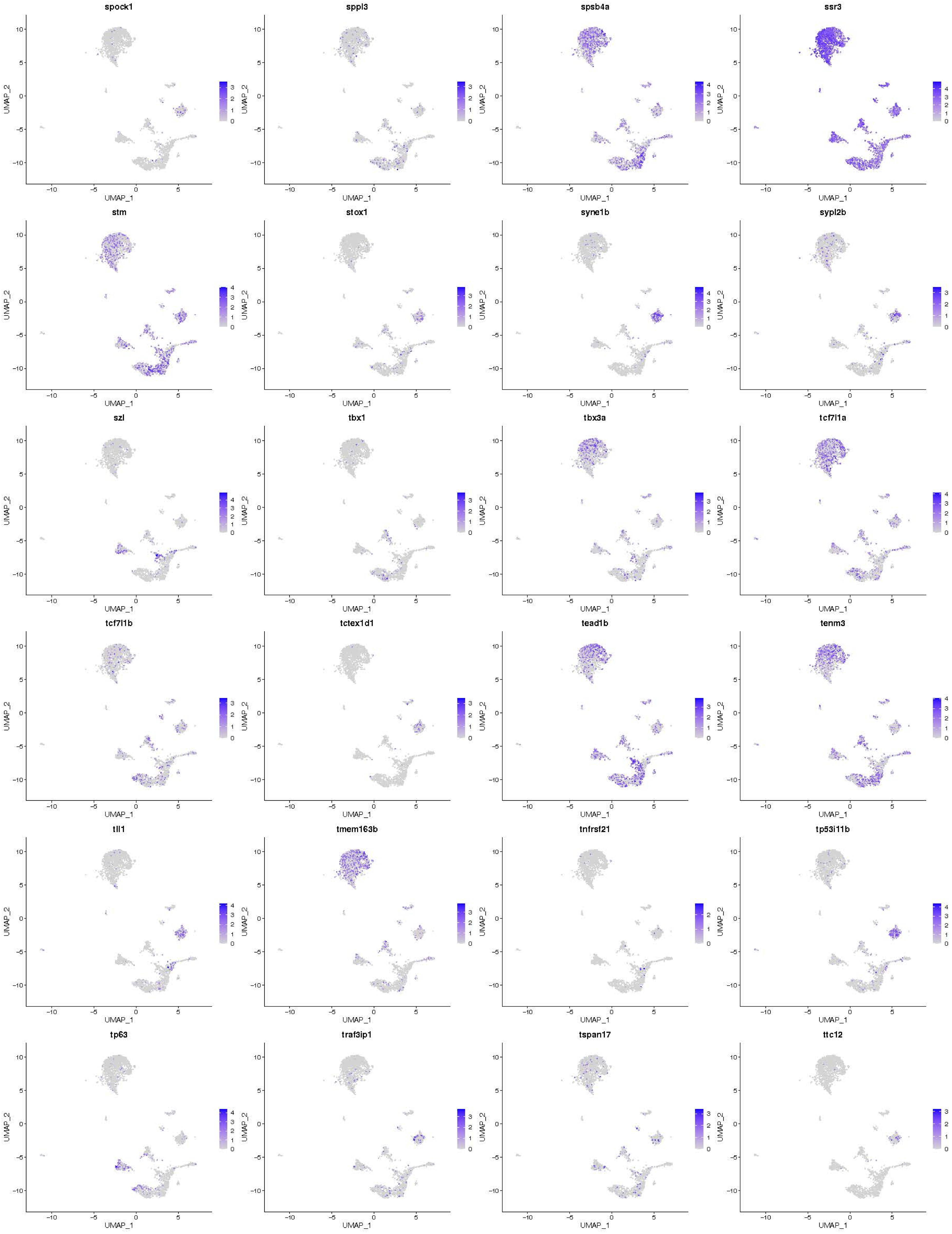
spock1
sppl3
spsb4a
ssr3
stm
stox1
syne1b
sypl2b
szl
tbx1
tbx3a
tcf7l1a
tcf7l1b
tctex1d1
tead1b
tenm3
tll1
tmem163b
tnfrsf21
tp53i11b
tp63
traf3ip1
tspan17
ttc12
UMAP_1
UMAP_2

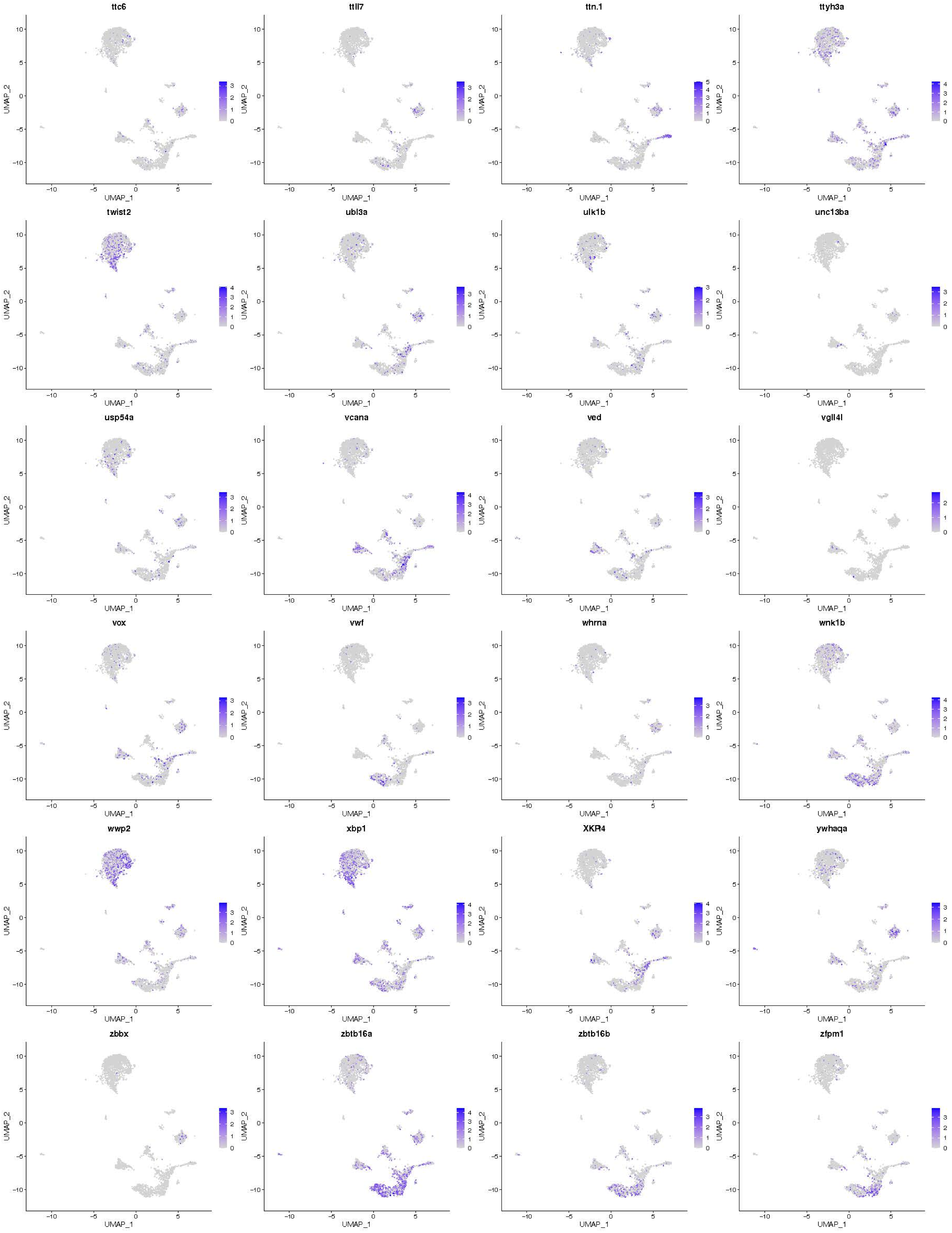
ttc6
ttll7
ttn.1
ttyh3a
twist2
ubl3a
ulk1b
unc13ba
usp54a
vcana
ved
vgll4l
vox
vwf
whrna
wnk1b
wwp2
xbp1
XKR4
ywhaqa
zbbx
zbtb16a
zbtb16b
zfpm1
UMAP_1
UMAP_2

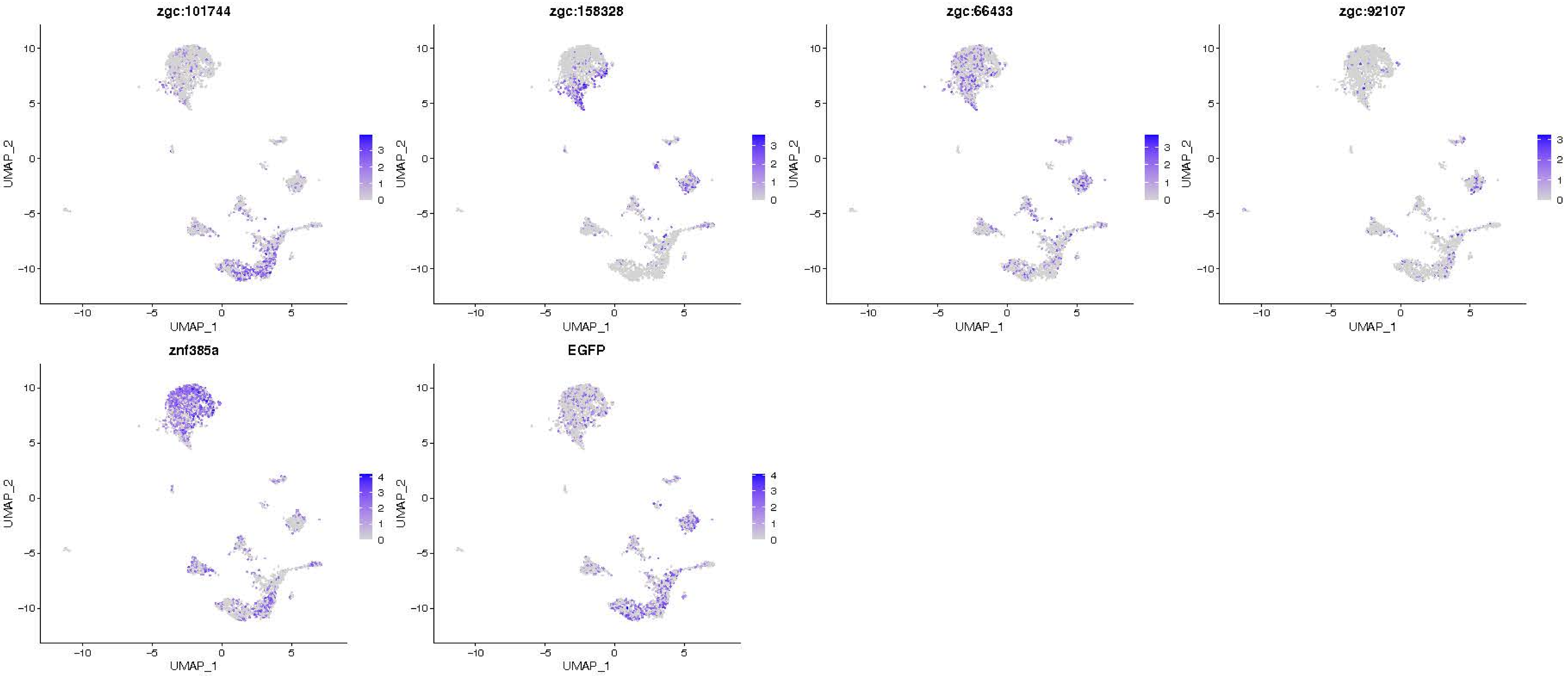
zgc:101744
zgc:158328
zgc:66433
zgc:92107
znf385a
EGFP
UMAP_1
UMAP_2

### Endoderm — 8 hpf

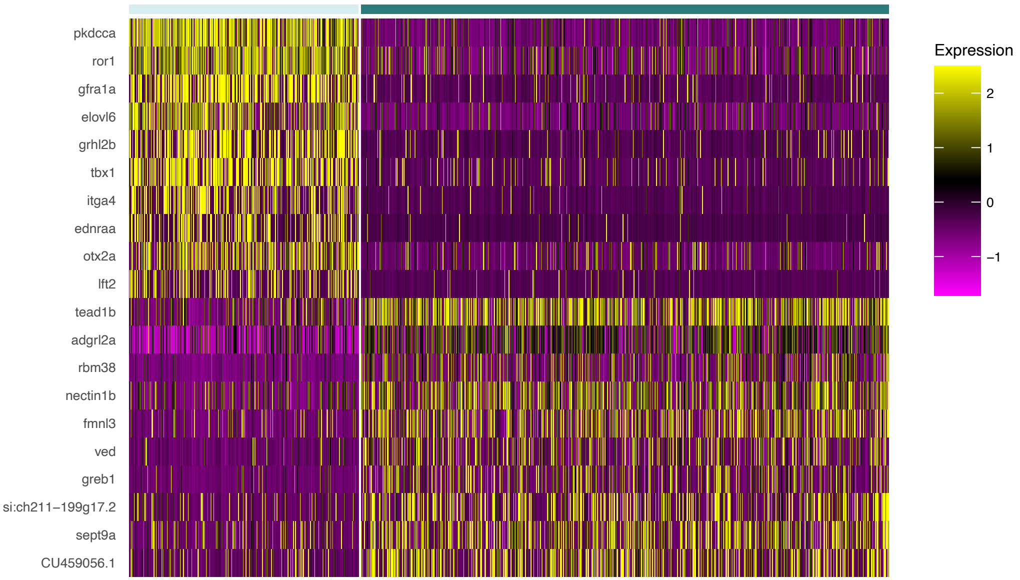

### Endoderm — 10 hpf

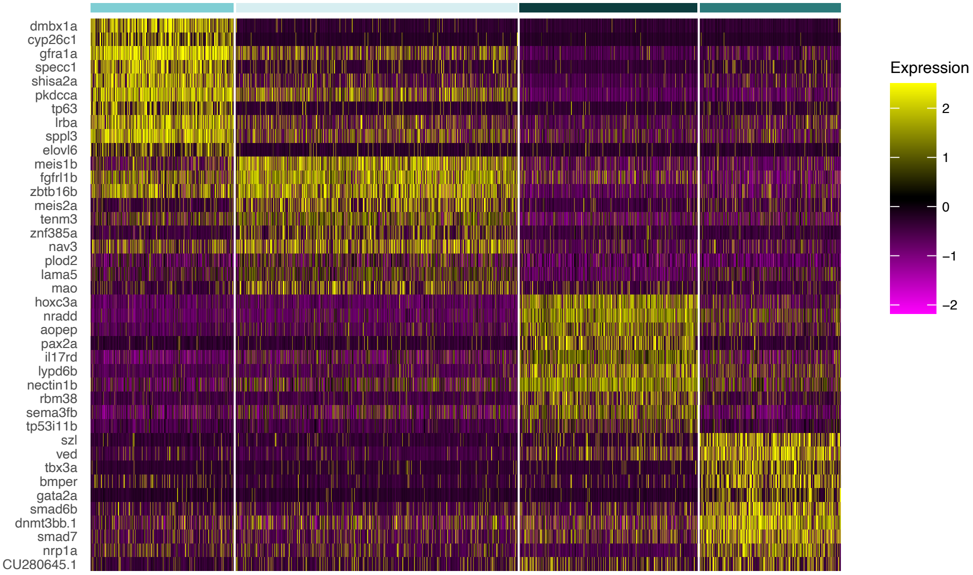

### Endoderm — 12 hpf

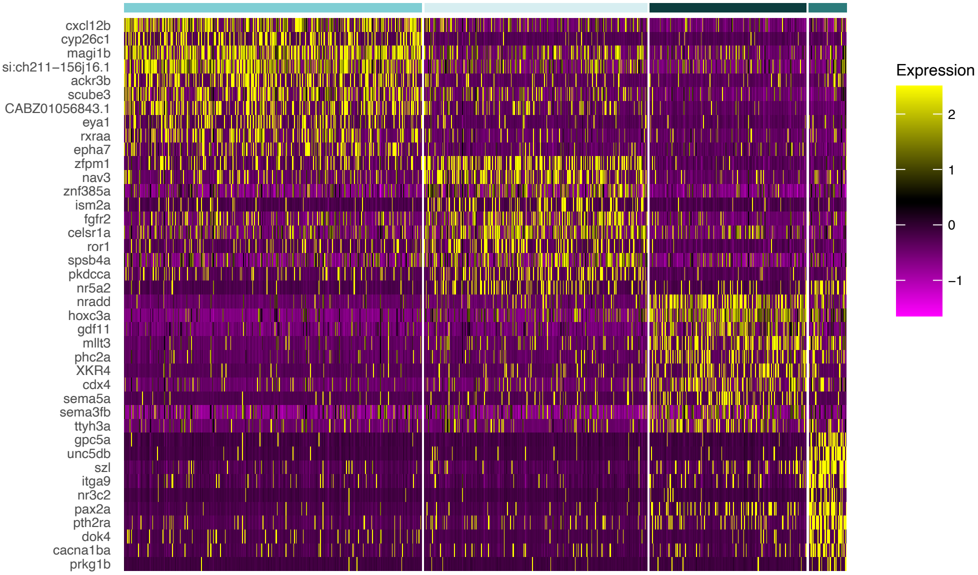
